# Effects of recombination on multi-drug resistance evolution in *Plasmodium falciparum* malaria

**DOI:** 10.1101/2024.12.18.628924

**Authors:** Kien Trung Tran, Tran Dang Nguyen, Daniel B Weissman, Eric Zhewen Li, Sachel Mok, Jennifer L Small-Saunders, Teun Bousema, Robert J Zupko, Thu Nguyen-Anh Tran, Maciej F Boni

## Abstract

When multiple beneficial alleles are present in a population but not linked together in any one individual, there is no general evolutionary result that determines whether recombination will speed up or slow down the emergence and evolution of genotypes carrying multiple beneficial alleles. Translated to infectious disease control, this evolutionary uncertainty means that when multiple types of drug resistance are present we do not know whether recombination will act more strongly to (1) bring together single-resistant genotypes into multi-drug resistant (MDR) genotypes, or (2) break apart MDR genotypes into single-resistant genotypes. In this paper, we introduce a new version of an established and validated individual-based malaria transmission model where we have added individual mosquito bites, interrupted feeding by mosquitoes, 25 drug-resistance related loci, and individual recombination events of different *Plasmodium falciparum* genotypes inside the mosquito. Recombination among *P. falciparum* genotypes in this model occurs from two sources of variation, multi-clonal infections and interrupted feeding by mosquitoes, and we show that 80% to 97% of MDR recombinant falciparum genotypes are projected to occur from single uninterrupted bites on hosts with multi-clonal infections (for malaria prevalence > 5%). Increases in the model’s interrupted feeding rate slowly increase the number of recombination events occurring from interrupted feeds. A comparison of drug-resistance management strategies with this new model shows that, over a 15-year timeframe, triple artemisinin-combination therapies (ACT) strategies show the largest reductions in treatment failures and the longest delays until artemisinin resistance reaches a critical 1% threshold. Multiple first-line therapies (MFT) are second best under these criteria, and ACT cycling approaches are third best. MFT strategies generate a greater diversity of recombinant genotypes but fewer recombination events generating MDR and slower emergence of these recombinant MDR genotypes.

## 1 Introduction

*Plasmodium falciparum* malaria caused an estimated 249 million symptomatic malaria cases and 608,000 deaths worldwide in 2022 [1]. The majority of the world’s malaria infections and deaths occur in Africa, in children under the age of five, despite the fact that highly efficacious artemisinin combination therapies (ACTs) have been available worldwide as first-line therapy for falciparum malaria for nearly twenty years. The emergence of artemisinin-resistant malaria parasites in East Africa in the middle of last decade has the potential to undermine our current approach to malaria control. As of mid-2024, five different genotypes – with mutations in the *pfkelch13* gene that is associated with slower parasite clearance by ACTs – are known to have spread successfully in Rwanda, Uganda, Tanzania, Kenya, and Ethiopia and reached local genotype frequencies above 0.20. Mathematical modeling analyses project that these genotypes frequencies will go up quickly over the next the decade [2,3].

As for all pathogens, the relationship between genotype and drug-resistance phenotype in *P. falciparum* is complex. The magnitude of resistance benefits varies across loci and drugs. Multi-allelic resistance types are known. And, pleiotropy occurs at several loci conferring opposing properties of drug resistance and drug sensitivity depending on the drug used. A minimum of twenty-five loci are known to have some effect on conferring drug resistance to artemisinin and the partner drugs used in ACTs, and epistatic effects are known to exist but difficult to measure because genotype information is typically not presented in therapeutic efficacy studies and randomized controlled trials of ACT efficacy. To add to this complexity, two types of transmission events are known to provide opportunities for recombination among falciparum genotypes: mosquito bites on individuals infected with multiple falciparum genotypes, and interrupted mosquito feeds where a blood meal is initiated on one host but completed on another. The influence of recombination will be stronger in high-transmission regions than in low-transmission regions, but it is not known how recombination affects the diversity and evolution of drug-resistant falciparum genotypes.

Understanding and correctly modeling this evolutionary complexity is critical for producing forecasts for *P. falciparum* evolution in Africa over the next decade. We have upgraded an individual-based simulation of falciparum malaria epidemiology and evolution [4] to add explicit chromosome structure, drug-resistance phenotype information for 25 loci, and individual (trackable) recombination events that can occur as a result of bites on multi-clonal hosts or interrupted feeds. We use the model to ask (1) which of the two recombination processes is more common, (2) which drug distribution strategies are best at delaying and slowing resistance in the context of these recombination processes [5–7], and (3) which strategies are generate more/fewer double-resistants and triple-resistant genotypes through recombination [8]. The development of a multi-locus model, covering all known loci associated with reduced efficacy of ACTs, is necessary to carry out practical evaluations of drug-resistance response policies in specific African countries that will be implementing them this decade.

## 2 Methods

The model used in this study builds upon the original individual-based model of *P. falciparum* evolution and epidemiology developed by Nguyen et al (2015, 2023) which has been utilized in various studies evaluating drug rotation strategies, deployment of multiple first-line therapies (MFT), and triple ACT strategies [4,5]. The previous version of the model had a genotype structure limited to 128 predefined genotypes modeling resistance markers in *P. falciparum* chromosomes 5, 7, 13, and 14. With the recent emergence of complex and diverse resistance patterns – especially in the *pfkelch13* and *pfcrt* genes – a more flexible model was necessary to address these challenges effectively. We introduced new features into this model (called model version 5.0) which include a chromosome-based genotype structure and a ‘representative mosquito cohort’ to keep track of the diversity of sampled genotypes and the effects of recombination on the evolution of drug resistance.

### 2.1 Chromosome-Based Locus Structure

In the previous model, a genotype was defined as a string with six characters representing six different loci that could mutate in any *Plasmodium falciparum* genome. These six loci were K76T in *pfcrt*, N86Y and Y184F in *pfmdr1*, C580Y in *pfkelch13*, and copy number variation (CNV) for *pfmdr1* and the *plasmepsin-2*,*3* genes on chromosome 14. For version 5, we extended the genotype structure to cover all 14 chromosomes of *Plasmodium falciparum* and introduced a new string format to represent genotype in the model. In this format, a wild-type genotype is denoted by:

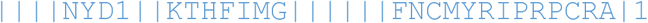

where the thirteen vertical bars denote breaks between chromosomes, and each character represents an allele or a copy number in a specific gene. This string serves as a key in a key-value pairing (i.e. a ‘map’ or ‘dictionary’ associative array structure) to obviate the memory needs of permanently storing information on millions of genotypes. Entries are added and removed from this map as genotypes appear or disappear in the simulation. Full details of this new genotype structure and the number of alleles integrated into the new model are provided in Supplement A Section 2.

A masking structure was built in to the simulation to allow for mutation at only some loci. This was done to allow the simulation to keep genetic variation restricted to what is observed in the field. For example, a maximum of one *pfkelch13* mutations is allowed in all simulations here since double mutants have not been observed yet. In the results presented in this paper, only 10 mutation types or CNVs are enabled: two loci and copy number variation in *pfmdr1*, five loci in *pfcrt*, a single amino-acid change in *pfkelch13*, and copy number variation in the plasmepsin genes.

### 2.2 Assumption of Non-Epistasis Among Loci

Epistatic interactions among genes – where the effect of a pair or a group of mutations is different than the sum of their individual effects – are both critical and difficult to measure when attempting to model and forecast evolutionary trajectories. In the new version of the model, we begin with an assumption of non-epistasis among loci (meaning that the phenotypic effects of mutations are independent of one another and follow a simple multiplicative fitness model) and we add in epistatic effects for pairs of loci for which there is sufficient supporting evidence. This is done in the pharmacodynamic component of the model, where the shift in the EC50 factor (drug concentration at which parasite killing is 50% of maximum) is assigned in a multiplicative fashion when multiple mutations accumulate that increase a parasite’s resistance level to a particular drug (see Supplement A Section 13). In doing this, we reduce the time and space complexity of computing drug efficacy on 32 million potential genotypes in the model. For mutational pairs where epistatic effects are known or have been inferred (as in Supplement 2 of [4]) the inferred fitness value (with epistasis) overrides the multiplicative fitness value calculated under an assumption of no epistasis.

### 2.3 Post-Recombination Mosquito Cohort (PRMC)

The new model includes a representative mosquito population but does not model all individual mosquitoes in a population. The representative mosquito population is modeled via a cohort of 100 to 500 individual mosquitoes that carry parasites in the post-recombination phase (ookinetes, oocysts, sporozoites) of parasite development in the mosquito. This mosquito cohort carries representative genotypes from the population of human hosts in the model. Mosquitoes in the cohort are populated with falciparum genotypes every day via biting and parasite sampling events that include interrupted feeds through which the parasites can sample parasites from two different hosts. The interrupted feeding rate (IFR) is defined as the proportion of completed mosquito feeds that occurred on two or more (not necessarily infected) humans. Free recombination among falciparum’s 14 chromosomes occurs after mosquito sampling, and a single genotype is chosen through this random process to be assigned to the sporozoites population that will infect human hosts 11 days later. As a result, the mosquito cohort acts as a pool for introducing novel recombinant genotypes into the human population. The mosquito cohort has two configurable factors: cohort size and the interrupted feeding rate. Interrupted feeding rate is set to between 0% and 20%; this is the percentage of fed mosquitoes that have blood from two or more humans, regardless of human infection status. A cohort size of 100 means that 1100 mosquitoes are tracked at any time since 11 days of mosquito history must be tracked. Details of this transmission mechanism and mosquito cohort design are described in Supplement A Section 8.

### 2.4 Counting Recombinations in Interrupted Feeds and from Multiclonal Infections

To determine what percentage of recombination events occur due to interrupted feeds, we counted recombination events that produce double-resistant genotypes from single-resistant genotypes. A base scenario was run where recombination among several clones circulating in a single host was not permitted in a simulation (this is a “within-host recombination turned off” mode of the model); in this scenario, all recombination occurs as a result of interrupted feeds. Every set of base scenario runs was compared to a set of ‘normal runs’ where the within-host recombination is “turned on” in the model (as it is in real life). This approach was taken to confirm that implementation of the interrupted-feeding mechanism in the code was done correctly. After completing the analysis, it was apparent that counting different types of recombination events could be done either by turning within-host recombination “on” or “off”, or equivalently by comparing results between IFR=0.0 and IFR=0.2 scenarios.

### 2.5 Scenario Calibration

To calibrate the model, we adjusted the mutation probability so that the 580Y allele frequency in the model would reach 0.01 after 7.0 years in a population of 100,000 individuals, with a *P. falciparum* prevalence of 10% and with 40% of malaria cases receiving dihydroartemisinin-piperaquine (DHA-PPQ) as first-line treatment. Following this calibration, simulations were configured with a population size of 100,000, and *P. falciparum* prevalence rates set at 5% and 25% for medium and high prevalence scenarios, respectively. Model simulations are run for 10 years as burn-in with 50% of the population treated with a generic 80% efficacy drug and 50% of malaria cases untreated; no private-market drugs are used during this period. The transmission parameter is calibrated so that malaria prevalence in 2-10 years olds (abbreviated PfPR here) reaches either 5% or 25% at the end of the burn-in period. This is intended to stabilize the simulation’s dynamics to an African epidemiological scenario in 2005 just prior to ACT deployment. Initial genotypes in simulation runs were wild-type alleles for *pfkelch13, pfmdr1*, and *pfpm2*,*3*, except for *pfcrt* 76T and a 50/50 ratio of *pfmdr1* N86 and 86Y. Genotypes with the N86 allele are selected for during burn-in due to the model’s built in cost of [9,10] resistance (*c*_*R*_). Parasites carrying drug-resistance alleles carry a fitness cost *c*_*R*_ (relative to wild-type parasites) in absence of drugs; see supplement A section 6). At the end of burn-in, the genotype balance between 76T-N86-Y184 (“TNY”) and 76T-86Y-Y184 (“TYY”) genotypes is typically around 50% - 55% TNY when *c*_*R*_ = 0.0005 and 70% - 90% TNY genotypes when *c*_*R*_ = 0.005.

### 2.6 Treatment strategies

After ten years of burn-in, different drug-resistance mitigation strategies are put into place and the model is run for an additional fifteen years. An assumption of 75% treatment coverage is made. A 15-year range was chosen so that a fair comparison could be carried out between MFT with three ACTs and a 5-year cycling policy that deployed all three ACTs for exactly five years each. During these 15 years, the outcome measures collected are (a) number of treatment failures (TF), (b) the time it took for artemisinin-resistant alleles to reach an 0.01 genotype frequency, (c) the number of recombination events that generated double-resistant genotypes from single-resistant genotypes, and (d) the number of recombination events that generated triple-resistant genotypes from single-resistant and double-resistant genotypes. It was assumed that artemether-lumefantrine (AL), artesunate-amodiaquine (ASAQ), and DHA-PPQ were available as possible first-line therapy for antimalarial treatment.

Four common and previously analyzed [4,5] treatment strategies were evaluated: (1) multiple first-line therapies (MFT) where all three ACTs are distributed simultaneously in equal amounts with 33.3% of malaria cases treated by each ACT; (2) 5-year cycling where first-line therapy is changed every five years, regardless of the treatment failure rate at the end of every five-year period; (3) adaptive cycling where first-line therapies are switched when the current therapy’s treatment failure rate rises above 10%; and (4) deployment of a triple ACT, either dihydroartemisinin-piperaquine combined with mefloquine (DHA-PPQ-MQ) or artemether-lumefantrine combined with amodiaquine (ALAQ). The current treatment failure rate for each therapy is calculated over a 60-day window of past treatments. In the adaptive cycling approach, a drug switch takes place one year after TF rises above 10% and a new therapy must be used for one year before another switch occurs.

Two new approaches are evaluated in this analysis. First, cycling approaches are split into whether the long half-life partner drugs are used first in the cycling strategy with the short-half life drugs used last – this is an LMS (long-medium-short) strategy – or whether short half-life partner drugs are used first and long half-life drugs last (SML). In an LMS cycling strategy, the order of the therapies is DHA-PPQ first, ASAQ second, AL third. Under SML, the order is AL first, ASAQ second, DHA-PPQ third. Past cycling analyses [5,8] used LMS as this approach reduces malaria prevalence more and earlier than an SML approach. Second, since MFT is non-adaptive, an adaptive MFT approach was created. The adaptive MFT approach begins with 33.3% of cases treated with each ACT, and with readjustments made every two years based on current treatment failures (with a one year delay allowed for implementation). In other words, treatment failure percentages for each ACT are measured at years 2, 5, 8, and 11, and adjustments to ACT distribution are made in years 3, 6, 9, and 12. The purpose of this adaptive MFT approach is to increase the usage of high efficacy therapies and decrease the usage of low efficacy therapies as the resistance landscape changes. Implementation of this scheme would be challenging in a real-world setting but this analysis can tell us how much room for improvement there is for traditional fixed MFT approaches. Details on how the distributions are updated are provided in Supplement A Section 11.

## 3 Results

Our new individual-based 25-locus epidemiological model for *P. falciparum* malaria can faithfully replicate classic epidemiological relationships for malaria, including (1) the relationship among prevalence, incidence, and the entomological inoculation rate (EIR) (Supplement A Figure S14, S15), (2) the relationship between age-specific incidence and EIR (Supplement A Figure S12), (3) the relationship between EIR and the multi-clonality of infections (Supplement A Figure S16), (4) the time-to-emergence period of artemisinin resistance based on approximate timelines in southeast Asia (Supplement A Figure S17), and (5) the efficacies of ACTs on a large range of genotypes known to be partially resistant to artemisinin or the partner drugs co-formulated in ACTs [11–14]. This model was created to have an accurate representation of the *P. falciparum* genome’s loci associated with drug resistance and to have an accurate representation of the recombination process that can put drug-resistance mutations together and break them apart. With an extensive set of simulations we verified that the presence of this new recombination mechanism in the model takes linkage disequilibrium to zero (example in Supplement A Figure S4) when there is no treatment and when fitness costs are set to zero.

In a range of model scenarios, recombination events generating double-resistant genotypes to AL or DHA-PPQ increase – irregularly and slower than linearly – with an increase in the model’s interrupted feeding rate (IFR); see Figure 1. Because the initial genotypes chosen in our simulation were amodiaquine-resistant 76T Y184 genotypes, recombination events generating double-resistants to ASAQ were not seen in the simulations due to the lack of single-resistant genotypes to artemisinin that carried no amodiaquine resistance. Initial recombination counts in a range of generic scenarios made clear that the IFR is unlikely to be the major driver of recombination, as increasing the IFR from 0.0 to 0.2, which is close to the maximum observed in field studies [15–24], showed a 17% increase (under MFT) and a 26% increase (under the two cycling policies) in the total number of double-resistance-generating recombination events recorded in the simulation. Note that the recording of a recombination event is specific to the partner drug, and that only one resistance mutation (or CNV) is required for a genotype to be counted as partner-drug resistant. Additionally, note that in the simulations done for Figure 1, lumefantrine has three mutations associated with resistance while piperaquine has five. When compared to cycling policies, MFT appears to generate more DHA-PPQ double-resistants through recombination but fewer AL double-resistant through recombination. The reasons appear to be that the number of loci that confer resistance to PPQ is higher than the number of loci conferring resistance to LUM, generating more opportunities for recombination under MFT, and (2) DHA-PPQ is used first in the two cycling strategies in Figure 1 (and in Supplement B Figure S1-S4), ensuring that it is used at a time when *pfkelch13* mutants are at low frequencies, reducing the chance of generating a double-resistant through recombination.

**Figure 1:**
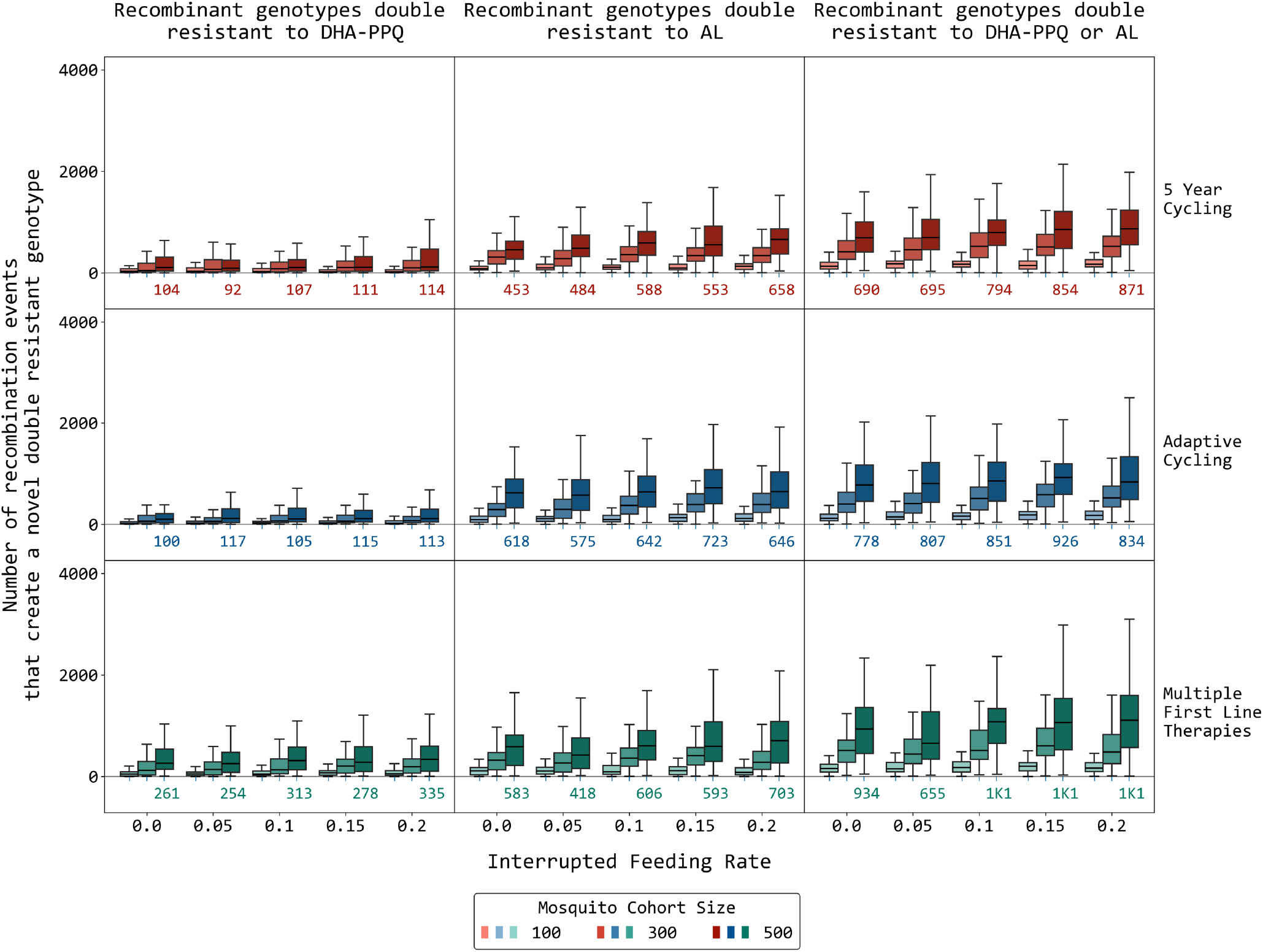
The number of recombination events through year 15 that produce novel recombinant double-resistant genotypes. A recombination event is counted when (1) the recombinant offspring has both a *pfkelch13* mutation and at least one partner-drug associated resistance mutation, and (2) the parental genotypes that recombined carried resistance alleles to only one drug (i.e. one parent carried a *pfkelch13* mutant and was fully susceptible to the partner drug, while the other parent was wild-type at the *pfkelch13* locus and carried at least one mutation to a partner drug). The three rows of panels show recombination counts under three different treatment strategies; cycling is LMS. The *x*-axis in each panel shows the mosquito interrupted feeding rate. The three boxplots in each group show the median/IQR for number of recombination events when the model uses a mosquito cohort size of 100, 300, or 500 (from left to right). Numbers beneath the boxplots on the right indicate the median number of recombination events for a mosquito cohort size of 500. Note that the absolute numbers of recombination events shown in this figure are only meaningful for a mosquito cohort of a particular size. Malaria prevalence is 5% and the daily per-allele fitness cost of resistance is 0.0005. Even though MFT has more recombination occurrences for DHA-PPQ double-resistants, these double-resistants are not necessarily successful and do not spread more quickly as can be seen in Supplement B Figure S6.

When comparing model runs where recombination via single uninterrupted mosquito feeds has been “turned off” to normal runs where recombination can occur in either single feeds or interrupted feeds (as in real life) a count of recombination events by type shows that a median of 80% to 97% of recombination events generating new genotypes (here double-resistants) occurs from single uninterrupted feeds (Figure 2). There did not appear to be large differences when comparing across treatment strategies or across mosquito cohort sizes of 100, 300, or 500, suggesting that a mosquito cohort size of 100 may be adequate for capturing malaria genetic diversity and recombination behavior in a population of 100,000 individuals. The higher or lower frequency of recombination through one process (single feeds) compared to another (interrupted feeds) depends on the fraction of the host population that is carrying multi-clonal infections, the overall genetic diversity of malaria, and the proportion of mosquito feeds that are interrupted. In the 5% and 25% prevalence scenarios evaluated here, 12.1% (IQR: 10.4 -14.6) and 22.5% (IQR: 20.1 – 25.0) of hosts, respectively, have multi-clonal infections. A maximum of 20% of mosquito bites are interrupted, and only 5% or 25% of these will be feeds where *both* hosts are infected, showing why mutliclonality likely plays a larger role in generating recombinant genotypes of malaria. Note that the level of mutliclonality declines with lower prevalence, but so does the probability of a mosquito feeding on two infected hosts.

**Figure 2:**
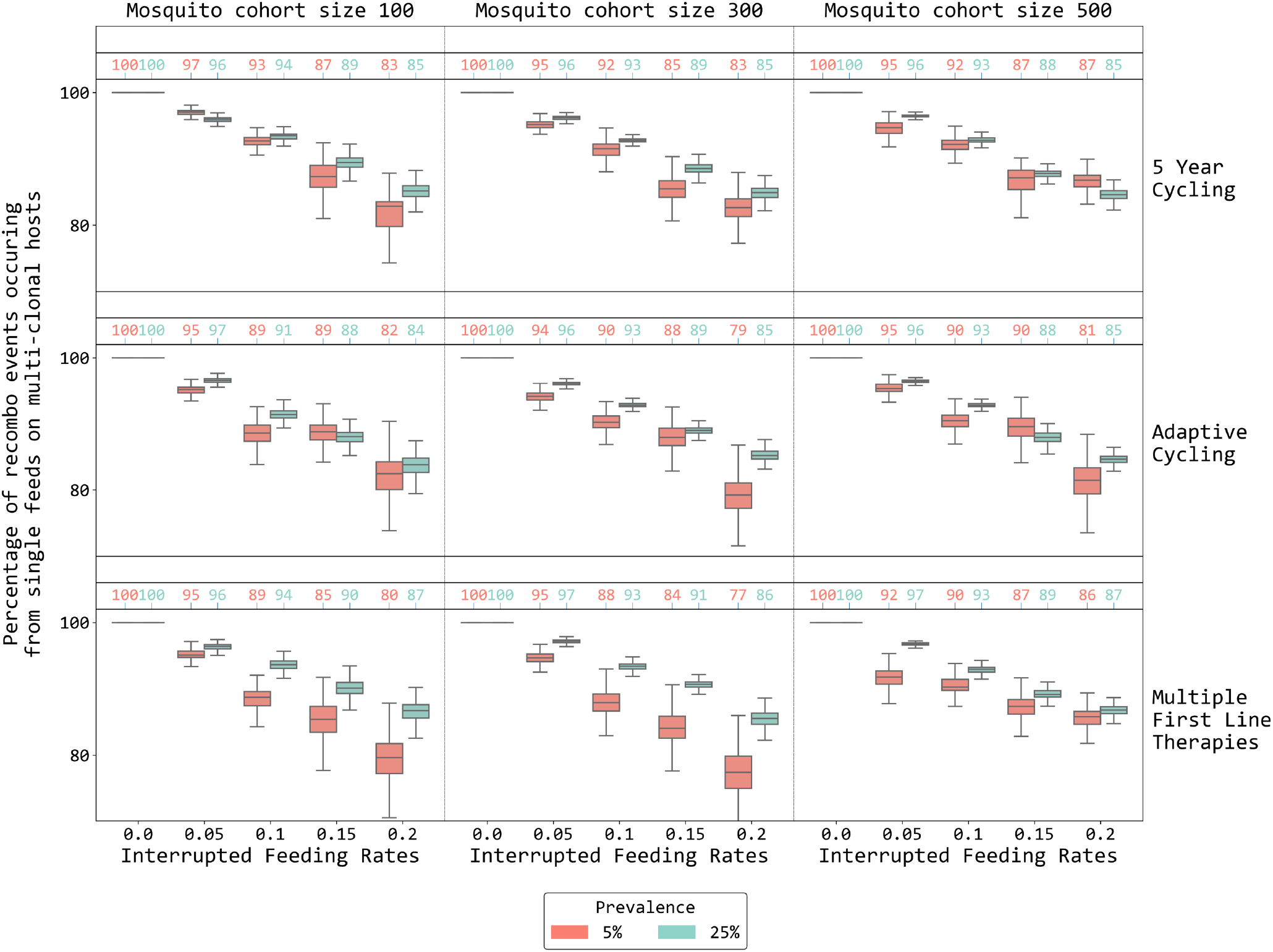
The percentage of recombinations producing double-resistant genotypes that come from single feeds on multi-clonal hosts. The percentage is computed by dividing the number of recombination events (within-host recombination turned off) by the number of total recombination events (within-host recombination turned on) and subtracting from 100%; this is done via sampling with replacement from the empirical recombination counts produced by the simulation. Panel columns indicate different mosquito cohort sizes while panel rows show different strategies applied; cycling is LMS. The *x*-axis indicates interrupted feeding rates of the mosquito while the *y*-axis shows the percentage of recombination events that can be attributed to single feeds on multi-clonal hosts. Red and green depict the percentage computed at 5% and 25% prevalence respectively. Numbers above the boxes are the medians for those box plots. In this figure the daily cost of resistance is 0.0005.

### Strategy Comparisons

The evolutionary effects of treatment strategies on drug-resistance evolution have been studied with a range of different models from classic population-genetic approaches [25,26] to modern spatially-structured individual-based models [2,6]. The general conclusions drawn from these analyses are that maximizing environmental variability slows down drug-resistance evolution. In the language of malaria treatment, parasite populations that encounter more drugs in short time periods will show the slowest or most delayed drug-resistance evolution [27,28]. However, it is not known whether recombination events affecting these evolutionary trajectories are expected to be more common or less common under one treatment strategy compared to another.

Across two prevalence scenarios (5% and 25% PfPR) and two scenarios with differing costs of resistance (17% and 84% annual fitness cost), MFT policies had marginally better outcomes than cycling policies in terms of the number of treatment failures (NTF) they were projected to generate over a 15-year period (Figure 3). In general, cycling policies did not have predictable outcomes when comparing adaptive cycling to 5-year cycling and LMS ordering to SML ordering of therapy deployment. MFT policies tended to have equal or slightly lower NTF ranges than cycling policies – median NTF for MFT ranged from 49% lower to 4% higher when compared to cycling policies for three of the four prevalence/fitness-cost combinations. At 5% prevalence and *c*_*R*_ = 0.0005, MFT-cycling comparisons were inconclusive with two cycling policies projected to have higher NTF counts and two cycling policies projected to have lower NTF counts than MFT.

**Figure 3:**
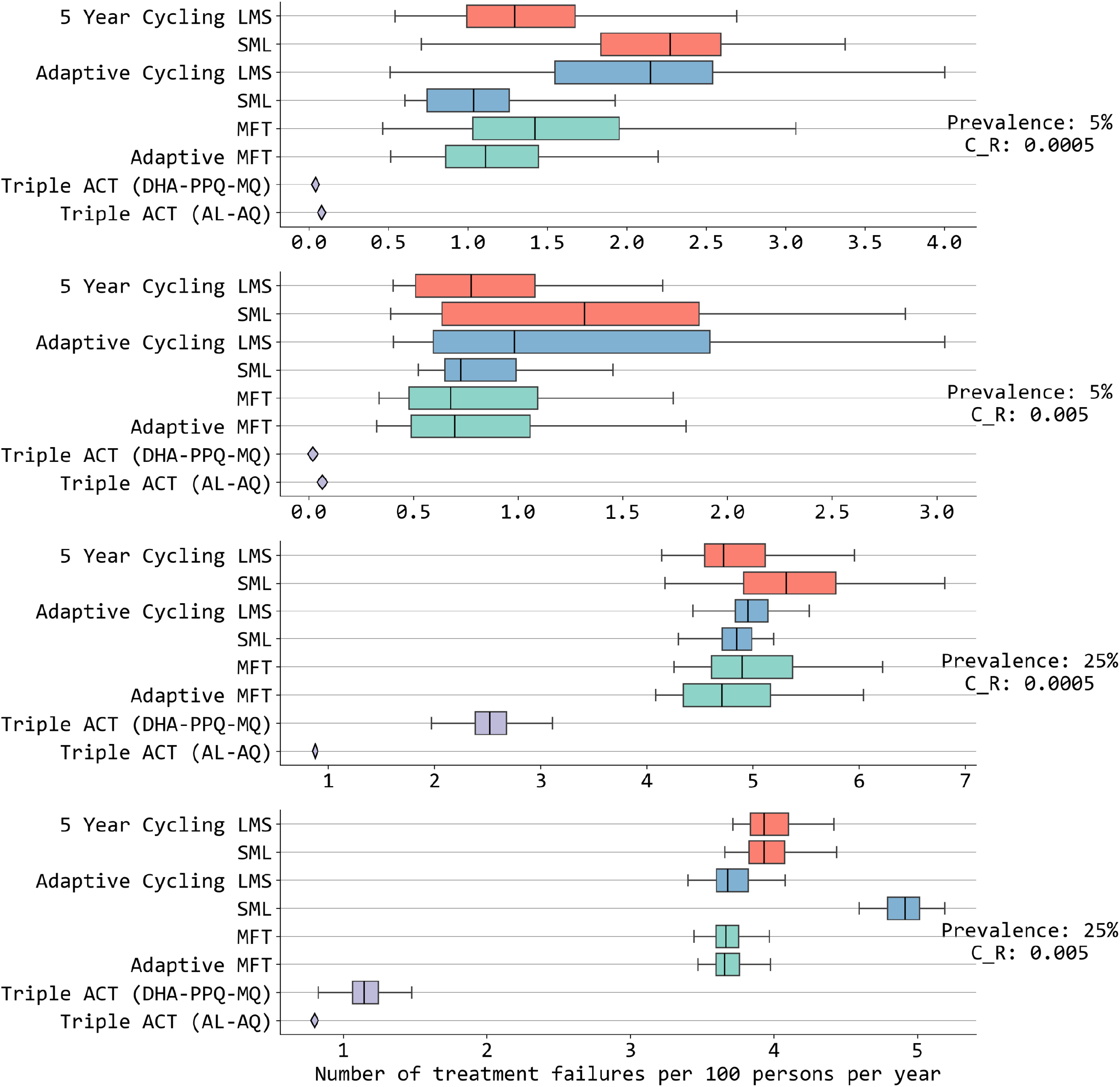
Comparison of annual numbers of treatment failures (NTF) among different strategies. Results shown for 15 years of simulation after applying different treatment strategies in four different prevalence and cost-of-resistance settings. The *x*-axis shows treatment failures per 100 person per year. Different colors are for different treatment strategies: 5-year cycling (red), adaptive cycling (blue), multiple first-line therapies (green) and triple ACT (violet). LMS indicates the order of drugs used in cycling strategies based on half-life of partner drugs (**<u>L</u>**ong: DHA-PPQ, **<u>M</u>**edium: ASAQ, **<u>S</u>**hort: AL). A single diamond corresponds to a boxplot where the IQR range is smaller than 0.05. Among these strategies, triple ACT use stands out with the lowest number of treatment failures for both low and high prevalence. MFT and adaptive MFT generally (but not always) have lower NTF values than cycling approaches. The mosquito cohort size is 100 and the interrupted feeding rate is 20%.

Cycling policies in some instances showed improvements over MFT policies. As noted in previous studies [6,7], MFT policies are predicted to delay and slow down drug-resistance evolution by a larger margin than cycling policies when all therapies (1) have equal efficacy currently and (2) are projected to have equal efficacy into the near future when drug resistance is expected to evolve. In our simulation set-up, certain amodiaquine-resistant genotypes are fixed at the beginning of the simulation (as they would have been in 2005 when ACTs were introduced) and AL efficacy is high (>92%) on these genotypes; thus, an adaptive cycling policy that uses AL first is likely to have high cure rates over a long period and may thus be favored over an MFT policy where all three ACTs are used equally. In the LMS 5-year cycling approach in Figure 3A, DHA-PPQ is high-efficacy, used first, and replaced after five years before *pfkelch13* variants are able to emerge and erode its efficacy; under SML 5-year cycling however, DHA-PPQ is used in years 10-15 when *pfkelch13* are likely to be circulating at higher levels, resulting in the highest levels of treatment failure seen among the three ACTs. This explains why LMS has more favorable drug-resistance mitigation outcomes than SML in this particular scenario, but there does not appear to be a general rule on which ordering of ACTs is optimal in a cycling approach (see *Discussion*). Note that adaptive cycling policies sometimes have equal resistance and TF outcomes to MFT policies, but adaptive cycling approaches are sensitive to the delay period between identifying resistance and switching therapies, leading to substantial increases in treatment failure counts when delays are long (Supplement B Figure S9). Adaptive MFT policies showed small and inconsistent improvements over fixed MFT policies, largely due to these adaptive policies’ small adjustments in drug distribution making their overall therapy use similar to a fixed MFT policy.

By far the most conclusive result, via a direct comparison of treatment failures counts, was that deployment of triple ACT was projected to have substantially lower NTF over a 15-year period than any cycling or MFT approach. Deployment of either triple ACT showed treatment failure counts that were 48% to 97% lower than the other strategies, confirming the findings of Nguyen et al (2023) and Zupko et al (2023) [2,7].

When looking at a drug-resistance prevention or delay metric – the time until an artemisinin-resistant genotype reaches 0.01 frequency in the host population (T_.01_) – MFT policies generally outperformed cycling policies, and LMS appeared to have small advantages over SML cycling policies. In six of the eight LMS-to-SML comparisons in Figure 4, LMS rotations had times to emergence that were 7.1% to 28.2% longer than SML rotations; in the remaining two comparisons T_.01_ was 2.1% and 2.5% shorter for LMS. If this behavior can be shown to be robust, it may be the result of piperaquine resistance (emerging first under LMS) requiring more mutations than lumefantrine resistance (Supplement B Figure S15). For this same reason – with the three therapies not being identical in their (a) current efficacy, (b) their future resistance profiles, and (c) the number of known mutations conferring resistance – cycling policies are able to sometimes outperform MFT policies if they use their best therapies first. As an example, at 5% prevalence and *c*_*R*_ = 0.0005, SML 5-year cycling had an emergence time T_.01_ that was 1.3% longer than under MFT, and LMS adaptive cycling had a T_.01_ that was 7.6% longer than under MFT. For the remaining 14 of 16 comparisons between cycling and non-adaptive MFT in Figure 4, MFT had times to emergence that were 1.3% to 47.9% longer.

**Figure 4.**
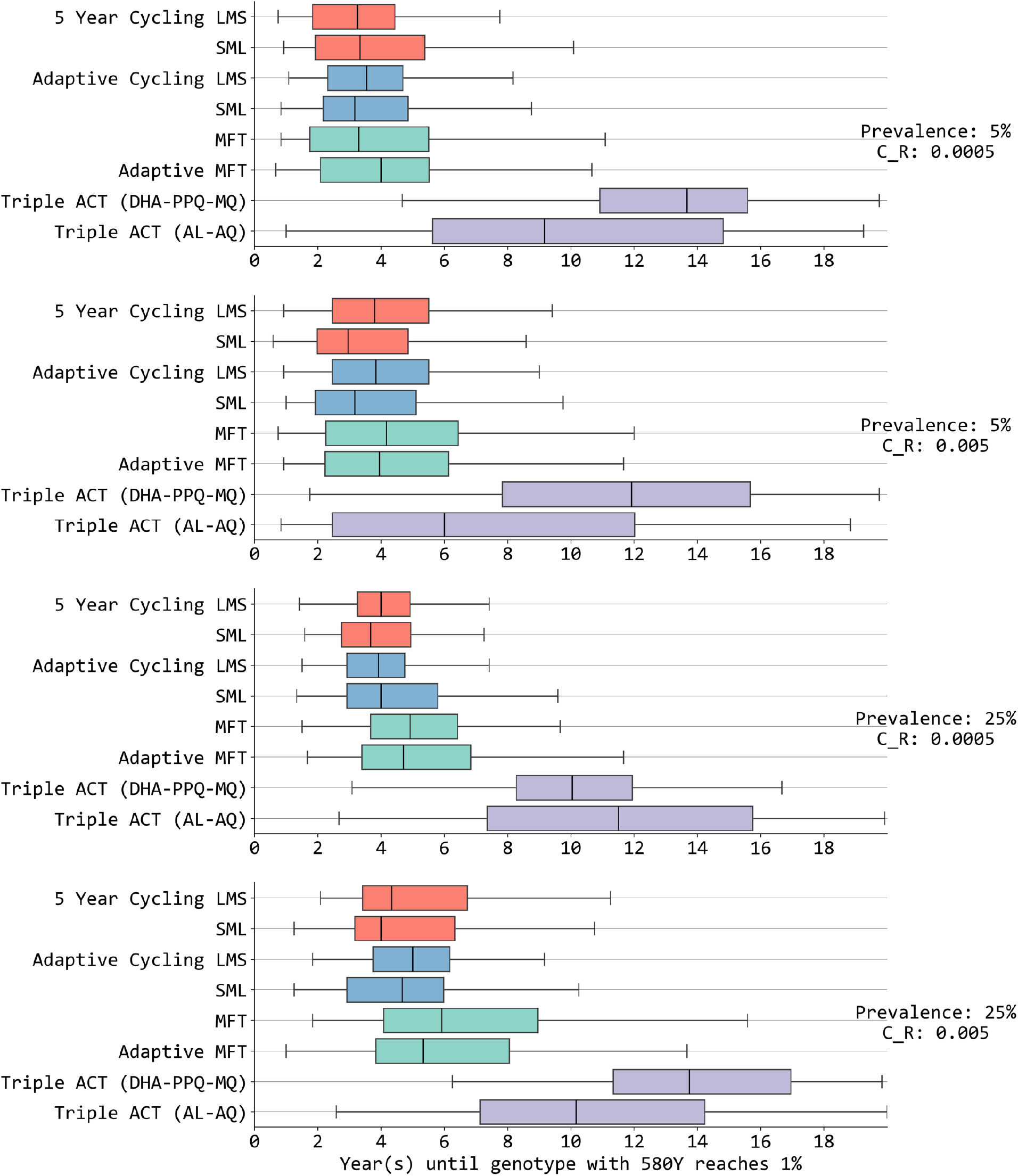
Number of years (*x*-axis) until a single-mutation *pfkelch13* variant reaches 0.01 allele frequency. Results are shown for four different prevalence and cost-of-resistance settings. Different colors are for different treatment strategies: 5-year cycling (red), adaptive cycling (blue), multiple first-line therapies (green) and triple ACT (violet). LMS indicates the order of drugs used in cycling strategies based on half-life of partner drugs (**<u>L</u>**ong: DHA-PPQ, **<u>M</u>**edium: ASAQ, **<u>S</u>**hort: AL). Triple ACT use stands out with the longest delay until artemisinin-resistance emergence. MFT and adaptive MFT are generally (but not always) associated with longer emergence times than cycling approaches. The mosquito cohort size is 100 and the interrupted feeding rate is 20%.

Again, the longest delay in artemisinin resistance’s time to emergence occurred, by a large margin, under triple ACT strategies. When using triple ACTs, the model projections showed that the median time until 0.01 genotype frequency of artemisinin resistance was 1.7 times to 3.4 times longer than under the best cycling or MFT policies.

### Risk of Multi-Drug Resistance

To evaluate the risk of early emergence of multi-drug resistant (MDR) genotypes, we counted *early recombination events* that generated double-resistant or triple-resistant genotypes. An *early recombination event* that generates a double-resistant genotype (from two single-resistants) is defined as one that occurs before the double-resistant reaches 0.01 genotype frequency. In other words, this count captures the occurrence of recombination when it is likely to have its largest impact – during a time period when double-resistance is still rare. Similarly, an *early recombination event* generating a triple resistant (from a single- and a double-resistant) is one that occurs prior to the triple resistant reaching 0.01 genotype frequency. MDR genotypes can be produced by either recombination or mutation, and they can increase in frequency due to selection pressure; all of these processes can be individually tracked during a simulation run as shown in Figure 5. The three recombination events shown in years 5 to 10 of this figure (open circles on orange lineage) are early recombination events.

**Figure 5.**
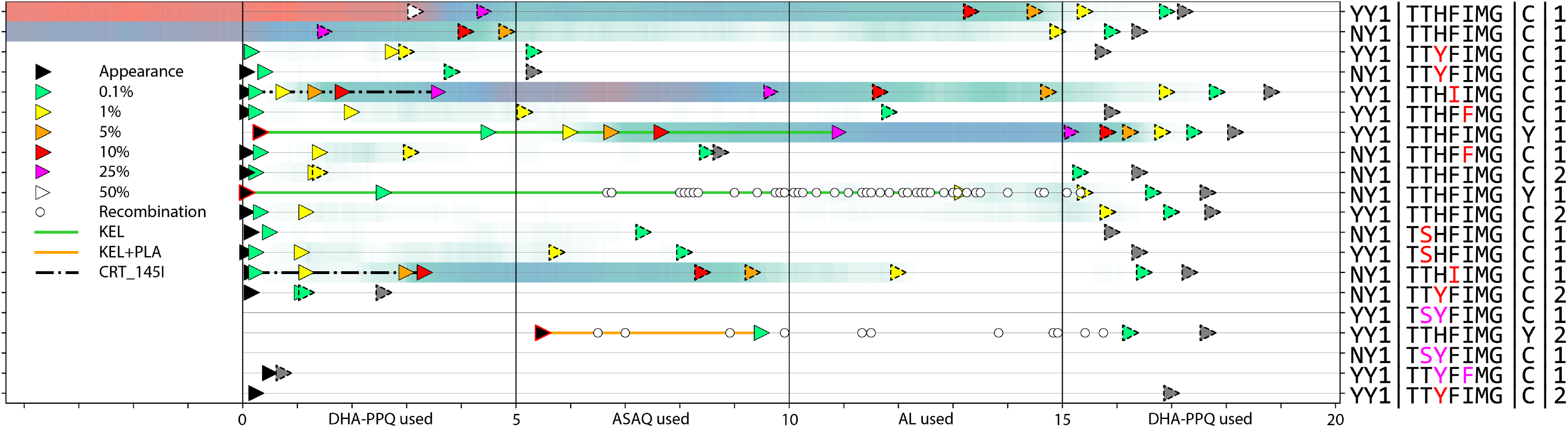
Evolution from a single 20-year model simulation of a 5-year cycling strategy with DHA-PPQ used in the first five years, ASAQ in the following five years, AL during years 10-15, and DHA-PPQ again in years 15-20. Prevalence is 5% and cost of resistance is set to *c*_*R*_ = 0.0005. Graph above shows evolution of 20 chosen genotypes (rows) with colored triangles showing genotype frequency milestones (see legend at left). Triangles with dashed borders indicate that a genotype frequency is decreasing, and the dark gray triangles indicate that a genotype has disappeared. The darker sea green and lighter sea green shadings correspond to genotype frequencies in the 0.05 to 0.25 range. Circles show when a genotype is generated through a recombination event, and the three circles on the orange KEL+PLA lineage are *early recombination events*. Genotype codes at right show genes on chromosomes 5, 7, 13, 14, separates by vertical lines. The first group of alleles (chromosome 5) shows the alleles at position N86Y, Y184F, and the copy number of the *pfmdr1* gene. The second group of alleles (chromosome 7) shows seven *pfcrt* loci with mutations at positions 76, 93, 97, 145, 218, 343, and 353 highlighted in red (single mutants) and pink (double mutants); these are the key mutations (except position 76) known to reduce PPQ efficacy. Colored lines show certain key lineages like the KEL-lineage where a single *pfkelch13* mutation is present but plasmepsin copy number (the “1” on the RHS) has stayed at single copy. The “145I” lineage (seventh from bottom) is associated with substantial reduction in PPQ efficacy and increases quickly from years 0 to 5.

The total count of early recombination events generating double resistance (Figure 6, top panels) is 11% to 21% lower under MFT than under cycling policies, despite the emergence times being longer under MFT. As in Li et al [8], simulations run under an MFT strategy do not appear to generate any additional MDR emergence risk even with all therapies deployed simultaneously. The likely reason for this effect is that MFT will tend to keep novel alleles rare for longer periods when compared to cycling policies, making recombination events ‘doubly rare’ when they have to recombine two rare types to create a double- or triple-resistant. Similarly, the total count of early recombination events generating triple resistance is 15% to 27% lower under MFT than under cycling policies (Figure 6, bottom panels).

**Figure 6.**
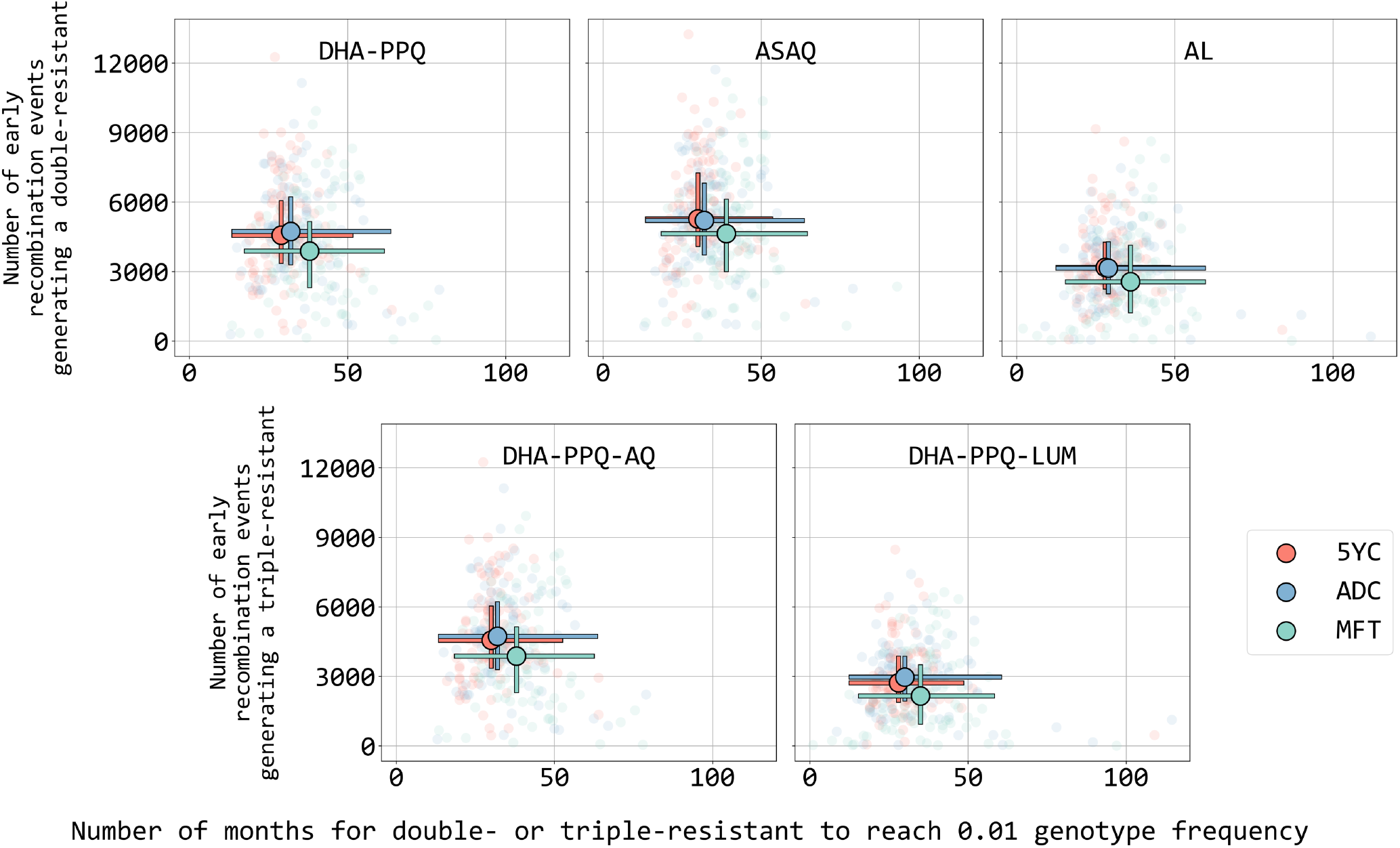
Number of recombination events (*y*-axis) creating a double-resistant or triple-resistant genotype to a particular ACT or triple ACT (see text inside each panel) that occur prior to the time that the frequency of this MDR genotype reaches 0.01. The *x*-axis shows the number of months, starting from first appearance, for each MDR genotype to reach 0.01 genotype frequency. Circles show medians and bars show interquartile ranges from 100 simulations from three different treatment strategies: MFT (green), 5-year LMS cycling (red), and LMS adaptive cycling (blue). Prevalence is 5%, *c*_*R*_ = 0.0005, mosquito cohort size is 100, and the interrupted feeding rate is 20%.

Times to reach certain triple resistance milestones can only be counted for artemisinin/piperaquine/amodiaquine triple resistance and artemisinin/piperaquine/lumefantrine triple resistance. As noted previously [7], the triple ACT ALAQ does not naturally drive any known triple resistant genotype as currently known resistance markers for these drugs indicate that lumefantrine-resistant parasites will be amodiaquine-sensitive and that amodiaquine-resistant genotypes will be lumefantrine-sensitive. For this reason, only two triple resistant genotypes are tracked in Supplement B Figure S7 which shows that time to triple resistance is 56% to 95% longer under MFT than under cycling polices. Using DHA-PPQ-MQ as first-line treatment, time to emergence of the triple-mutant is 1.3 to 2.3 times longer than under MFT or cycling policies. Note that in all these scenarios the triple mutant starts at zero when the simulation starts; pre-existence of a triple-mutant is a risk factor for deploying triple ACTs [7].

## 4 Discussion

With a novel and more genetically accurate model of drug-resistance evolution in *Plasmodium falciparum* malaria, we are able to identify which recombination process in malaria has a larger effect on generating multi-drug resistant genotypes through recombination – specifically, we find that recombination in falciparum parasites is primarily generated by multi-clonal infections and less often by interrupted mosquito feeds. Additionally, a comparison of drug-resistance management approaches with this new model shows that triple therapy outperforms MFT, and that MFT outperforms cycling approaches, using a range of different outcome measures. Finally, MFT strategies do not appear to accelerate MDR evolution when compared to cycling approaches, reaffirming a recent result on this specific question [8].

There continues to be a need for the development of general theory on the question of whether recombination acts more strongly to bring drug-resistance mutations together into MDR genotypes or to break apart MDR genotypes into less resistant offspring. On the one hand, one could argue that when drug resistance is rare recombination should have a stronger effect on breaking apart multi-genic drug resistant genotypes than it does at putting them together. The rationale behind this is that a multi-genic resistant genotype faces the possibility of recombination (and thus breaking apart of multiple resistant alleles in the same genome) in at least 10% of transmission events (this is the multi-clonal rate at 5% prevalence). However, recombination bringing two rare resistant alleles together requires two rare types to be sampled by the same mosquito – something that is *O*(*x*^2^) rare when single-resistants are circulating at genotype frequency *x*. One the other hand, higher recombination rates should lead to *earlier* emergence of the first recombinant double-resistant genotypes. Even if such a recombination event is rare, several emergence events may be sufficient to bring MDR types up to high levels if positive selection is strong enough. Recombination breaking up these MDR types may occur in large numbers when MDR genotype frequencies are high, but break-up by recombination may not be stronger than positive selection when MDR genotype frequencies are low. To the best of our knowledge, it is not known how these forces balance each other out.

A second open question in drug-resistance evolution that does not appear to have a general treatment in the literature is the effect that the order of drugs in a cycling approach has on drug-resistance outcomes. Traditional analyses in this field [29–31] have assumed that the considered drugs or antibiotics are equal. In reality however, different drugs can have different (1) efficacies, (2) efficacies on future resistant genotypes, (3) mutation rates to resistant types, (4) numbers of mutations conferring resistance, and (5) pharmacokinetic clearance properties. When all five of these properties are equal among the considered therapies, past analyses have shown that MFT outperforms cycling policies; however when these quantities vary among drugs we may see situations when using the ‘best drugs first’ in a rotation strategy is preferable to MFT [6]. When variation exists across all of these properties (as for currently used ACTs) it is not clear which ACT has some optimal and undefined set of characteristics of high treatment success and avoidance of high-grade resistance evolution. A general study investigating an approach to ranking these properties according to their propensity to speed up or slow down resistance evolution would go a long way, in malaria policy, in helping countries decide which ACTs to prioritize for deployment and therapy switches. Finally, this analysis would have to consider whether – under the most favorable surveillance and deployment conditions – adaptive cycling policies (i.e. ones that trigger switching only when treatment failures meet a certain threshold) are superior to pre-planned fixed rotation strategies. It may be the case that early switches in adaptive policies have detrimental long term effects by moving a health system to use the wrong therapy at the wrong time.

### Practical implications

The applied purpose of these research questions is to lay the foundation for scenario-specific national-level mitigation strategies for the artemisinin-resistant genotypes that began circulating in Africa last decade. Our previous effort in designing such a strategy for Rwanda [2] used information at just six key loci and was not able to accurately model the full range of pathways to piperaquine resistance. This is critical as DHA-PPQ and ASAQ are the next two ACTs that will be considered in most national contexts as AL use is reduced or phased out. Clinical trials from SE Asia suggest that PPQ deployment may come with high risks of treatment failure [14]. The efficacies shown in these trials, some close to 25%, may represent a worst-case scenario for DHA-PPQ use or may be dependent on certain immunological or parasitological features specific to SE Asian malaria settings. Deployment of DHA-PPQ in Africa is not guaranteed to follow the same pattern seen in SE Asia, but it should nevertheless be accompanied by near real-time molecular surveillance so that PPQ-resistant genotypes can be identified shortly after they emerge. Pyronaridine-artesunate (PYAS) is the fourth ACT that will be considered for deployment. PYAS enjoys high efficacy [32] and a lack of any known pyronaridine-resistant phenotypes, but suffers from high prices and low supply. If PYAS is deployed in an African context and resistant genotypes are found and characterized, the present model will allow for response planning to this emergence event.

A feasible and practical option that is currently being planned by three African National Malaria Control Programs is fast rotation of ACTs [33] – both as preparation for the arrival of resistant genotypes and to slow geographic spread of recently introduced kelch13 variants. The pre-planned rotation periods that are being considered are one, two, and three years, and the shortest of these is likely to see resistance mitigation benefits similar to those of MFT. An early field study indicated that a particular type of stock management will make this rotation feasible in many health-system settings [34] and general social and behavioral studies have indicated that patients, providers, and health systems are ready for a changeover to a rotation or MFT-like approach [34–37]. Evaluating the early emergence and disappearance of resistance alleles in this fast rotation scheme will be critical to understanding which facets of these strategies work better and worse when the cycling period is shortened.

## Limitations

This is the first model, to our knowledge, to compare the relative frequencies of recombination through multi-clonal host sampling and interrupted feeding. Further corroboration, through independent modeling efforts or field studies looking at parasite diversity within mosquitoes, will be necessary to build an evidence base showing that one type of recombination event is more common than the other. As in all large complex individual-based pathogen transmission models, model behavior needs to be continuously validated against field data. As an example, the model’s cost of resistance *c*_*R*_ has been kept deliberately low in comparisons of MFT and cycling approaches because high *c*_*R*_ values in models generally result in MFT having large advantages over cycling policies. However, for practical application and development of national-level strategies, it is more important that *c*_*R*_ estimates are accurate than conservative. Future development in this area will likely combine field data [38] with in vitro data [13] to obtain realistic estimates of fitness cost for drug-resistance mutations in *P. falciparum*.

The major challenge in the evolutionary epidemiology of *P. falciparum* for the near to medium future is building and expanding a genotype-phenotype map for drug-resistant genotypes. Our original drug-by-genotype map, developed for six loci, required substantial approximation as direct treatment-failure data are rarely available for specific genotypes. In the current model version – with 25 relevant loci or CNVs – there are now millions of genotypes and dozens of phenotypes to define (assuming that all of these can emerge and provide some resistance benefit). The sparsity of data for this problem means that substantial uncertainty will remain for the near future on the behaviors, pharmacodynamics, and treatment efficacies on these genotypes.

Finally, additional corroboration will be needed on resistance emergence times for the major ACTs used in Africa and SE Asia. Our model’s mutation rate is currently calibrated against approximate historical data for SE Asia, but not yet adequately to the conditions in the 2010s that led to the emergence of kelch13 variants in Africa under persistent use of artemether-lumefantrine. The decision in SE Asia in the early 2000s to rely on DHA-PPQ and the decision in Africa to rely primarily on AL over the past twenty years could have been an addition major factor (higher drug coverage and lower immunity being two others) influencing earlier emergence of artemisinin in SE Asia and later emergence in Africa.

The global malaria community, together with endemic-country national malaria programs and supporting international institutions, has a major challenge ahead in its efforts to slow the spread of and mitigate the effects of currently circulating artemisinin resistance [39,40]. Novel non-artemisinin therapies and triple combination therapies may be available later this decade. In the meantime, optimal use of current therapies appears to be the only major national-level option available as a specific remedy to spreading drug-resistance. Accurate, robust, and feature-rich mathematical models of malaria transmission will be a key tool in this optimization and treatment strategy design process. A commitment to model evaluation, strategy design, feasibility discussions, and constant iteration of this process will be the key to ensuring that artemisinin-resistant genotypes do not establish themselves continent-wide in Africa by the 2030s.

## Supporting Information

Supplementary Materials A – Full model description and model diagnostics for version 5 of individual-based stochastic simulation of *P. falciparum* transmission and evolution.

Supplementary Materials B – Sixteen supplementary figures showing sensitivity analyses and robustness checks.

## Acknowledgements

KTT, TDN, MFB were funded by National Institutes of Health grant NIAID R01AI153355 and Bill and Melinda Gates Foundation grant INV-056612. KTT, TDN, MFB, TN-AT, RJZ were funded by National Institutes of Health grant NIAID R01AI153355 and Bill and Melinda Gates Foundation grant INV-005517. EZL was funded by the University of Washington’s Malaria Modeling Consortium grant from the Bill and Melinda Gates Foundation (OPP159934) and by Bill and Melinda Gates Foundation grant INV-005517. DBW funded by NSF grant 2146260. JLS-S is grateful for support from NIH (K08AI163497 (PI JLS-S) and DP2AI190432 (PI JLS-S)) and a Doris Duke Charitable Foundation Physician Scientist Award (Grant 2019121) and a Louis V. Gerstner, Jr. Award. SM acknowledges the support from the Human Frontier Science Program Long term Fellowship (LT000976/2016-L) and funding from National Institutes of Health NIAID grant R01 AI182318. TB supported by a European Research Council (ERC) Consolidator Grant to Teun Bousema (ERC-CoG 864180; QUANTUM). Computations for this research were performed on the Pennsylvania State University’s Institute for Computational and Data Sciences’ Roar supercomputer and on Temple University’s College of Science and Technology Owl’s Nest Cluster.

## Author Contributions

**Conceptualization:** Kien Trung Tran, Tran Dang Nguyen, Maciej F Boni

**Formal analysis:** Kien Trung Tran

**Funding acquisition:** Maciej F. Boni

**Investigation:** Kien Trung Tran, Tran Dang Nguyen, Daniel Weissman, Eric Zhewen Li, Robert J Zupko, Thu Nguyen-Anh Tran

**Methodology:** Kien Trung Tran and Tran Dang Nguyen

**Supporting Data:** Teun Bousema, Sachel Mok, Jennifer L Small-Saunders

**Software:** Kien Trung Tran

**Supervision:** Maciej F Boni

**Visualization:** Kien Trung Tran

**Writing – original draft:** Kien Trung Tran, Maciej F Boni

**Writing – review & editing:** Kien Trung Tran, Maciej F Boni

## Supplementary Materials A for

Effects of recombination on multi-drug resistance evolution in *Plasmodium falciparum* malaria by Tran, Nguyen, Weissman et al (2024)

### 1 Basic Model Description for Malaria Simulation Version 5

The simulation model described here is called version 5 and was developed on top of version 3.3 whose most recent publications and descriptions are Li et al [1] and Nguyen et al [2]. Version 4 is the spatial version [3–6] which was built on top of version 3.3. The spatial structure from version 4 is not included in version 5.

First, the genotype structure was upgraded to include all 14 chromosomes explicitly to be able to include *Plasmodium falciparum* resistance markers and their clinical phenotypes (or approximations of these clinical phenotypes by in vitro phenotypes). The reason for this change was the increased number of recently identified drug-resistance markers in *P. falciparum*, especially to piperaquine [7–10]. A major technical upgrade to the codebase was necessary here as version 3.3 used a traditional population-genetic recombination table (size 2^*m*^ × 2^*m*^ × 2^*m*^) where *m* is the number of loci — this becomes computationally challenging once *m* ≥ 10. Version 5 of the model includes 23 loci with known single amino-acid changes associated with drug-resistance and two copy-number variations for the *pfmdr1* gene on chromosome 5 and the *plasmepsin-2*,*3* genes on chromosome 14.

The second new feature, added to facilitate handling a large number of loci, was a mosquito population (or cohort) that explicitly bites individuals both to sample from infected individuals and to inoculate new individuals (infected and uninfected) with malaria. Recombination now takes place as an individual event in a mosquito that has just sampled parasites from humans during a blood meal. Mosquitoes may bite two different hosts during a feed – this is called interrupted feeding. The mosquito cohort, its history, and its infecting parasites are all tracked for 11 days so that infectious bites occurring today are from mosquitoes who sampled parasites 11 days ago. The mosquito cohort size is 500 mosquitoes, which was chosen to allow for rare genotypes circulating at frequencies between .001 to .01 to be sampled by mosquitoes in the simulation. This cohort is not intended to represent all mosquitoes with all parasites they would be carrying. Rather, it is meant to be a cohort that holds a representative sample of the genotypes circulating in the human population. Tests were run with cohort sizes of 100, 300, and 500 and no substantial differences in simulation speed or evolutionary outcomes were seen for these different cohort sizes.

The remaining core components of the model are largely unchanged. The model can be viewed as a population of individual agents (humans) that are updated stochastically using a daily time-step via an asynchronous event management system. Each person in the population has basic attributes and exhibits certain behaviors that impact malaria infection such as (*i*) age and mosquito attractiveness – assigned once at birth or at the beginning of the simulation, or (*ii*) a decision to seek treatment and take a particular antimalarial – randomly drawn each time independently after a malaria fever event for each individual, or (*iii*) the number and genotype of different malaria clones that the person carries – which is dependent on recent biting history and recent treatment history.

### 2 Genotype Model

The new model stores genotype information for each within-host clonal parasite population via a string structured like so

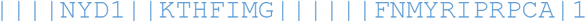

with the 13 vertical bars representing chromosome separators and the individual string characters corresponding to amino-acids or other genetic features such as gene copy number. The string structure maps to an integer key in a genotype database and the string itself is used for filtering and parsing output. In the notation above, as an example, the NYD1 characters appearing after the four vertical bars correspond to a particular genotype of the *pfmdr1* gene which sits on *P. falciparum* chromosome 5. This genotype is <u>N</u>86Y, <u>Y</u>184F, <u>D</u>1246Y which is the wild-type genotype of *pfmdr1*, with the “1” indicating that a single copy of *pfmdr1* is present on chromosome 5. The mutations 86Y, 184F, and 1246Y correspond to alleles whose drug resistance properties have been previously described; details in section 13.

In addition, the simulation has built in functionality to handle multiple genes on the same chromosome, through a data structure based on this string configuration

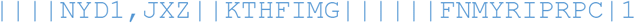

where chromosome 5 now has a second gene, with three relevant loci, coded into the simulation’s genotype database. This is done to allow for intergenic recombination rates on the same chromosome to be specified but is also not used in the current version as no two major resistance-associated genes sit on the same *P. falciparum* chromosome. Table S1 below described all loci used in version 5 of the simulation.

**Table S1:**
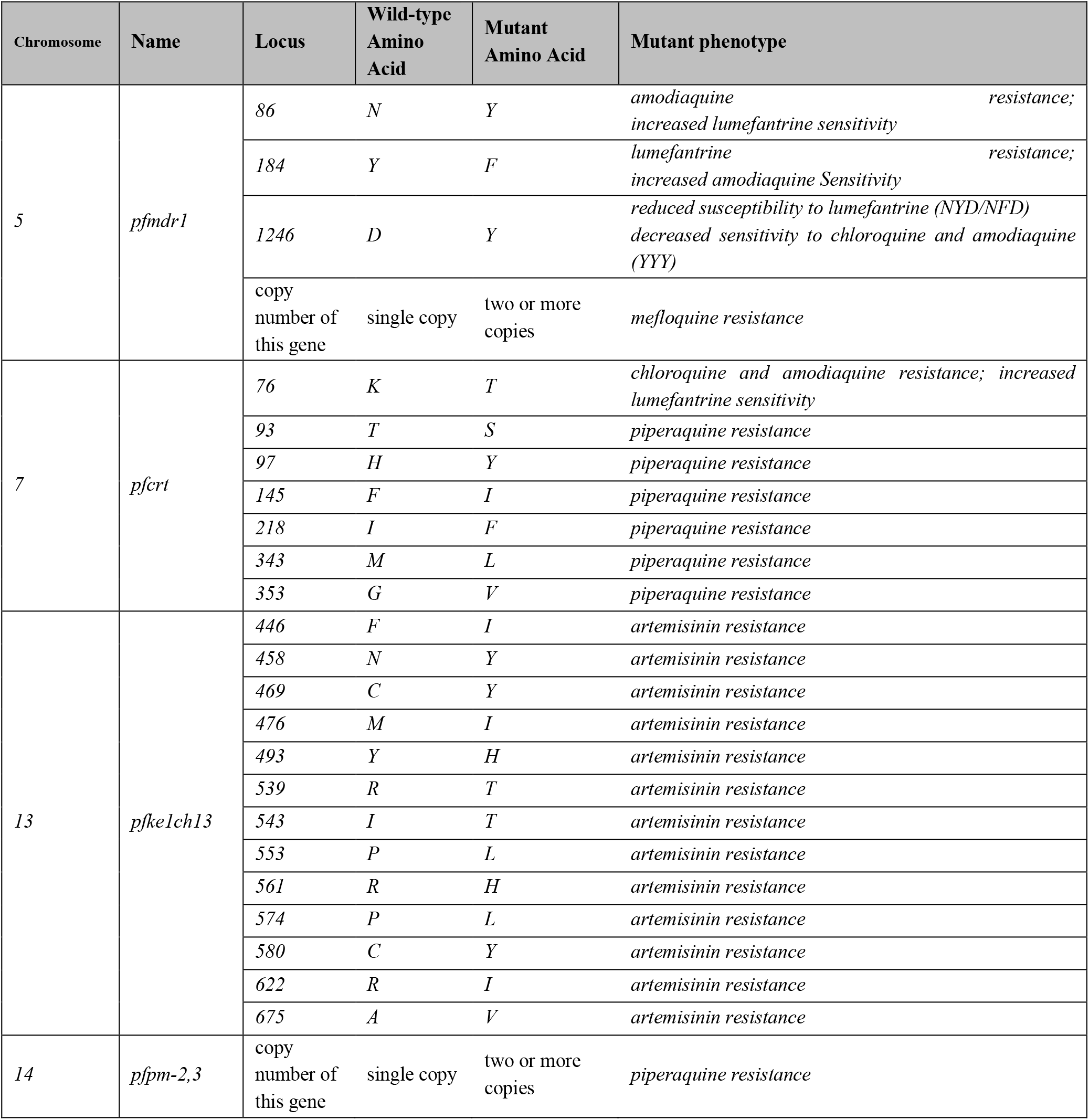
All loci and copy number variants used in version 5 of the simulation.

A second string is used as a mask to allow or disable mutation at particular loci, with 0 and 1 corresponding to disable and allow, respectively, like so

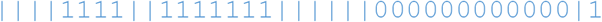

This feature is used for simulation burn-in (if we know for example that *pfkelch13* mutations have not yet appeared in a region) or for specific sub-analyses comparing genotypes or calibrating genotype behavior. For the simulation analysis in the main paper, these masks were used to allow mutation or copy-number variation at 10 positions; see section 2.1 of the main paper.

With this new genotype structure, the model has capability to encode new resistance markers in the future as they are discovered and confirmed. The challenge is that the model also requires the phenotype information for all parasites (across all genotypes) carrying any new allele introduced into the system.

As in all population-level malaria simulations [11] mutation occurs within-host and assumes that the new mutant genotype replaces the resident genotype in a very short period after mutation occurs (i.e. during the course of treatment).

### 3 Multi-clonal infections

Each host can be infected with multiple parasite populations of different genotypes (or identical genotypes); these parasite populations are sometimes described as ‘clones’ or ‘clonal populations’ and they exist (in our simulation) only at the erythrocytic stage of the parasite life cycle. Each parasite clone has its own parasite density. To describe the density of each clone, let *c*_*i*_ be the number of parasite clonal populations in host *i*, and let *j* be the index (from 1 to *c*_*i*_) of each clone in host *i*. Let *D*_*i*_ denote the total density of transmissible parasites (i.e. stage 5 gametocytes, but these are not modelled explicitly) in host *i*. Each parasite clonal population *j* can be any genotype described in section 2. The mathematical notation for parasite density *D*_*i*_ in host *i* is:

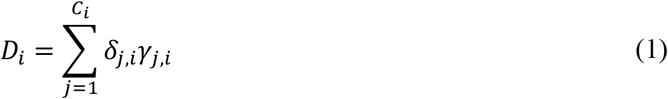

In equation (1) above, we use *δ*_*j*,*i*_ to describe the asexual parasite density (stored as a per µl quantity in the simulation) of clone *j* in host *i*, and we use *γ*_*j*,*i*_ as a binary variable to denote the presence or absence of gametocytes for clone *j* in host *i*.

The binary *γ*-value is set to zero for the first four (children) or six (adults) days of an infection and then fixed at one for the remainder of the infection. Gametocytaemia is not modelled explicitly. Transmissibility of an infection to a mosquito is based on the relationship between asexual parasite density (total, across all clones) and the probability that at least one functional gametocyte is taken up during a mosquito bite. This probability is shown in section 7 and is derived from Ross et al [8], with an alternate description provided in Nguyen et al [12] (see supplementary materials, pp.6-7).

### 4 Parasitaemia Model

Parasitaemia is tracked independently for each clone. In general, the parasite density of a person can be classified into 9 different levels, or thresholds, or types which are described below:

**Table S2:** Parasite log-density thresholds and levels used in the model. In the middle columns, the per µl value is also shown on a log_10_ scale.

| Variable name | Parasites per $\mu\text{l}$ | Usage in simulation |
| --- | --- | --- |
| *log_parasite_density_cured* | 0.00002 per $\mu\text{l}$<br>(value = -4.699) | When a person is consider cured from a particular parasite infection. This corresponds to 100 total parasites for this particular clone, and when the parasite population falls below this level, it is simply set to zero. |
| *log_parasite_density_from_liver* | 0.01 per $\mu\text{l}$<br>(value = -2.0) | This is the number of parasites in the blood immediately after the first group of merozoites bursts from the liver into the blood stream. This corresponds to about 55,000 total parasites. |
| *log_parasite_density_asymptomatic* | 1,000 per $\mu\text{l}$<br>(value = 3.0) | Approximate number of parasites per $\mu\text{l}$ for an asymptomatic infection. For an individual who is bitten but does *not* progress to symptoms, a log-parasitaemia $y$ is drawn from a normal distribution with mean 3.0 and standard deviation 0.5, and this person is assigned a starting point of $10^y$ total parasites in the blood. Afterwards, this parasitaemia declines slowly due to immune clearance. |
| *log_parasite_density_clinical_from* | 2,000 per $\mu\text{l}$<br>(value = 3.301) | Bottom boundary for number of parasites for a person experiencing their first day of fever after an infectious mosquito bite. |
| *log_parasite_density_clinical_to* | 200,000 per $\mu\text{l}$<br>(value = 5.301) | Upper boundary for number of parasites for a person experiencing their first day of fever after an infectious mosquito bite. For a febrile patient, the parasitaemia at fever is drawn uniformly on [3.3, 5.3] with a resulting parasitaemia between $10^{3.3}$ and $10^{5.3}$. |
| *log_parasite_density_detectable* | 10 per $\mu\text{l}$<br>(value = 1.0) | Threshold to determine if a person still has parasites after treatment on day 28. This is a lower threshold than below because microscopy detection limit is lower in a research setting. |
| *log_parasite_density_detectable_pfpr* | 50 per $\mu\text{l}$<br>(value = 1.699) | Threshold to determine if a person has parasites and is counted as ‘parasite positive’ in a PfPR survey. This is based on a typical microscopy threshold that is achievable in a field setting. |

### 5 Duration of infection

For an asymptomatic and untreated infection, the duration of infection varies among individuals depending on that person’s immune state. For an infectious bite that leads to an asymptomatic infection or a treatment failure that results in persistent parasitaemia, hosts are transferred into an ‘asymptomatic state’ and their parasitaemia is set to a randomly drawn value with a mean of 1000 parasites per µl of blood (normal distribution with mean 3 and standard deviation = 0.5).

As in our previous publication, we parameterize the duration of infection to malaria therapy data. According to Maire et al [13] and Eyles and Young [14], mean durations of infection were 169, 121 and 222 days for groups of patients untreated or occasionally treated with quinine during clinical episodes. In addition, the World Health Organization’s Garki Project Report [15] indicates an asymptomatic duration between 55 and 500 days and this report shows that older individuals have shorter infection durations due to the development higher immunity.

Using these data as guides for the minimum and maximum length of an asymptomatic infection, the duration of infection in the model is set to have a minimum of 60 days and a maximum of 300 days, for individuals with full immunity and individuals with no immunity, respectively. Note that in our model individuals with naïve immunity will acquire immunity during infection so those individuals will not have 300-day long infections. The expected duration of an asymptomatic infection for an individual starting with no immunity is 281 days for a 1-year old child and 197 days for an 20-year old adult.

Immune system clearance of parasites, for each clone, during the asymptomatic phase occurs according to the following discrete growth model that is evaluated every *K* = 7 days

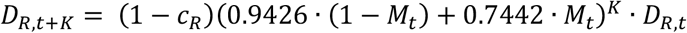

where *D*_*R*_ is the parasitaemia of clone *R* and *M*_*t*_ is the immune level (between 0 and 1) at time *t*. For genotypes with a single mutation the fitness cost above is set to *c*_*R*_ = 0.0005 which will result in an approximate 17% annual fitness cost for genotypes carrying one mutation. For a genotype with *m* mutations, the fitness cost is set to (1 – *c*_*R*_)^*m*^. For some mutations where *in vitro* fitness costs have been measured extensively, we make exceptions, here based on Small-Saunders et al [10], and assign them specific *c*_*R*_ values. For *pfcrt* T93S, we have *c*_*R*,93S_ = 0.0000314. For *pfcrt* F145I, we have *c*_*R*,145I_ = 0.00102. For *pfcrt* I218F, we have *c*_*R*,218F_ = 0.000453. These three values average to *c*_*R*_ = 0.0005 and have relative differences according *in vitro* cost-of-resistance measurements.

### 6 Host Attractiveness to Mosquitoes

Mosquitoes find and select humans for blood meals based on a range of factors including body heat, presence/detection of carbon dioxide, or odor and as a result each person in the simulation is assigned their own ‘attractiveness’ value to mosquitoes. This attractiveness is implemented as relative biting rate, and each host is randomly assigned a biting rate from a truncated Gamma distribution with mean 5.0 and standard deviation 10.0 (truncated at 0.35 on the left and 35.0 on the right) allowing for a maximum 100-fold difference between biting attractiveness between any two individuals. To improve the biting algorithm in the model, a ‘roulette sampling’ system was developed which can rapidly and proportionally sample from one million individuals according to their biting attractiveness. This is described below.

#### 6.1 Roulette sampling

In the update from model version 3.3 to model version 5.0, we switch from multinomial sampling to roulette sampling because every individual now has a relative biting rate that is drawn from a Gamma distribution instead of being grouped into 100 discrete groups as in the previous version. Below is the pseudocode of the roulette sampling implementation.

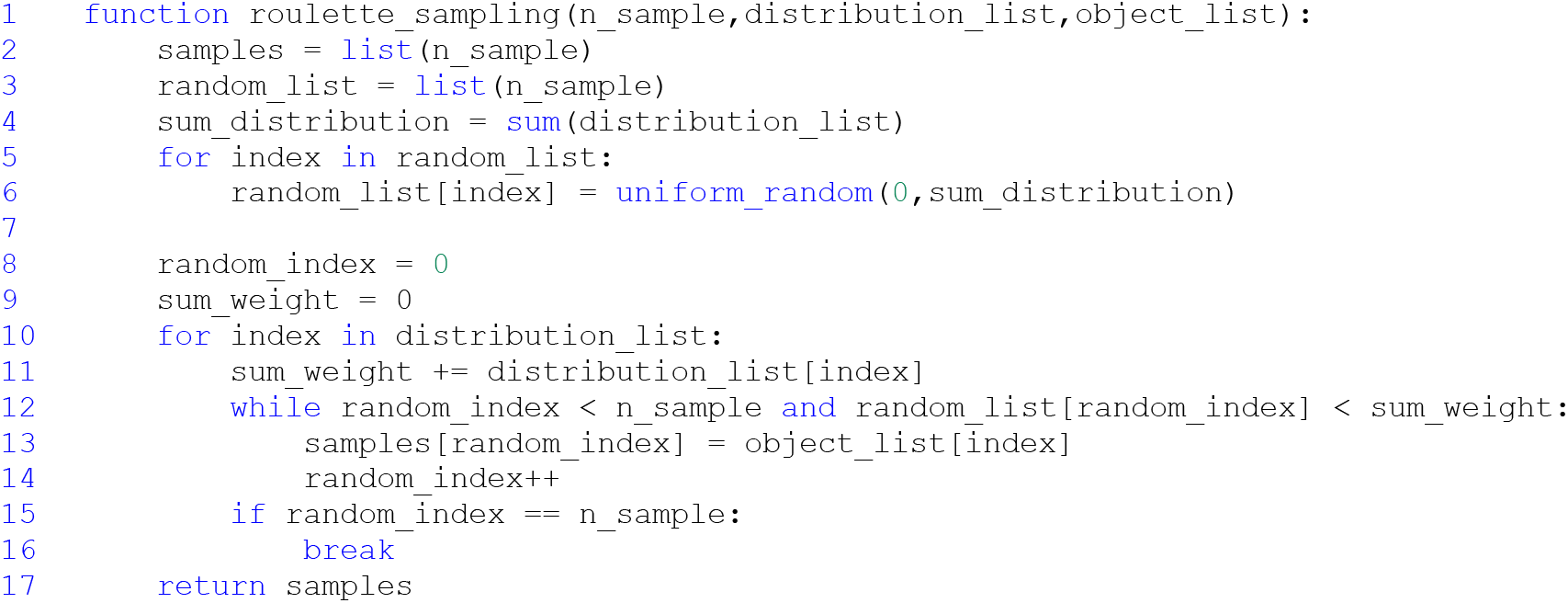

The following example describes how the roulette sampling works in the model. Assume we have a list of six people (this is everyone in the population in this example) and each person has a relative biting rate. We want to select four people from this list based on their biting rate (i.e. attractiveness to mosquitoes). In this selection scheme, selection is done with replacement which means that one individual can be selected and bitten multiple times. A person who has a higher relative biting rate has a higher chance of being selected. The list below contains everyone in the population and it does not need to be shuffled or sorted before beginning the selection algorithm.

|  |  |  |  |  |  |  |
| --- | --- | --- | --- | --- | --- | --- |
| Person index $i$ | 1 | 2 | 3 | 4 | 5 | 6 |
| Biting rate | 0.1 | 0.4 | 0.9 | 0.2 | 0.2 | 0.5 |

First, we compute the sum of all the relative biting rates of six persons, which is 2.3. We will use this to perform random uniform draws on the interval [0, 2.3]. We then create a random list with length four (number of bites, or number of people sampled to be bitten) and each list member is a uniform random draw from 0.0 to 2.3, in ascending order. This list must be sorted.

| Uniformly Drawn Random Numbers<br>("boundary markers" on $[0, 2.3]$) | Bitten<br>Index $j$ | 0 | 1 | 2 | 3 |
| --- | --- | --- | --- | --- | --- |
|  | Value | 0.23 | 0.575 | 0.828 | 1.495 |

Next, we loop through all individuals. In this loop, we keep track of all “biting rate sums”. In other words, the relevant value in each step of the loop is the sum of all biting rates from person 1 to person *i*. If the extension of one “biting rate sum” at loop step *i* to the next “biting rate sum” at loop step *i* + 1 crosses a boundary marker from the sorted list of boundary markers above, person *i* is selected to be bitten. If *k* boundary markers are crossed at this step, person *i* is bitten *k* times. Once the desired number of bites has been reached, the loop stops.

This process has a complexity of *O*(*n* + *m*) where *n* is number of individuals and *m* is number of bites. The running time to sample 2000 individuals out of one million individuals, once per day for 10,000 days, takes around 30-40 seconds.

**Table S3:** Roulette sampling explained step-by-step.

| Random Value<br>(uniformly drawn<br>“boundary marker”) | Sum weight | Steps | Selected person |
| --- | --- | --- | --- |
| 0.23 | 0.1 | $0.1 < 0.23$:<br>- move to person 2<br>- bitten index $j$ is unchanged (0)<br>- add weight of person 2 to sum weight ($0.1 + 0.4 = 0.5$) | |
| 0.23 | 0.5 | $0.5 > 0.23$:<br>- add person 2 to the “bitten” list<br>- increase bitten index $j$ to 1<br>$0.5 < 0.575$:<br>- move to person 3<br>- bitten index $j$ is unchanged (1)<br>- add weight of person 3 to sum weight ($0.5 + 0.9 = 1.4$) | 2 |
| 0.575 | 1.4 | $1.4 > 0.575$:<br>- add person 3 to “bitten” list<br>- increase bitten index $j$ to 2<br>$1.4 > 0.828$:<br>- add person 3 to “bitten” list<br>- increase bitten index $j$ to 3<br>$1.4 < 1.495$:<br>- move to person 4<br>- bitten index $j$ is unchanged (3)<br>- add weight of person 4 to sum weight ($1.4 + 0.2 = 1.6$) | 2, 3, 3 |
| 1.495 | 1.6 | $1.6 > 0.828$:<br>- add person 4 to “bitten” list<br>- increase bitten index $j$ to 4 | 2, 3, 3, 4 |
|  |  | Loop finishes and exits due to random index equaling 4 |  |

**Figure S1:**
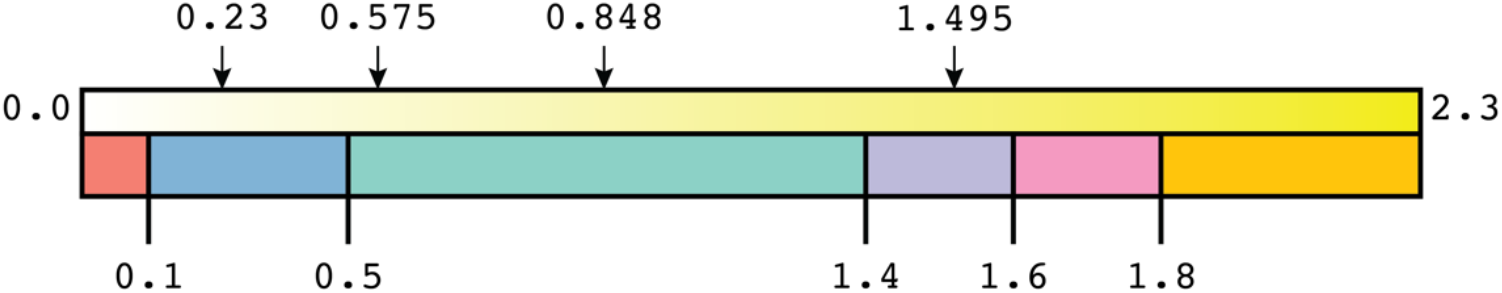
Illustration of roulette sampling. The gradient bar on top shows the full range of uniform draws with the four “boundary markers” shown as the four random variates that were drawn. Color blocks at bottom correspond to individuals that have different biting rates (length of block is proportional to the biting rate). Every arrow pointing inside a particular ‘person block’ corresponds to one bit on that person.

The final list of selected people (i.e. people to be bitten) will be:

|  |  |  |  |  |
| --- | --- | --- | --- | --- |
| Selected individuals to be bitten | 2 | 3 | 3 | 4 |
| Individual’s Biting rate | 0.4 | 0.9 | 0.9 | 0.2 |

### 7 Host Infectiousness to Mosquitoes

When a mosquito bites an infected person, there is a chance that the mosquito will take up gametocytes and become infected. A function from asexual parasite density to probability of successful infection was developed by Ross et al [8] and re-described in our previous model (version 3.3 – see Nguyen et al [12] pp.6-7 of supplement). This same function, shown below, is used in version 5 of our model.

**Figure S2:**
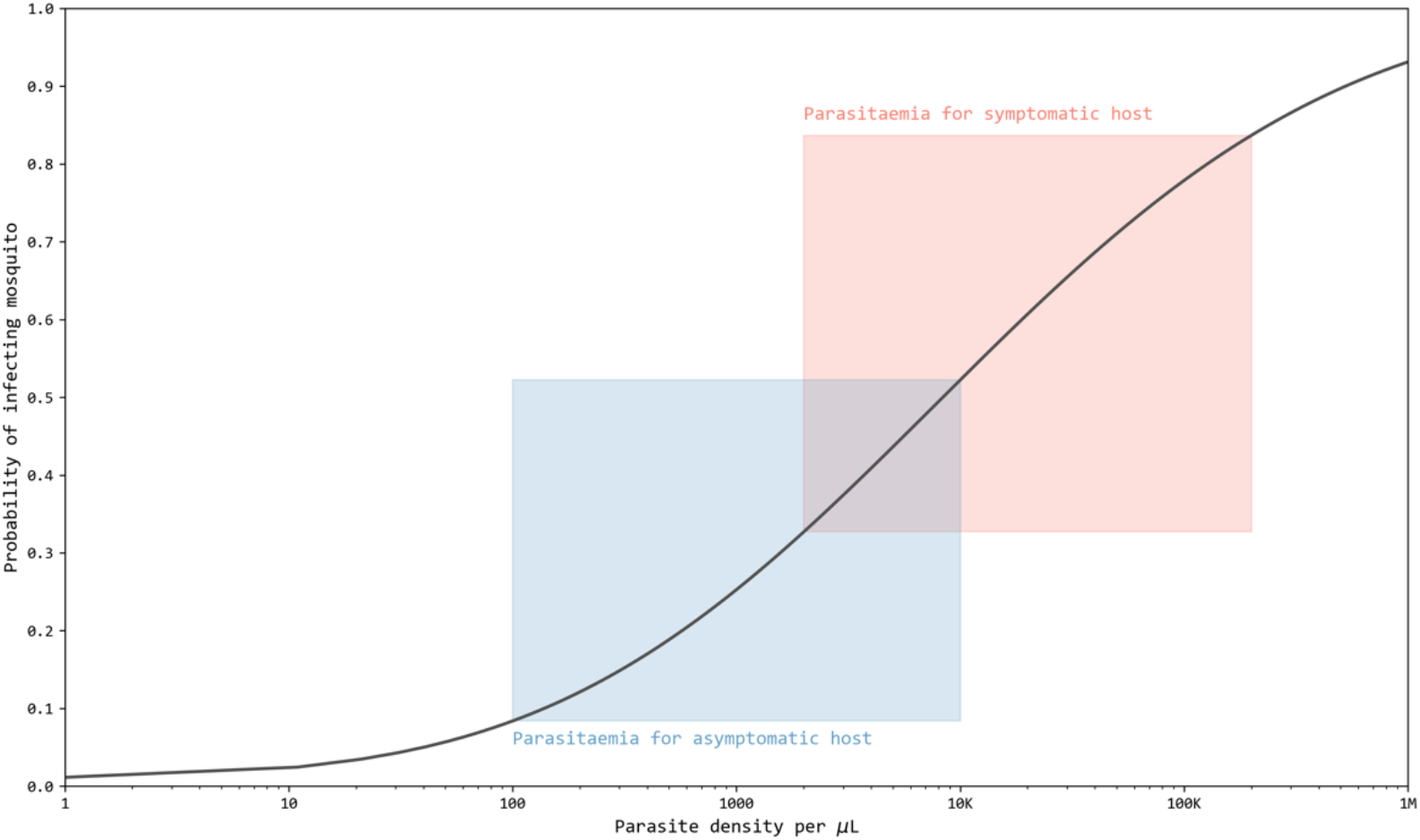
Relationship showing probability of infecting a mosquito during a single feed as a function of asexual parasite density (which is assumed to be proportional to gametocyte density during most stages of an infection). Ranges for parasitaemia and infection probability for asymptomatic and symptomatic hosts are shown in blue and red (respectively).

### 8 Mosquito Biting Model

In sections 2 through 5, we defined the *P. falciparum* genotypes used in the simulation, the multi-clonal structure of infection, and the parasitaemia level for each genotype including how it is set, increased, and decreased. In sections 6 and 7 the level of biting on individuals and whether this biting leads to infected mosquitoes is described. Here in section 8, we describe a major new model component describing how mosquitoes take parasites (sometimes multiple parasites with different genotypes) up from infected hosts and how these parasites recombine within the mosquito.

#### 8.1 Post-recombination mosquito cohort (PRMC)

First, each day, we create a new 500-mosquito cohort of infected mosquitoes that is meant to hold a representative sample of currently circulating falciparum genotypes. Cohort history is tracked for 11 days so that human hosts infected today are bitten by mosquitoes that fed 11 days ago. A daily cohort is created by simulating 500 bites on human hosts – sampled via roulette sampling according to the product of biting attractiveness and host infectiousness to mosquitoes. These 500 bites on human hosts may be interrupted feeds on pairs of hosts or single feeds on single hosts. These bites may take up multiple gametocytes with different genotypes. Random union of gametes followed by random segregation of chromosomes (i.e. traditional recombination for sexually reproducing organisms) occurs in the model for the gametocytes the mosquito has taken up. No recombination occurs within chromosomes in this version. Then, one parasite haplotype post-recombination is assigned to the subsequent population of ookinete, oocysts, and sporozoites that will be able to infect new human hosts 11 days later.

When a mosquito takes a blood meal, it will take either a full blood meal on one host or be interrupted by the human reaction to the bite and it may then complete its meal on a second host. This interrupted feeding probability (i.e. the probability that a fed mosquito has human blood from two distinct humans) has been measured to be as high as 20% in some contexts [16–25], and we vary this probability in our current analysis. The probability of mosquitoes taking interrupted feeds is used in the biting model whose details are described in below in the diagram below.

**Figure S3:**
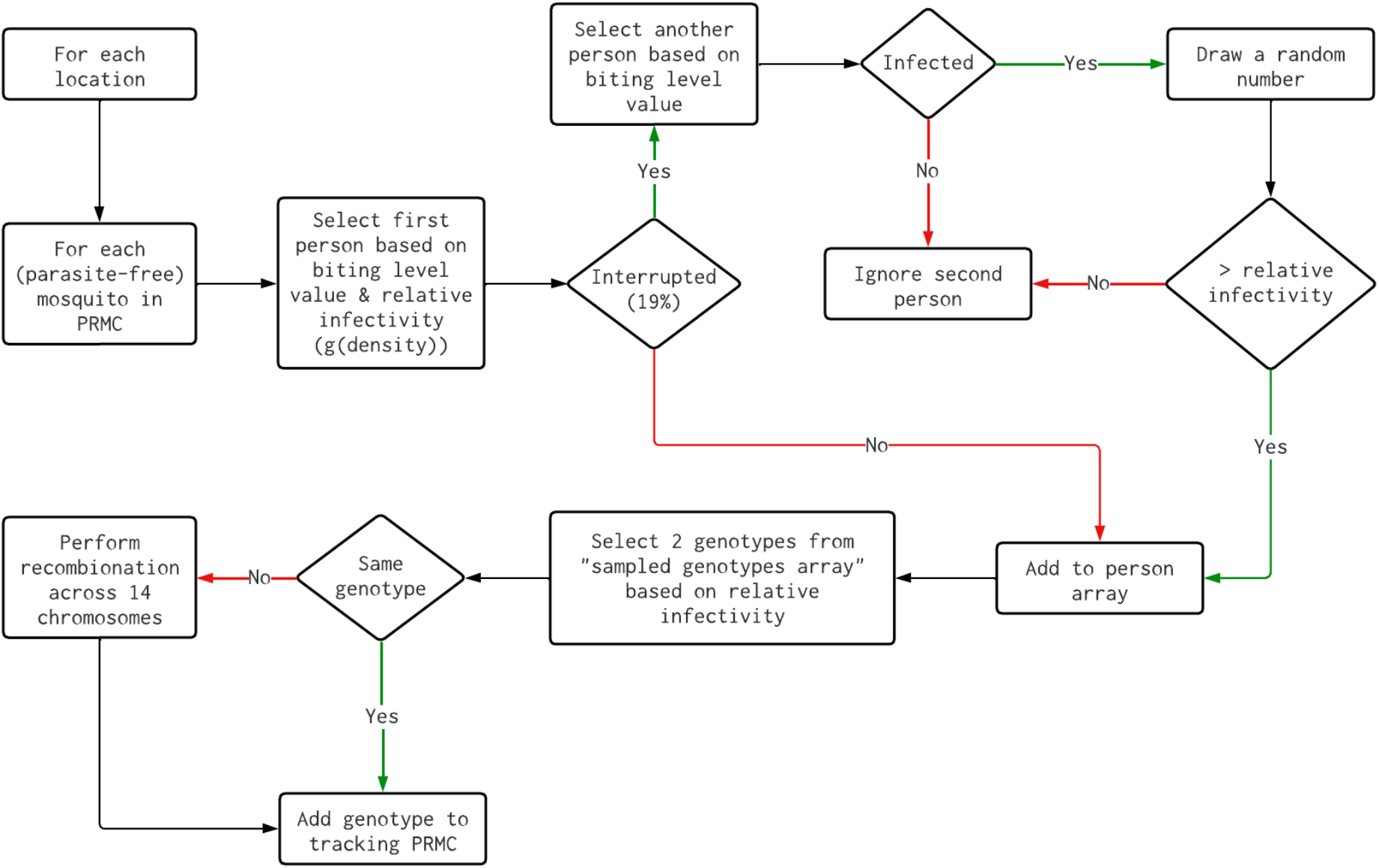
The flow diagram of how each mosquito in the cohort (the PRMC) is assigned a particular genotype after recombination has taken place. Selecting genotypes from the ‘sampled genotypes array’ is done with replacement.

#### 8.2 Total force of infection

In the previous model version 3.3, the force of infection (FOI) of each genotype (accounting for recombination with a mating-table approach) was used to compute the number of bites of that genotype. In the current version 5, the FOI is generated based on the total number of parasites in the population across all genotypes. The equation below defines the total force of infection Λ_*t*_ at time *t* in the model.

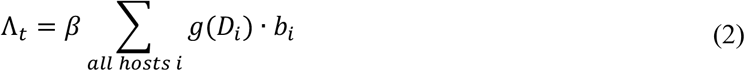

where (*i*) the overall transmission parameter *β* is calibrated in all model runs to achieve a particular malaria prevalence level, (*ii*) the infectivity-to-mosquitoes function *g*, which depends on parasite density *D*_*i*_, is described in section 7, and (*iii*) the biting levels or human biting attractiveness parameters *b*_*i*_ are described in section 6. The number of new infectious bites on a particular day *t* in a particular location is sampled from a Poisson distribution with mean set to Λ_*t*−1_.

When an individual is bitten and infected, a new *P. falciparum* genotype is chosen randomly (uniformly) from the 500 mosquitoes in the PRMC. An 11-day history of the PRMC is tracked, so today’s infections will have their genotypes sampled from the PRMC created 11 days ago.

#### 8.3 Validation of recombination mechanism using basic measures of linkage disequilibrium

Linkage disequilibrium (LD) is the non-random association of alleles at different loci in a population. For alleles *A* and *a* at one locus and alleles *B* and *b* at a second locus, we use the following simplified equation to describe the LD between *A* and *B:*

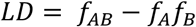

without any normalization typically used in LD equations. The parameter *f* simply denotes the allele frequency of A or B, or the genotype frequency of AB. We do this simply to verify that LD approaches zero in situations with no selection, no mutation, and no population structure. We simulate a population with two parasite genotypes (K76 C580 and 76T 580Y) at equal 0.50 genotype frequencies as the starting point with no drug treatment, no cost of resistance for mutant alleles, and no mutation allowed in these runs. In all situations tested (*N* = 1600 simulations in total) the LD between these loci approached zero and the four genotype frequencies reach equilibrium frequencies of 0.25.

**Figure S4:**
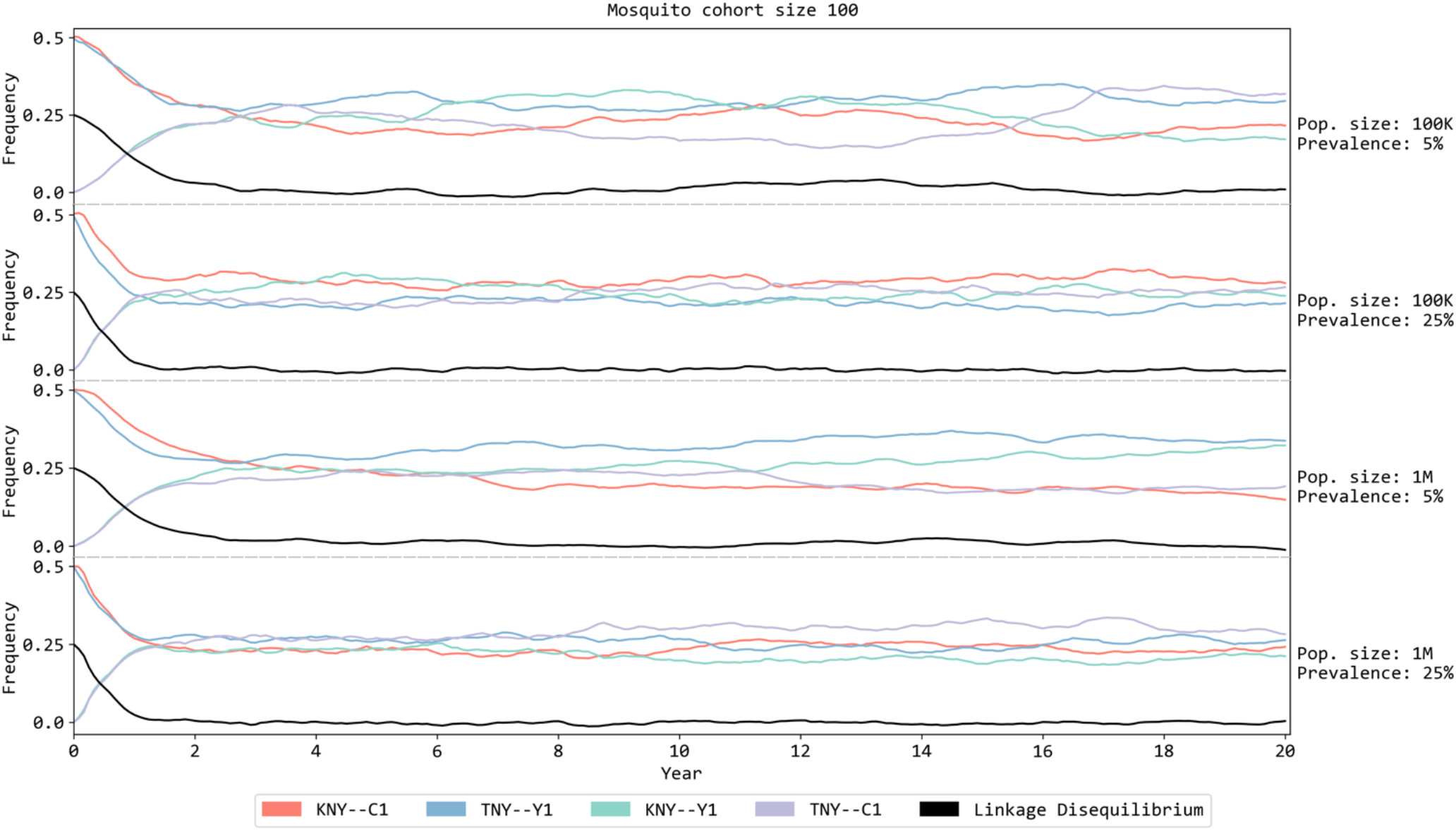
The linkage disequilibrium (LD) output of the model with two population sizes in two prevalence settings. The *x*-axis indicates the number of years run while *y*-axis shows the frequency of the genotypes and LD. In this setting, two initial genotypes (KNY—C1 in pink color and TNY—Y1 in cyan color) are seeded into the population (i.e. the two distinct genotypes are K—C and T—Y). No treatment is applied in this setting and the mosquito interrupted feeding rate is set to 99% to accelerate recombination. The expected result is four genotypes distributed with approximately 0.25 genotype frequency each and with LD approaching or close to zero.

### 9 Treatment Seeking Model

In our model, the probability that an individual seeks treatment is called treatment coverage and this probability is different for individuals aged under 5 and over 5. One treatment coverage is set when running the model in the burn-in phase (while the model approaches equilibrium) and another treatment coverage is set when treatment strategies begin to be evaluated. Increases in treatment coverage can be modeled during the treatment strategy period. Individuals can choose to seek treatment in the public or private market, and data for this are typically obtained from Demographic Health Surveys (DHS). Depending on the health-care setting individuals can receive the recommended first-line therapy or one of a number of therapies/drug sold in private clinics or pharmacies (again, parameterized using data from DHS surveys).

### 10 Mass Drug Administration Model

In malaria interventions, mass drug administration (MDA) is sometimes deployed to give treatment to everyone sick or healthy, infected or uninfected – in a particular geographical area. Since everyone is receiving treatment, malaria transmission among individuals is reduced and lowers the number of new infections in the population, sometimes to zero. This strategy is a good option to eliminate malaria in low-endemicity regions (with prevalence < 1.0%) where everyone has good access to treatment, vector control, and routine diagnosis and surveillance. In our model, the number of people that receive treatment during MDA is determined by individuals’ variability in participating (i.e. showing up) in the MDA. Each individual is assigned a participation probability drawn from a β-distribution with standard deviation set to 0.3; see Nguyen et al [2]. In the analysis in the present manuscript, MDA scenarios were not evaluated.

### 11 Adaptive Multiple First-line Therapies

Deployment of multiple first-line therapies (MFT) is a strategy where different ACTs (or in general, different therapies) are distributed simultaneously in a population to provide a variable drug environment for the purpose of slowing the evolutionary/adaptive process of malaria parasites [26]. This strategy has the advantage of both delaying the emergence of resistant parasites and reducing the number of treatment failures, according to both mathematical and agent-based models [12,26].

In version 3.3, MFT is implemented with an equal distribution of all ACTs. In this version, we also introduce an “adaptive MFT” approach in which the distribution of a drug into a population is adjusted based on the number of treatment failures of that drug. Initially, for three therapies, the distribution of each drug is set at 33.3%. After a specified duration (2 years), the fractional distribution of each therapy (denoted here by *µ*_*i*_) is adjusted for each drug based on its most recent 60-day treatment failure rate *F*_*i*_:

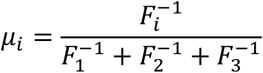

where *F*_*i*_ is replaced with 5% if the failure rate is below 5% (all others are then rescaled or renormalized to maintain their relative proportions). The new distributions are put in place after a delay of one year. This adjustment will use drugs in inverse proportion to their failure rates. In other words, if therapy *a* has three times the failure rate of therapy *b*, therapy *a* will be used for one-third as many malaria cases as therapy *b*.

**Figure S5:**
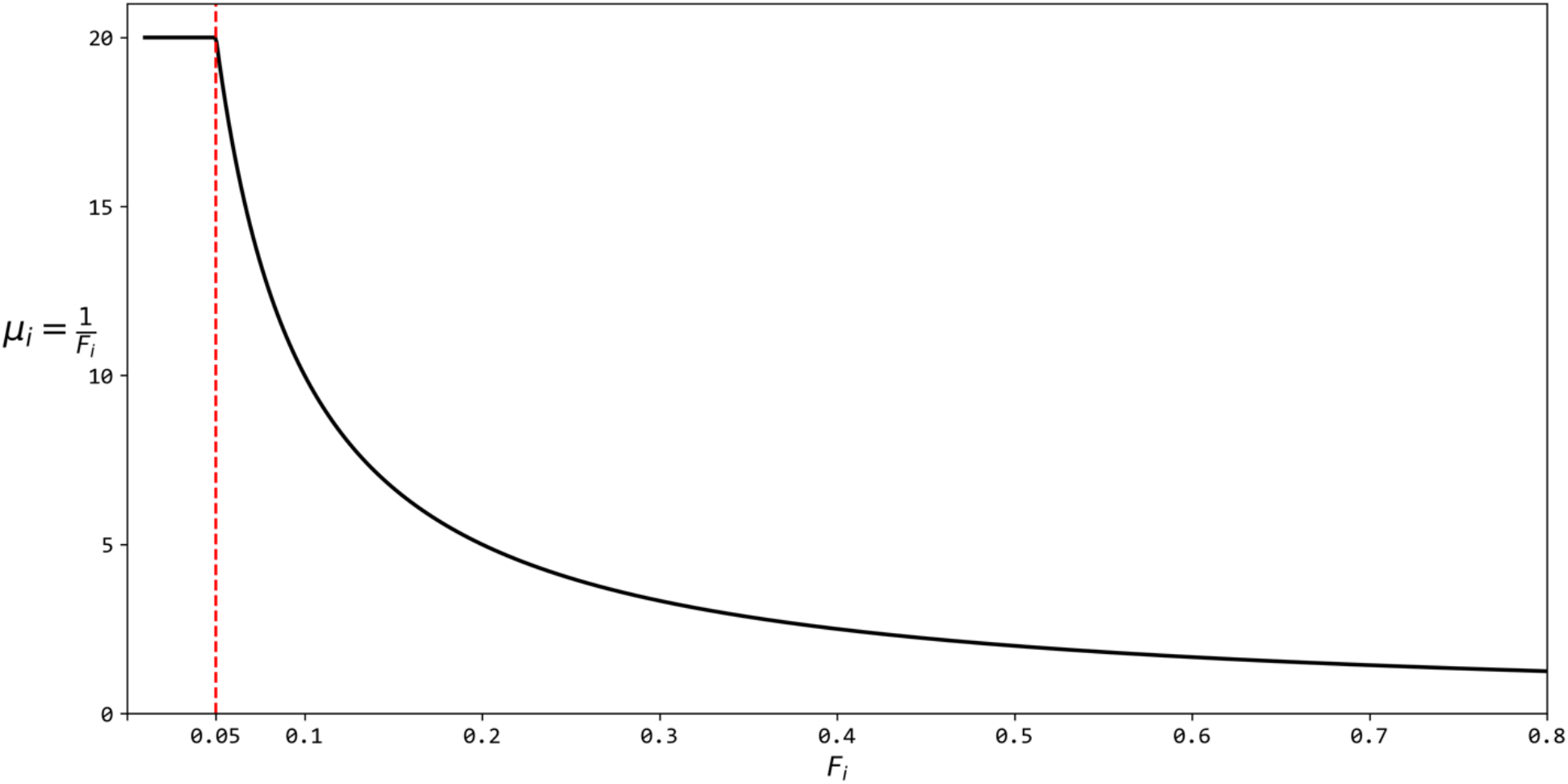
The relation between treatment failure of a drug (*F*_*i*_ on *x*-axis) and its new distribution (*µ*_*i*_, on y-axis) in adaptive MFT method.

### 12 Pharmacokinetics

In this version, a single compartment pharmacokinetic model is used, and drug half-life data are the same as in version 3.3 with following values used:

**Table S4:** Half-lives of monotherapies. Source in Nguyen et al [12].

| Drug | Half-life (t*) |
| --- | --- |
| Artemisinin derivatives | 4-11 hours (DHA: 11-12 hours) |
| Lumefantrine | 4.5 days |
| Amodiaquine | 9 days |
| Piperaquine | 28 days |
| Mefloquine | 21 days |
| Chloroquine | 10 days |
| Ganaplacide (KAF156) | 1.6 days |

### 13 Pharmacodynamics and Treatment Failure Model

In Nguyen et al [12], the pharmacokinetics model is a modified version of the standard Michaelis-Menton equation (or Hill equation)

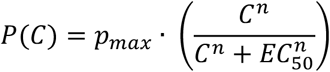

with *C* as the drug concentration in the patient’s blood (central compartment), and *p*_*max*_ as the daily fraction of parasites that are removed (killed) by a high drug concentration drug. The parameter *EC50* is the concentration level corresponding to 50% killing effect, and *n* determines how steep the concentration-effect curve is. The parasite density at day *t* is:

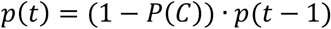

Since *EC50* determines the percent of parasites killed by a certain concentration of drug and since parasites can mutate to a higher-survivorship phenotype to that drug, we associate each genotype with an *EC50* value for a particular drug. In the previous version of the model (v3.3), there were 64 genotypes in total and the *EC50* values were computed (approximated) in advance as a lookup table of *EC50* values associated to genotypes.

In the new version of the model (v5) with the loci described in section 2, the number of possible genotypes has increased to 2^25^ (about 32 million) and the *EC50* values of new genotypes must now be calculated on the fly to save memory and computation time. This is done in two-step process:

**Step 1**. When a mutation occurs, a new genotype is formed that differs by one allele (normally) from a previously existing genotype. Considering all the mutant alleles (or derived alleles) and all drugs in the simulation, each mutant-drug combination is assigned an “EC50 shift factor” (parameter *e*) that shifts the EC50 value to the right from EC50 to *e* · EC50 generating a classic rightward-shifted dose response curve. The parameter *e* is denoted by multiplicative_effect_on_EC50 in the simulation’s input file, and it is best written as *e*_art,R561H_ to specify the increase in resistance associated with a particular allele (here 561H) when encountering a particular antimalarial compound (here artemisinin).

The mutant-drug combination is simply a genotype-environment combination. An increase in EC50 increases survivorship for a genotype in a particular environment, and the *e*-parameters therefore can be viewed as factors influencing each genotype’s Darwinian fitness, although the exact relative fitnesses in these comparisons are not 1.0 and *e*. When multiple mutations occur, the new EC50 is calculated simply as *e*_1_ · *e*_2_ · *e*_3_ · EC50 assuming a level of independence among these mutations’ three fitness effects. In other words, as assumption of non-epistasis is made in the initial calibration of the fitness effects across genotypes. We simply have single-locus effects in given drug environments. These single-allele effects are shown in the table below.

**Table S5:**
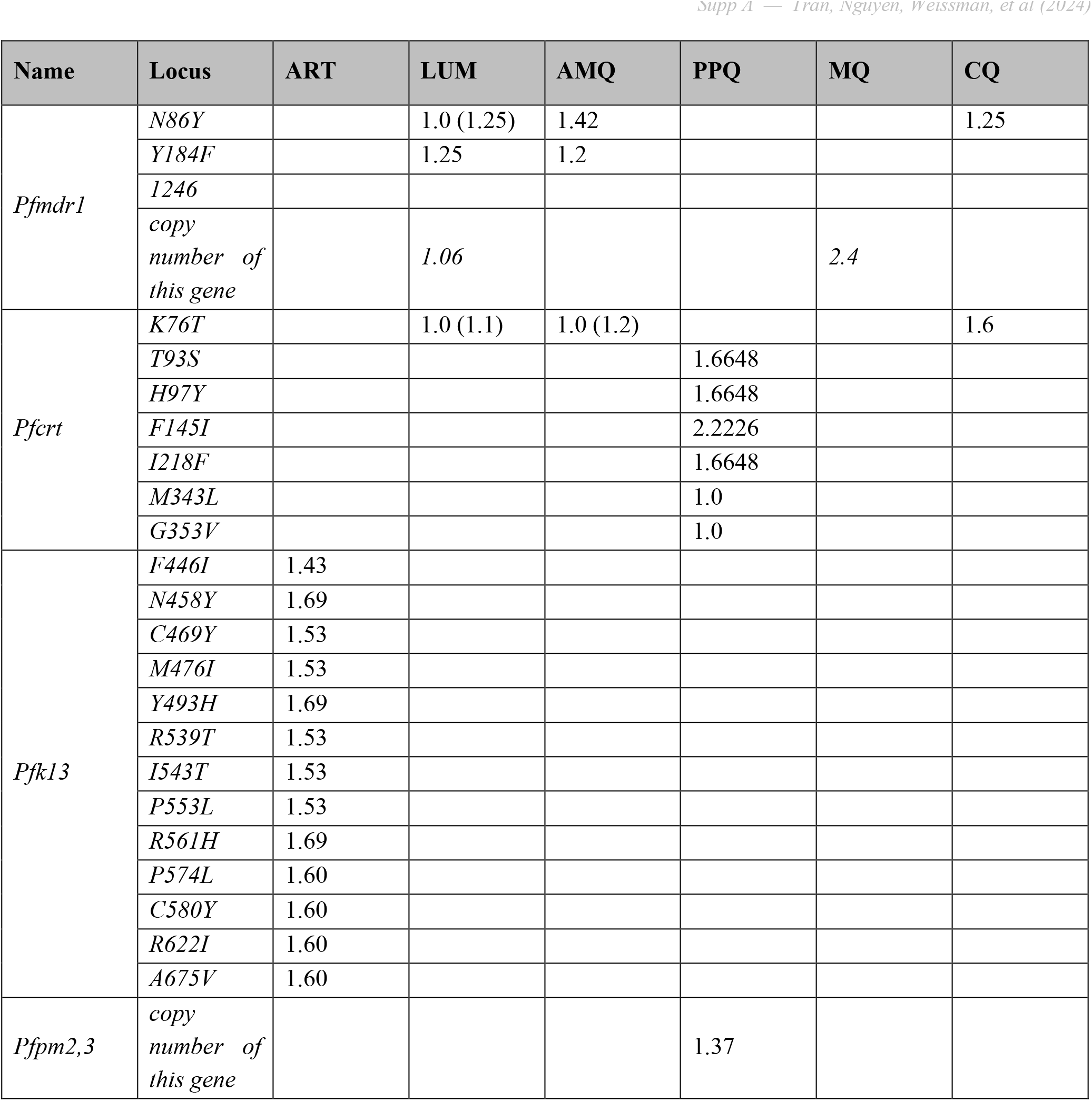
Multiplicative effect that each mutant allele has on each drug’s efficacy. Number shown in each cell is the *e-*value for that mutation’s effect on the EC50 to a particular drug; when no value is shown this means there is no effect (*e* = 1.0). In some cases, a mutation will lower an EC50 value, e.g. when 76T mutates to K76 the EC50 for lumefantrine increases. In these cases, the mutant’s EC50 is shown as with an *e*-value of 1.0, and the original *e*-value for the wild-type allele is shown in parentheses.

As an example, consider the following alleles in the *pfcrt* gene (chromosome 7) that influence piperaquine resistance:

**Table S6:** Multiplicative effects (bottom row) on EC50 of six different mutations in *pfcrt* associated with PPQ resistance.

|  |  |  |  |  |  |  |
| --- | --- | --- | --- | --- | --- | --- |
| amino-acid position in *pfcr*t: | 93 | 97 | 145 | 218 | 343 | 353 |
| Allele (wild type) | T | H | F | I | M | G |
| Allele (mutant) | S | Y | I | F | L | V |
| Piperaquine Base EC50 | 0.58 |  |  |  |  |  |
| EC50 factor (wild type) | 1.0 |  |  |  |  |  |
| EC50 factor (mutant) | 1.66 | 1.66 | 2.22 | 1.66 | 0.99 | 0.99 |

The EC50 value for piperaquine’s action on the wild-type genotype THFIMG is 0.58 × 1.0, the EC50 value for PPQ on the 1-point mutant THIIMG is 0.58 × 2.22, and the EC50 for PPQ on the 2-point mutant T<u>YI</u>IMG is equal to 0.58 × 1.66 × 2.22.

This summarizes the process of approximating EC50 values across all loci in the current simulation when multiple resistance mutations are present.

**Step 2**. For some genotypes, fitness values (in the form of treatment failure rates) will have been measured in clinical trials or therapeutic efficacy studies (TESs). These measured or approximated genotype-specific treatment failure rates can be seen in Supplement 2 of Nguyen et al [2]. When a genotype-specific approximation for an EC50 or a treatment failure rate is available, this value will override the calculated value in Step 1 above.

In steps 1 and 2 above, it is assumed that base EC50 values exist for the wild-type genotype. These base EC50 values are calculated using well-known treatment efficacies on wild-type parasites. Since the method of updating EC50 values has changed (from v3.3 to v5) due to the additional genotypes to the model, the efficacy of each drug needs to be calibrated again, especially for piperaquine and piperaquine-resistant genotypes in *P. falciparum* chromosome 7. Some of this calibration process is described below.

#### 13.1 Example calibration of piperaquine-resistant alleles in pfcrt gene

According to a number of in vitro studies [7,9,10,27] and in vivo studies [28], six amino acids in *pfcrt* gene are confirmed to confer the resistance to piperaquine: T93S, H97Y, I145F, F218I, M343L and G353V. Four of these alleles had in vivo efficacies measured as part of the TRACII study in SE Asia (during 2015-2018 [28]). The EC50 values for the T93S, H97Y, F145I, and I218F alleles were estimated as follows.

Out of 140 patients that received DHA-PPQ in this study, 119 had their *pfcrt* alleles characterized and 31 had none of the four alleles listed above. The study population as a whole was characterized as 74% with double-copy plasmepsin and the 580Y allele, 17% with single-copy plasmepsin and the 580Y allele, and 9% with single-copy plasmepsin and wild-type C580. A simulated clinical trial population was created with these genotype proportions (74, 17, 9) and EC50 value for the effect of multi-copy plasmepsin on PPQ activity was inferred by matching the simulated efficacy to the measured efficacy on these 31 patients (87.0% efficacy). See top row of Figure S6.

For each patient subgroup, stratified by genotype (see Figure 1B in van der Pluijm et al [28]), we re-ran a simulated clinical trial with the above population genotype distribution (74, 17, 9) to match the measured DHA-PPQ efficacies on patients carrying one of the 93S, 97Y, 145I, or 218F alleles. In the trial, patients carrying 93S, 97Y, and 218F had an estimated 28-day treatment efficacy of 50.7% and patients carrying the 145I allele had an estimated treatment efficacy of 39.8%.

The mutations 343L, 353V are currently modeled as having no effect on piperaquine resistance because of the lack of in vivo data.

**Figure S6:**
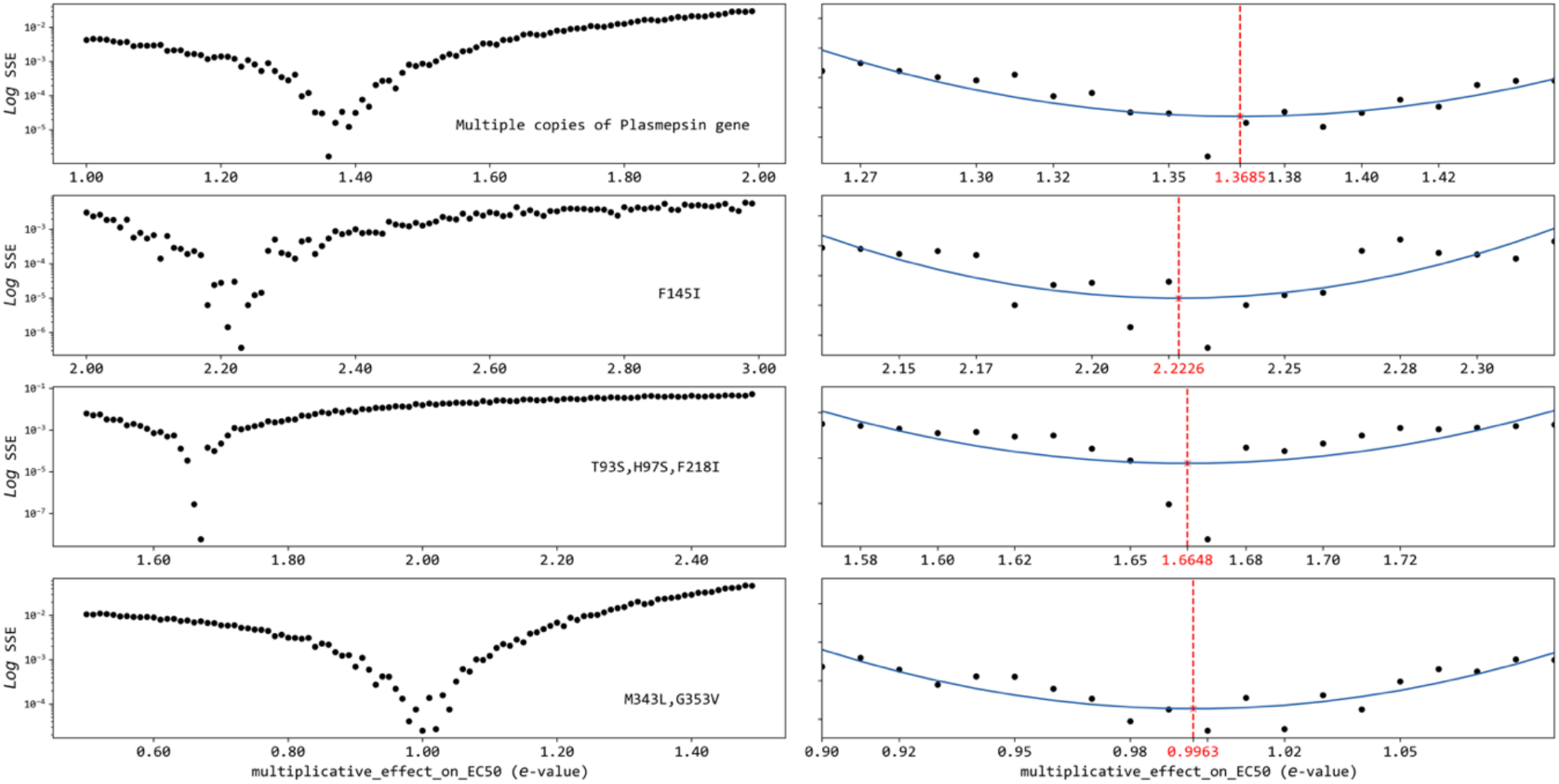
Log likelihood fitting the *e*-value to get the DHA-PPQ efficacy of the model to match clinical data. Each panel row presents a likelihood-fit of the *e*-value using sum of squared error and the left panels presents the errors of fittings while the right panels show the optimum *e*-value. The *x*-axis shows the *e*-values of each allele in the *pfcrt* gene or copy-number variation of the plasmepsin gene. The *y*-axis presents the sum of squared error in log10-units. Each black dot shows the sum of squared error between DHA-PPQ efficacy in the simulated data with a specific *e*-value (summed across 10,000 patients) and the clinical data. The blue line indicates the quadratic fitting of 20 data points around the lowest *e*-value on the left panel. The value in red indicates the maximum likelihood *e*-value.

#### 13.2. Calibration of EC50 for kelch13 alleles

To calibrate the EC50 of various *pfkelch13* alleles, we used parasite clearance half-life (PCT_1/2_) estimates from a WWARN meta-analysis [29] to estimate the parasite density after one day with artemisinin monotherapy (assuming the partner drug had little effect during this time). From Figure 4 in this meta-analysis, the mean clearance half-lives of all *pfkelch13* alleles in Table S5 can be grouped into three groups shown in Table S7 below.

**Table S7:** List of *pfkelch13* alleles, their parasite clearance half-lives (PCT_1/2_), and the calculated mean parasite density 24 hours after presentation based on this PCT_1/2_. It is assumed that mean parasite density at presentation is 10^4.301^ or 20,000 parasites per microliter. The parasite clearance half-life is shown in hours.

| Allele | Range of $PCT_{1/2}$ | expected log<sub>10</sub>-parasite density after 24 hours |
| --- | --- | --- |
| F446I | 5.389 | 2.961 |
| R539T/I543T/P553L/C469Y/M476I | 5.87 – 6.43 | 3.122 |
| P574L/C580Y/A675V | 6.61 – 6.71 | 3.218 |
| N458Y/Y493H/R561H | 7.21 – 7.52 | 3.323 |

In our model the mean log-parasite density across 10,000 symptomatic patients with 580Y parasites is 10^4.301^ or 20,000 parasites per microliter (at presentation) and after one day of using artemisinin monotherapy the parasite density should be reduced (according to the half-lives above) to a level between 10^2.9^ and 10^3.4^ (i.e. between 900 and 2200 parasites per microliter). This would be consistent with a parasite clearance half-lives in the range of 5.8 to 7.5 hours for mutant *pfkelch13* genotypes. Our model achieves an approximate version of this behavior (roughly a 12-hour half-life) as it is the 28-day efficacy of 580Y parasites, and not their half-life, that is calibrated to data in our model.

Reduction in PRR (parasite reduction ratios) in the model is done by increasing the EC50 parameter (for artemisinin) which reduces the model-calculated PRR values of artemisinin monotherapy on mutant *pfkelch13* genotypes. We increase artemisinin’s EC50 value even though pharmacokinetics of artemisinin are not modeled explicitly; most of the pharmacokinetic and pharmacokinetic model parameters do not have perfect correspondence with published PK/PD data. Variation (among patients) in PRR values is used to calibrate a 28-day efficacy of 25.5% for three days of artemisinin monotherapy on an uncomplicated falciparum infection with a 580Y genotype. This ∼25% efficacy was chosen based on genotype-stratified trial data from SE Asia last decade (see p11 in supplement 2 of Nguyen et al [2]). In a model-simulated clinical trial of patients infected with 580Y parasites and treated with three days of artemisinin monotherapy, 23% of patients have a daily PRR > 10 and 16% of patients have a daily PRR > 50, and it is this variation in the model that results in some patients being treated successfully while other patients fail treatment.

For alleles 446I, 539T, and 561H, the calibration is done differently. The mean daily PRR-values (calculated from PCT_1/2_ values in Table S7) for these three alleles are 21.9, 15.1, and 9.5, respectively. For these PRRs, the expected log-parasitaemia values after 24 hours are shown in Table S7; these values are adjusted up by a factor of 1.149, which is the exact difference between the expected log-parasitaemia at 24H for a 580Y infection based on PCT_1/2_ and the expected log-parasitaemia at 24H for a 580Y infection based a 28-day efficacy of 25%. This adjustment of 1.149 is included in the sum-of-squares calculation in Figure S7. The modeled 28-day efficacies of artemisinin monotherapy on these 446I, 539T, and 561H are 39.7%, 30.5%, and 20.3%, respectively.

**Figure S7:**
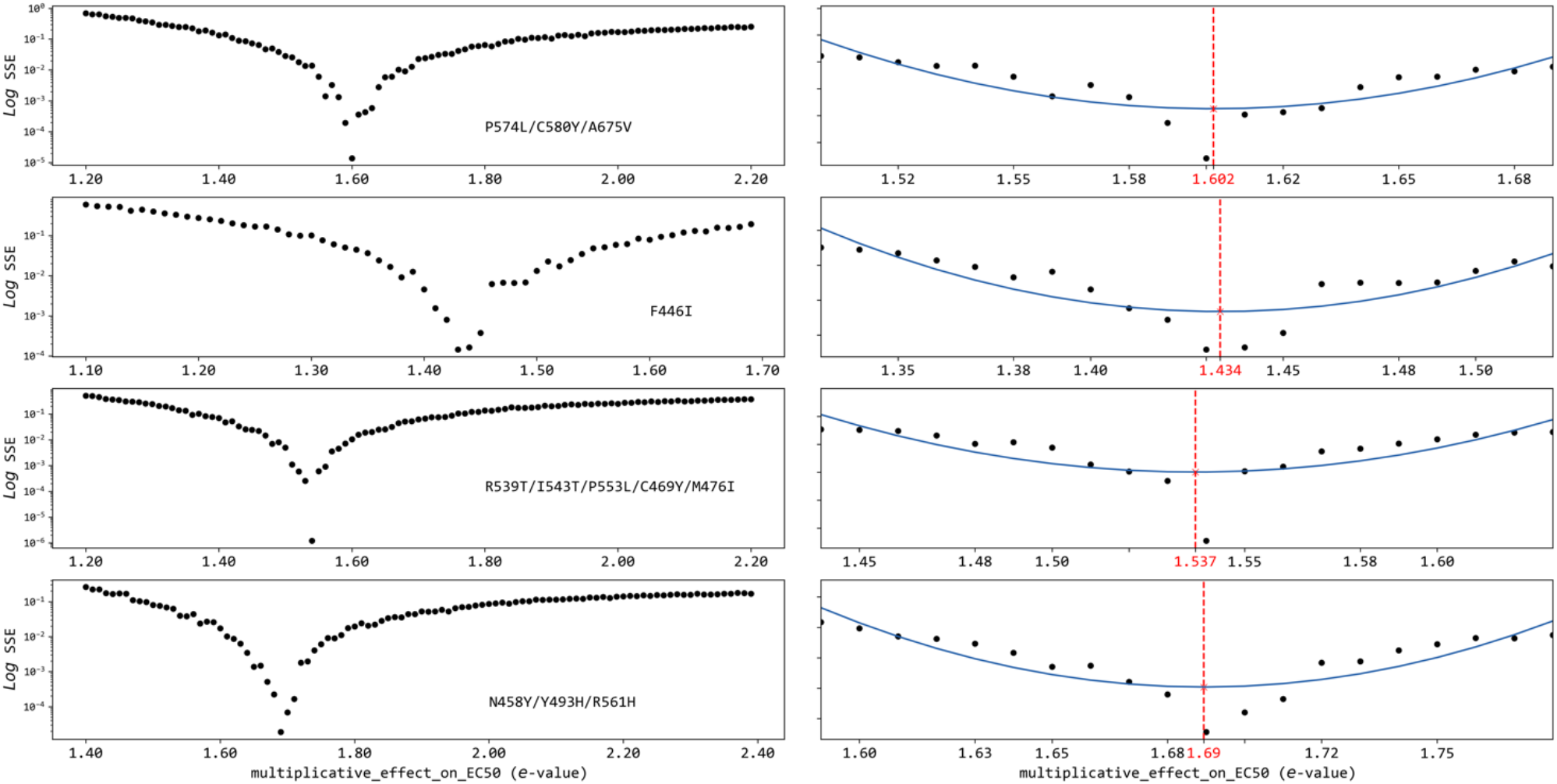
Log-likelihood fitting of the *e*-value based on modeled parasite density on day 2, from which a parasite clearance half-life is calculated; likelihood comparison done to match PC_1/2_ values in WWARN meta-analysis [29]. Each panel row presents a likelihood fit of an *e*-value using sum of squared errors; the left panels show all errors while the right panels show a zoomed-in version and the maximum-likelihood *e*-value. The *x*-axis shows the *e*-values of each allele in *pfkelch13* gene that confers resistance to artemisinin. The *y*-axis presents the sum of squared error in log10-units. Each black dot shows the sum of squared errors between parasitaemia on day 2 of the simulation (across 10,000 simulated patients) and the inferred parasite density from clearance half-life in the WWARN meta-analysis. The blue line shows a quadratic fit of 20 data points around the lowest *e*-value on the left panel. The value in red on the *x*-axis indicates the optimal *e*-value that matches parasite density on day 2.

### 14 Host Age Structure

People with different ages respond to malaria infections differently due to their prior malaria exposure history, current level of immunity, and acquisition rate of immunity. Age is tracked in the model for all individuals. Individuals are born in the model on specific days and are assigned birthdays. Individuals instantiated before burn-in are given random birthdays according to the expected population age structure.

For computational efficiency, the individuals are grouped into 21 age groups, able to be indexed in a particular array structure of the model for faster searching and reporting. In these 21 age groups, individuals younger than 15 are grouped into 15 groups based on one-year age bands. Individuals older than 15 and younger than 65 are grouped into 5 age groups: 15 to 24, 25 to 34, 35 to 44, 45 to 54 and 55 to 64. The last age group is individuals older than 65. The exact percentage of the population in each age group can be found in section 7 in supplement of Nguyen et al paper [12], taken from the Tanzania population report from the Tanzanian National Bureau of Statistics. These age groups are taken as representative, and the generic model setup is not meant to replicate epidemiological conditions of malaria in Tanzania (unless specifically calibrated).

### 15 All-cause mortality and Malaria Mortality by Age

The model’s all-cause and malaria mortality has not changed since the 2015 Nguyen et al paper [12] (section 6 of supplement) and is based on several large but older studies on malaria morbidity, severity, and mortality [30–33]. Briefly, for an untreated symptomatic malaria case, the model’s mortality rate is (1) 4.0% for patients 12 months of age or younger, (2) 2.0% for age group 1-4, or 12 months to 59 months of age, (3) 0.4% for age group 5-10, and (4) 0.1% for individuals 11 or older. These mortality rates account for the fact that a certain percentage of untreated malaria cases progress to severe malaria, but our simulation does not model the symptoms progression from uncomplicated to severe. For treatment failures, the age-specific mortality rates are the same as for the untreated cases shown above. For treatments that result in adequate clinical and parasitological response (ACPR) the malaria mortality rate is zero for all ages.

### 16 Probability that an Infectious Bite Causes an infection in a Human

When an infectious mosquito bites a human, there is a chance that the mosquito successfully infects that person with sporozoites and that the sporozoites develop in the liver and eventually progress to be released into the blood as merozoites. During simulation, only 10% of individuals that are bitten each day are selected to have a successfully established blood-stage infection. This probability is 10% in both versions 3.3 and 5 of the simulation. This probability affects the EIR values reported in the model, as EIR counts all bites by infectious mosquitoes even ones that do not cause established blood-stage infections.

### 17 Model of Immune Acquisition and Immune Waning

As in Nguyen et al [12] (supplement, section 10) the acquisition of immunity of an individual harboring blood-stage parasites in the bloodstream is defined by the following equation, where *M*(*t*_1_) represents the a person’s current “immunity level” on a scale of zero to one:

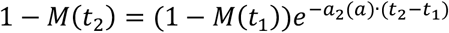

Above, *a*_2_ (*a*) is the age-dependent rate of immune acquisition (see below), and *t*_2_ − *t*_1_ is the time interval (in days) between assessments. In the absence of infection, an individual’s immunity wanes over time, as governed by the equation:

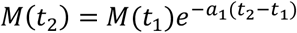

where *a*_1_ (*t*_2_ − *t*_1_) is the rate of immune decay during the time *t*_2_ − *t*_1_. We set the value *a*_1_ = 0.0025 corresponding to 90% immune loss after 2.5 years [34]. To account for age-dependent differences in immunity acquisition, we applied two different equations for individuals below/above 10 years of age. For those 10 and under, the daily immune acquisition rate *a*_2_ is:

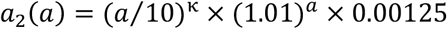

where the parameter *κ* controls the curvature of age-specific immune acquisition dynamics. A lower *κ* leads to similar immune acquisition rates between children and adults (see Figure S8). Conversely, higher values of *κ* result in children experiencing much slower rates of immune acquisition than adults. For individuals over the age of ten, the daily immune acquisition rate is set to

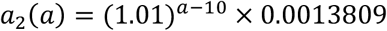

indicating that “full immunity” or “high immunity” to malaria is acquired by adults or older children in about one or two years. The parameter *κ* is calibrated (see section below) to match field data on age-specific malaria symptoms presentation.

### 18 Immune-mediated Symptoms Model

The probability of progressing to symptoms, denoted as *P*_*clin*_, is formulated as a function of the immunity level *M* according to the equation:

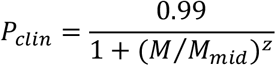

Here, the parameter *z* characterizes the relationship between immunity level and the likelihood of symptoms development. To explore its impact on clinical malaria patterns, *z* is varied between 2.0 and 8.0. We set *M*_*mid*_ to 0.4, consistent with the approach outlined in the Filipe et al paper [34]. Figure S9 shows probability of symptoms as a function of immune level *M* for a range of *z* values.

To calibrate *κ* and *z* we need data on age-specific incidence in different transmission settings. These data were assembled for our previous calibration (see supplement section 11 in Nguyen et al [12]) and they provide, for each transmission setting, a ratio of incidence in 2-year-olds to incidence in 10-year-olds. These ratios are the orange dots in Figure S10. The model-to-data comparisons in this figure show that *z*-values in the range 4.6 to 6.4, and *κ*-values between 0.1 and 0.4 give the best fits of model to data. The fine-scale heat map in Figure S11 shows that a basic mean-squared error optimization gives z = 5.4 and *κ* = 0.3 as the best-fit values. These values are used in the simulation.

**Figure S8:**
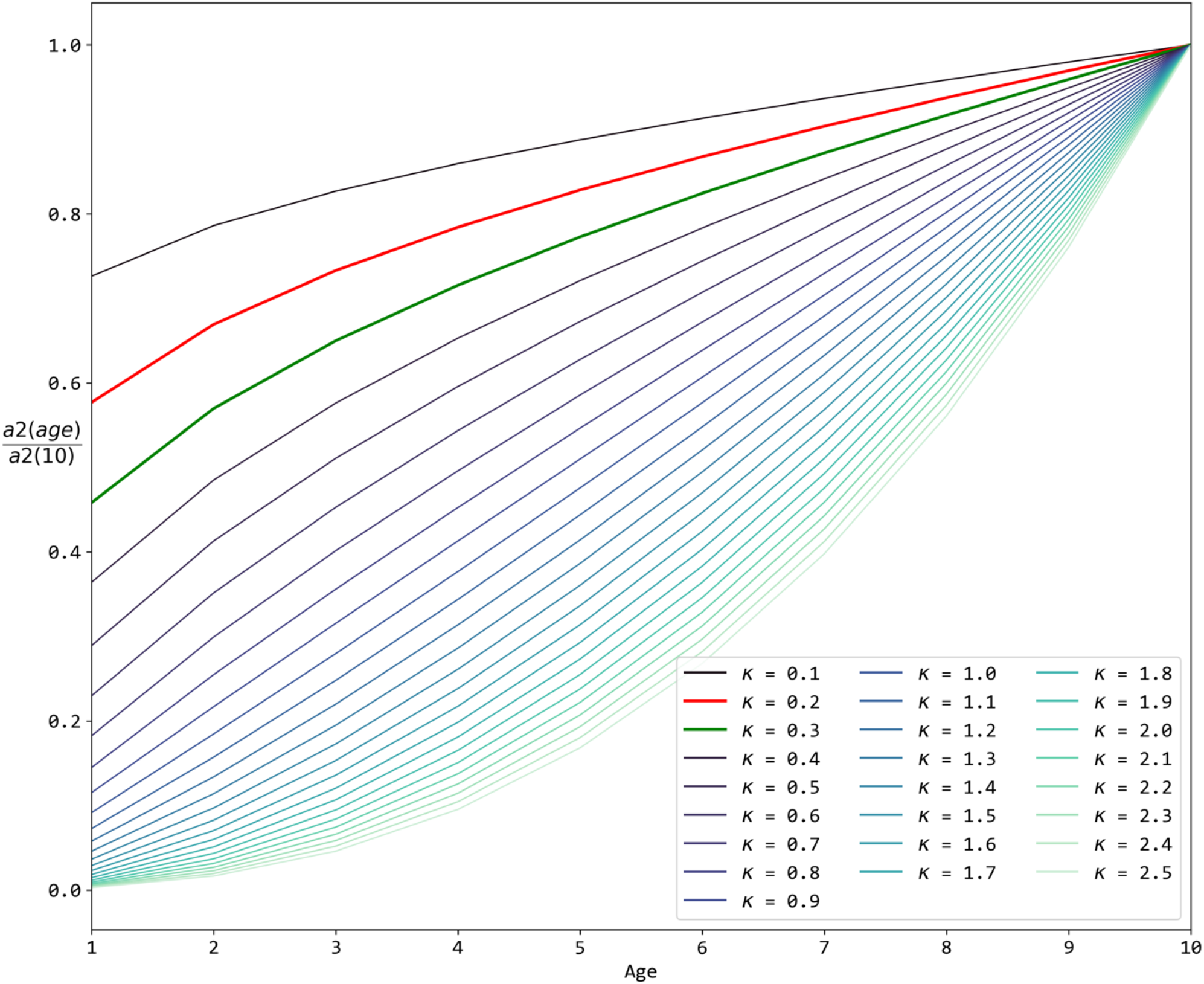
The *y*-axis shows a child’s relative immune acquisition rate when compared to a ten-year old, based on Nguyen et al [12]. Different lines are drawn for different values of the *κ* parameter ranging from 0.1 to 2.5. The red (*κ*=0.2, for the no treatment scenarios in Figure S11) and green (*κ*=0.3, for the 50% treatment coverage scenarios in the remaining figures) lines denote the selected *κ*-values after calibration. Higher *κ* means that children in lower age groups will have slow immune acquisition rates until they reach the normal rate calibrated for ten-year-olds.

**Figure S9:**
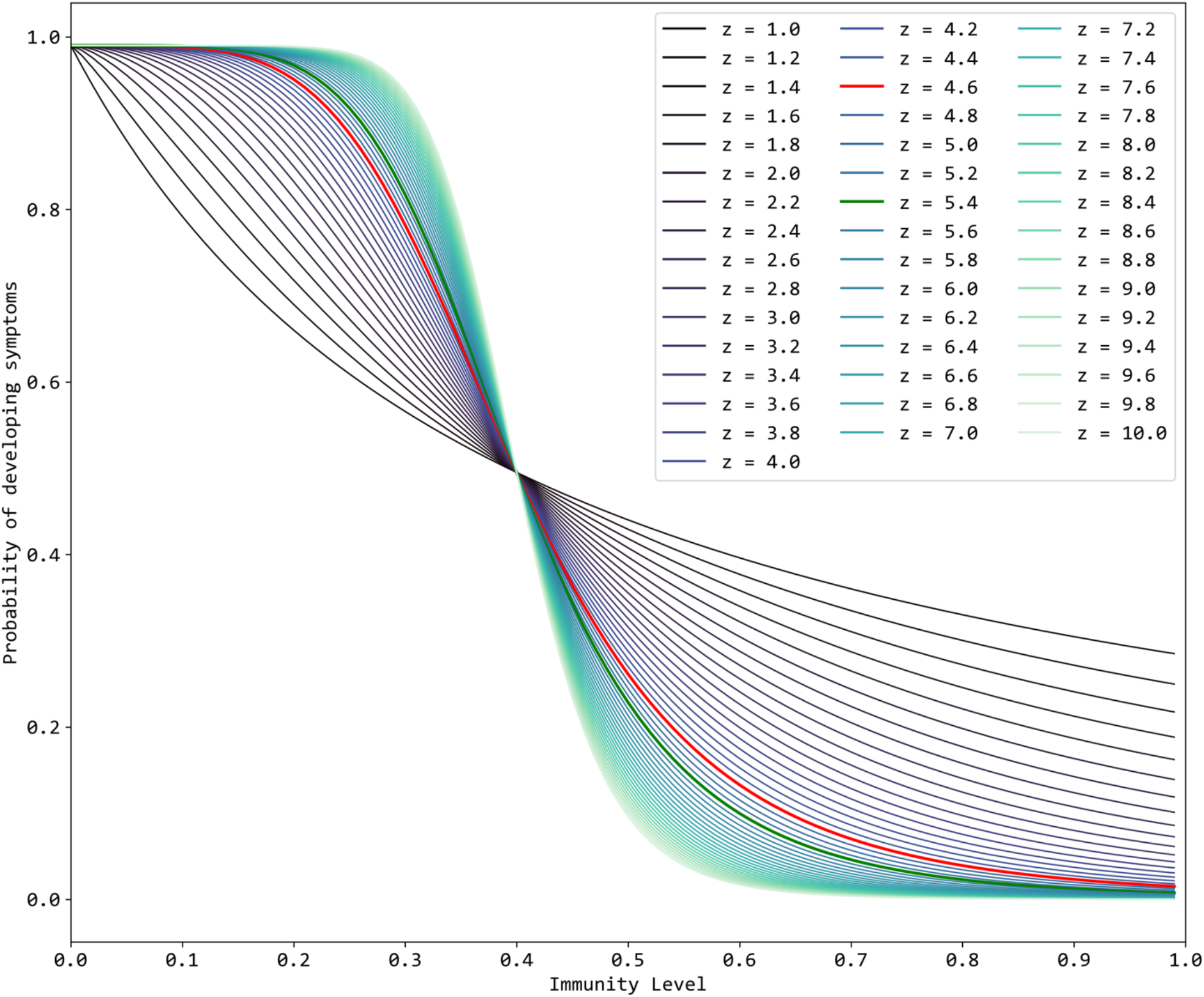
The probability of progressing to clinical disease after an infectious bite (i.e. a symptomatic case) based on the host’s immune level and the parameter *z*, according to Nguyen et al [12]. The red (*z* = 4.6, for the no treatment scenarios in Figure S11) and green (*z* = 5.4, for the 50% treatment coverage scenarios in the remaining figures) lines denote the selected z after calibration.

**Figure S10:**
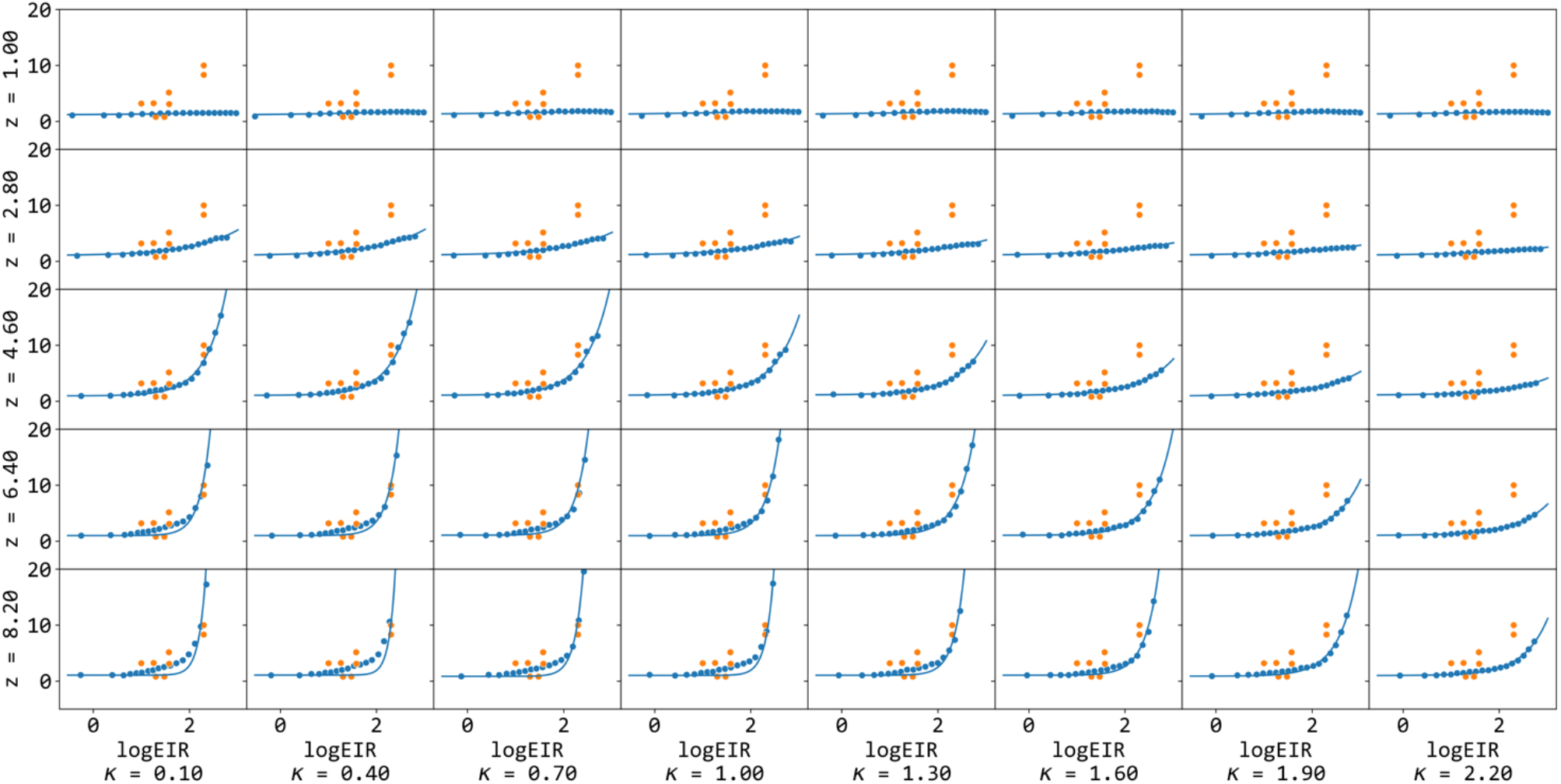
Fitting of *κ* and *z* values. The *y*-axis shows the ratio of the number of annual clinical episodes in 2-year-olds to the number in 10-year-olds plotted against log_10_-EIR. Ten simulations are run for each combination of *κ* and *z* (dots in blue) and those combinations are shown next to the data from Table S7 in Nguyen et al [12] (dots in orange). Treatment coverage here is 50% with an 80% efficacy, 7-day half-life drug.

**Figure S11:**
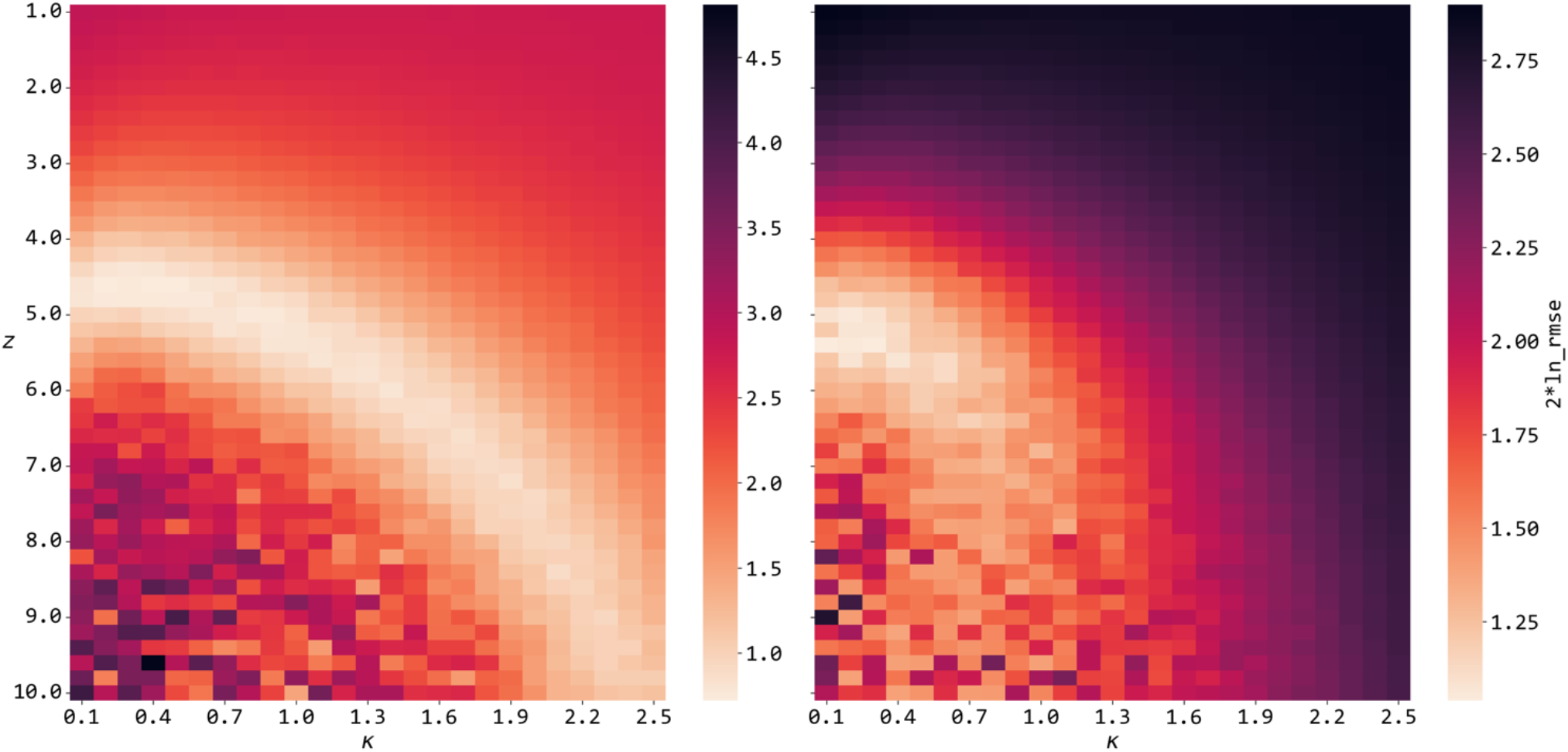
Heatmap shows the log sum of squared errors between the blue and orange dots in Figure S10. The left heatmap presents simulations with no treatment and the optimal fit *z* and *κ* (the brightest spot in the heatmap) were 4.6 and 0.2 respectively. When treatment coverage is 50% (80% efficacy drug with a 7-day half-life) the optimal values are *κ* = 0.3 and *z* = 5.4, corresponding to the brightest spot in the right heatmap.

**Figure S12:**
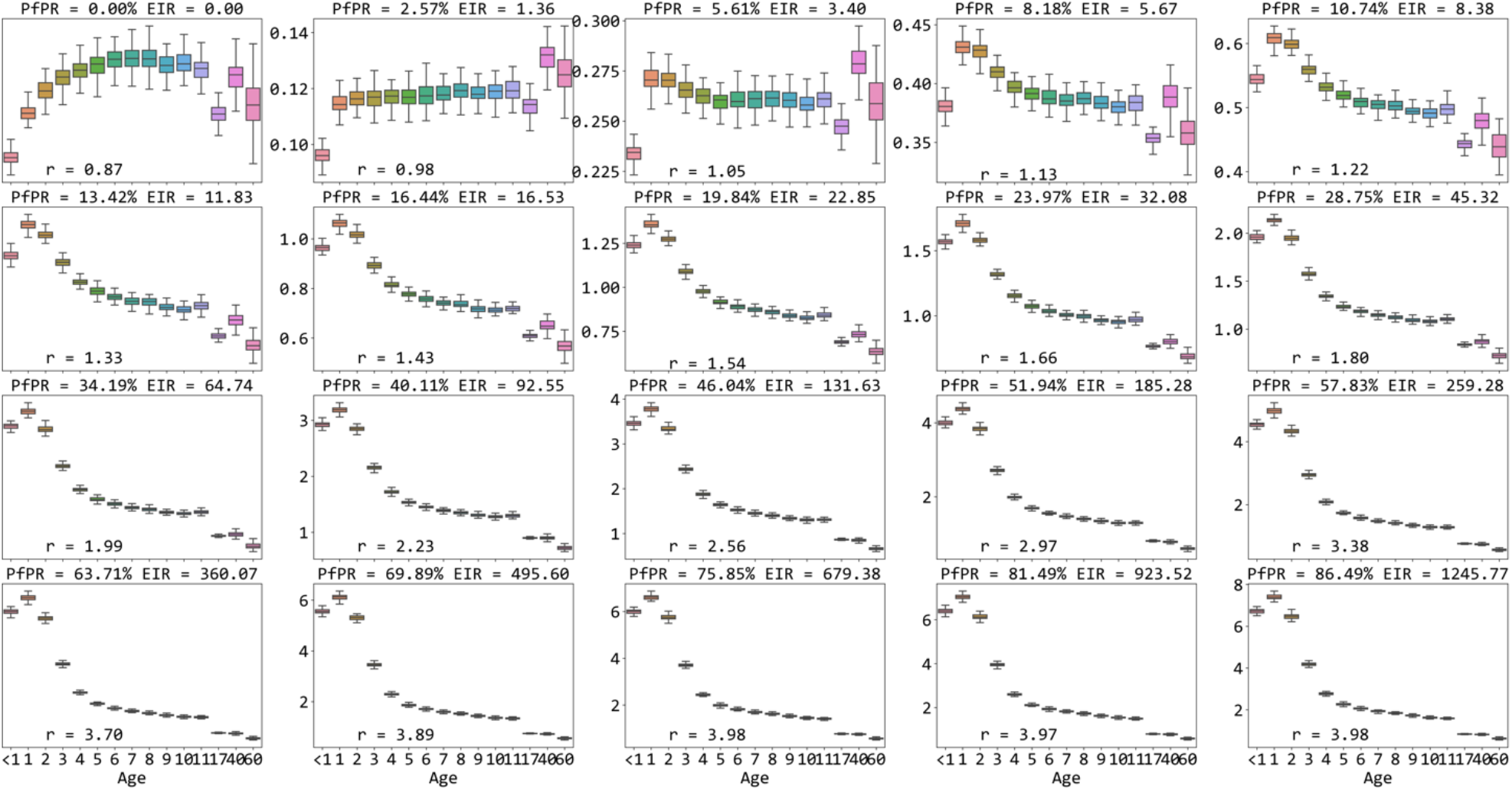
The age-specific clinical episodes per year, under different transmission intensities (EIR increases from left to right and from top to bottom). The *r*-values in the bottom left corner show the ratio of clinical episodes in two-year olds to clinical episodes in ten-year olds in the simulation. Boxplots show medians and interquartile ranges from 100 simulations. All simulations run with *κ* = 0.3 and *z* = 5.4 with 50% treatment coverage with a 7-day half-life drug of approximately 80% efficacy.

**Figure S13:**
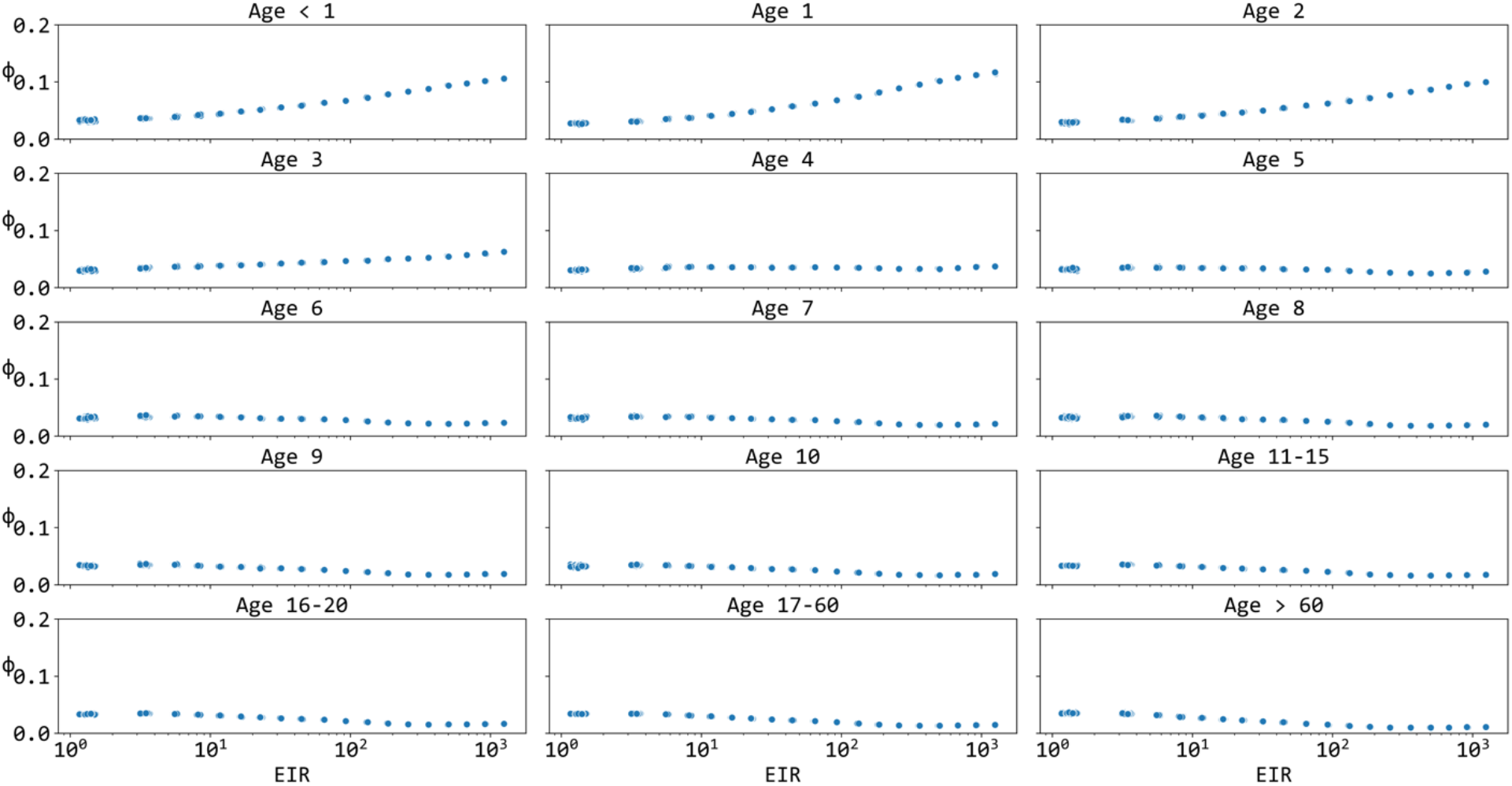
100 simulations were run to equilibrium at different EIR levels ranging from 1.0 to >1000. Treatment coverage is 50%, with a 7-day half-life drug of approximately 80% efficacy. Immune acquisition parameters *κ* = 0.3 and *z* = 5.4. The *y*-axis value *φ* shows what fraction of infections are in individuals experiencing symptoms.

**Figure S14:**
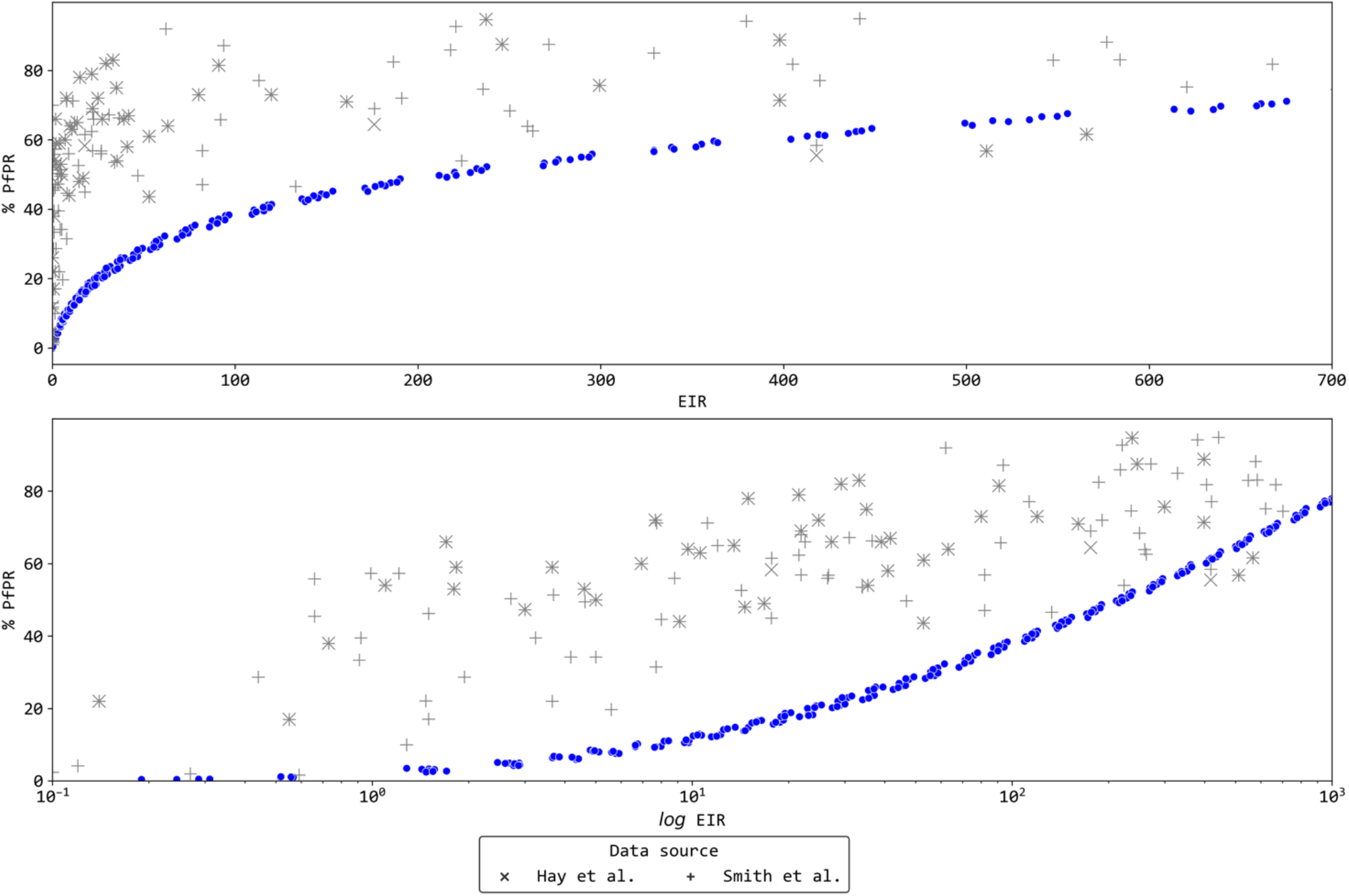
Relationship between EIR and blood slide prevalence shown on linear scale (top) and log scale (bottom). The data points in blue dot are from the model, and parasite positive individuals are counted if their parasitaemia is higher than 10 parasites per microliter. The data points with “+” and “×” symbols are data from Smith et al [35] and Hay et al and Cameron et al [36,37]. Model output shown in blue circles. The model was run with Γ-distributed relative biting rate with a coefficient of variation equal to 2.0. Treatment coverage is 50% with a 7-day half-life drug of approximately 80% efficacy.

**Figure S15:**
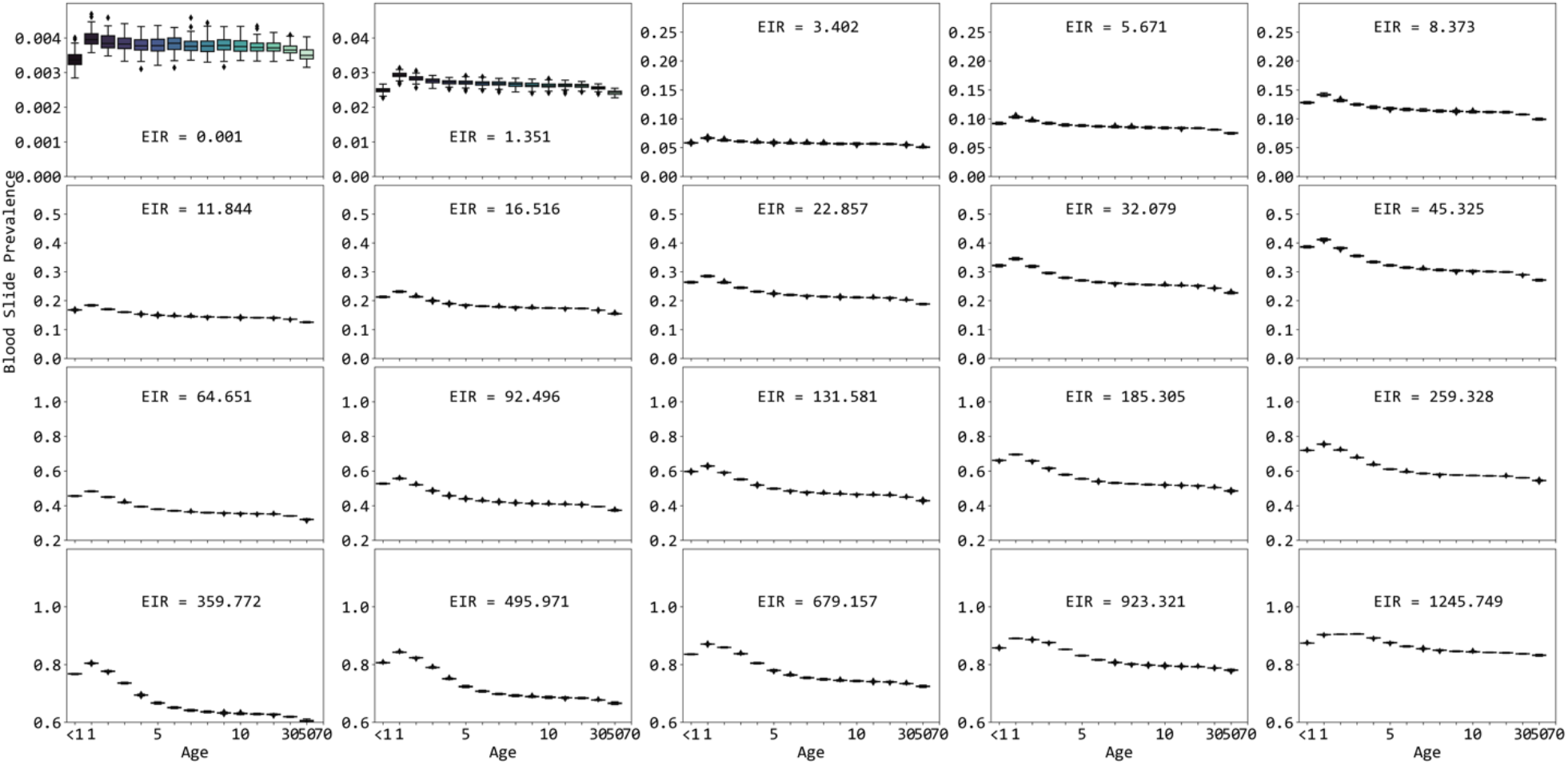
Panels shows age-specific blood-slide prevalence for different transmission intensities; *κ* = 0.3, *z* = 5.4, treated coverage is 50% with a 7-day half-life drug of approximately 80% efficacy.

**Figure S16:**
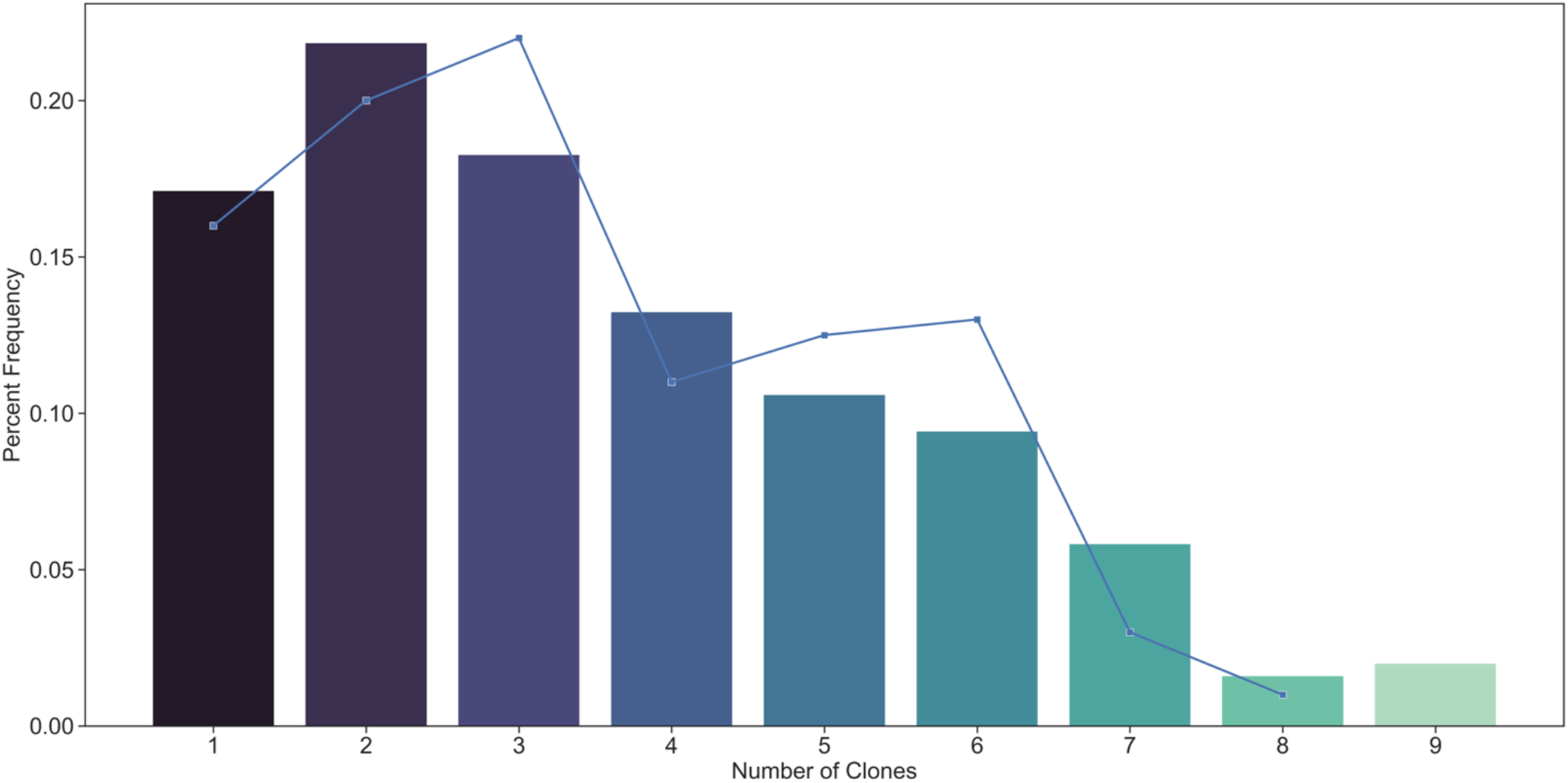
Distribution of number of clones per infection. The bar graph shows the output from our simulation with different EIR (*κ* = 0.3, *z* = 5.4, 50% treatment coverage, with a 7-day half-life drug of approximately 80% efficacy).

**Figure S17:**
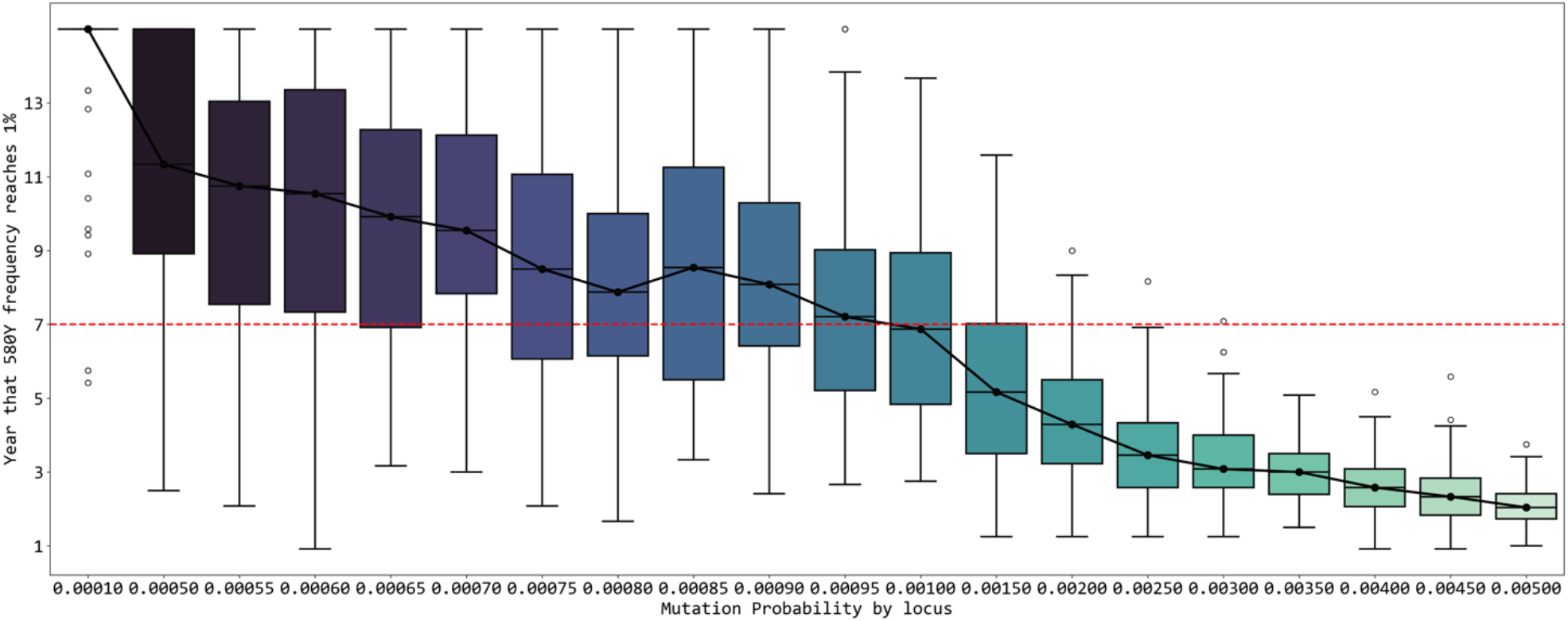
Calibration of mutation probability in the model. Target calibration is to have 580Y frequency reach 0.01 allele frequency after 7.0 years of treatment with DHA-PPQ with 40% treatment coverage, at 25% prevalence and *c*_*R*_ = 0.0005. The *x*-axis shows different values of the model variable mutation_probability_per_locus (mutation and within-host fixation probability if treatment is given that selects for that particular mutation) while the *y*-axis shows the year that 580Y frequency reaches 0.01. The calibration-selected value is mutation_probability_per_locus = 0.001.

---

**Figure S1a:**
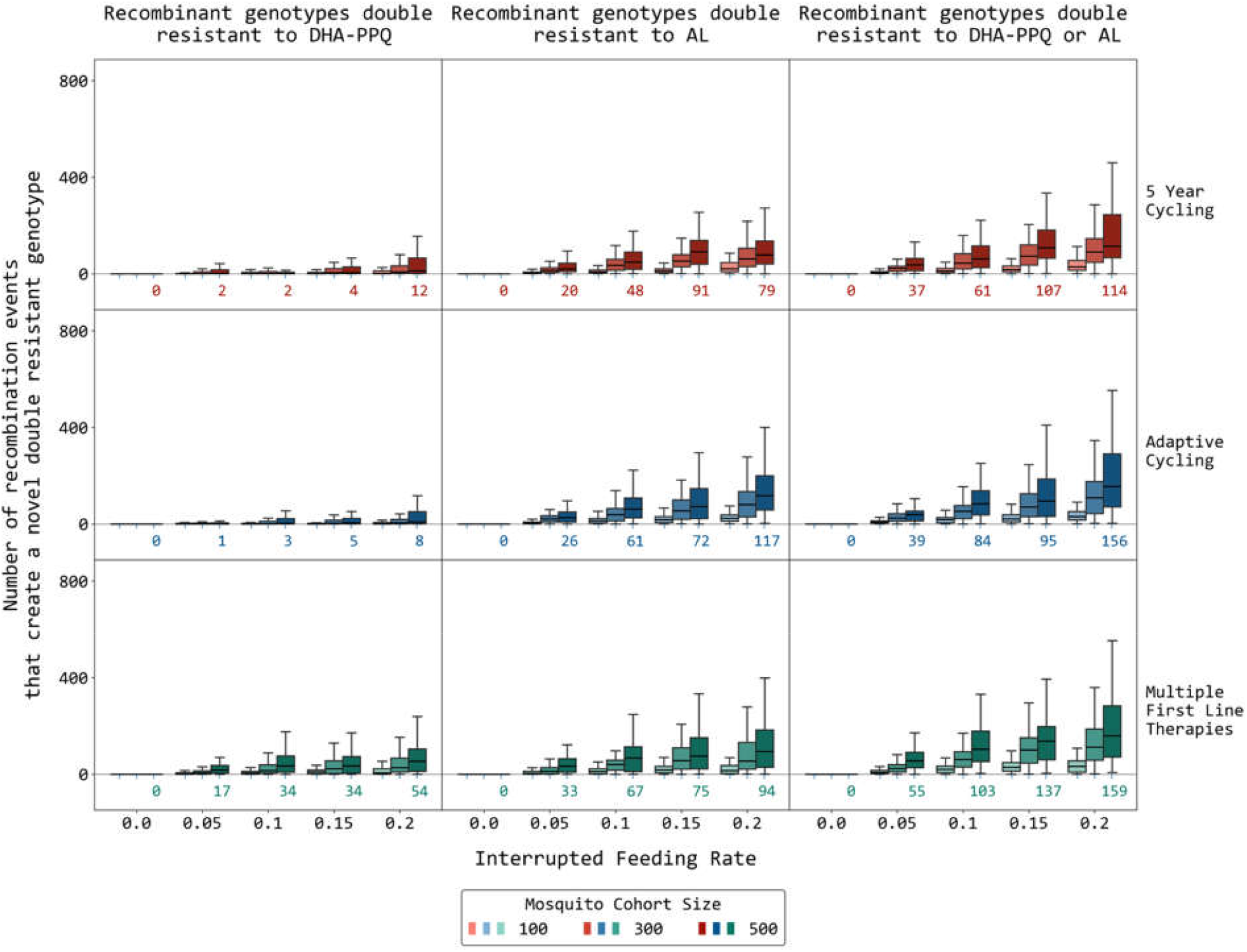
Number of recombination events that produce novel recombinant double-resistant genotypes. This figure has the same settings and format as in Figure 1 in the main text but recombination from multi-clonal within-host parasites infection is not activated, meaning all counted recombination events are from mosquito interrupted feeds.

**Figure S2a:**
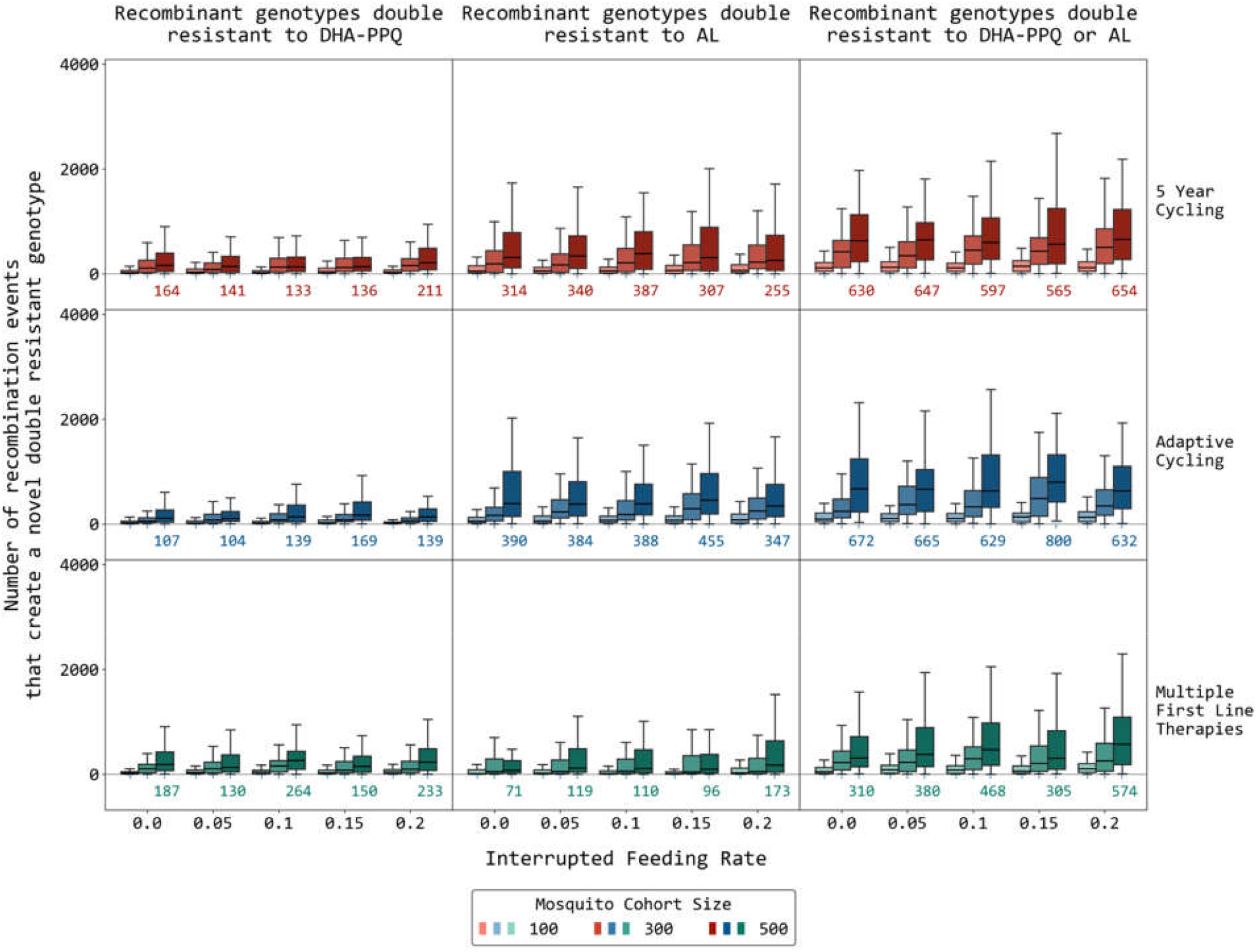
Number of recombination events that produce novel recombinant double-resistant genotypes. This figure has the same settings and format as in Figure 1 in the main text but with *c*_*R*_ = 0.005.

**Figure S3a:**
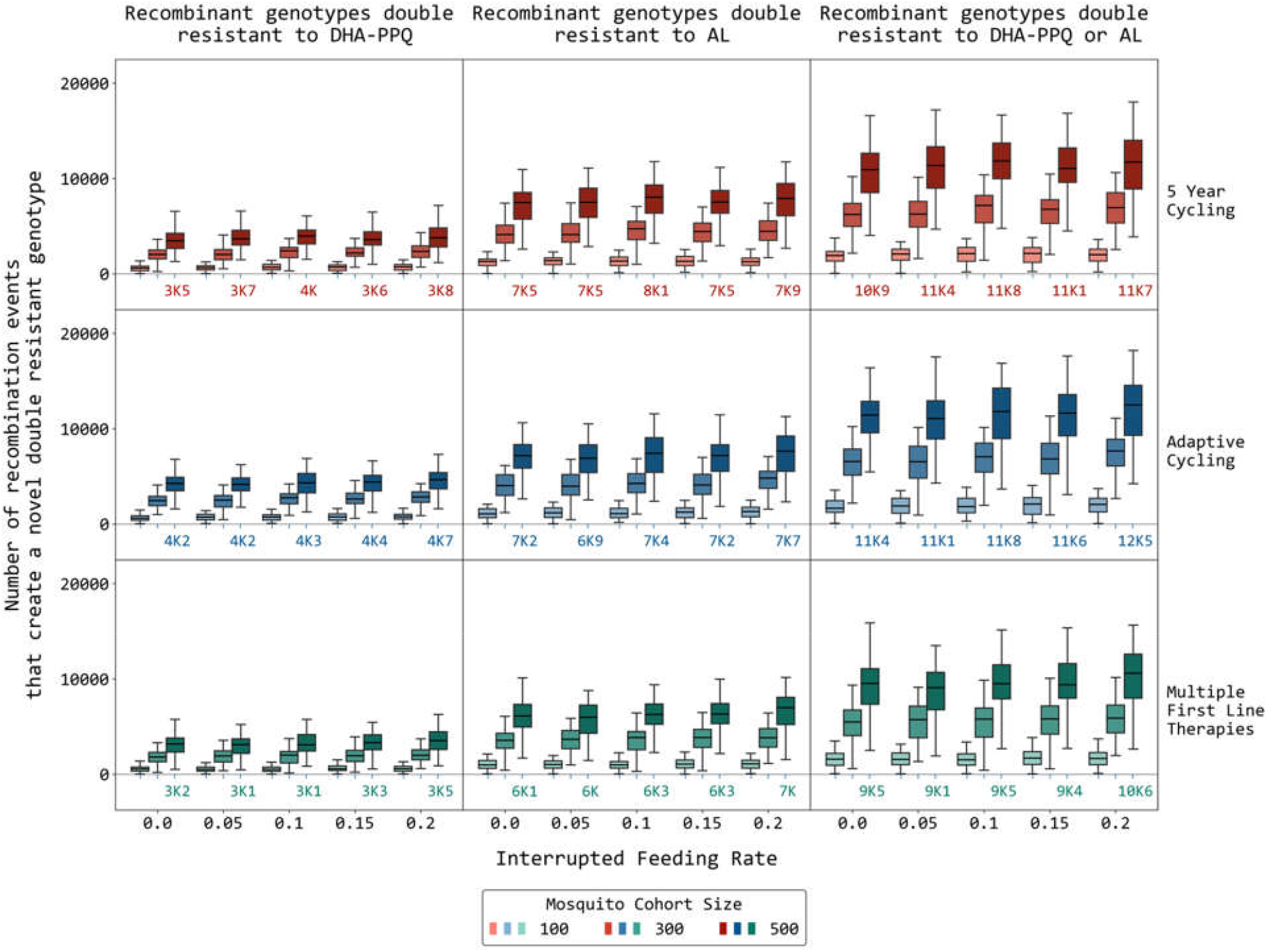
Number of recombination events that produce novel recombinant double-resistant genotypes. This figure has the same settings and format as in Figure 1 in the main text but prevalence is 25% and *c*_*R*_ = 0.0005.

**Figure S4a:**
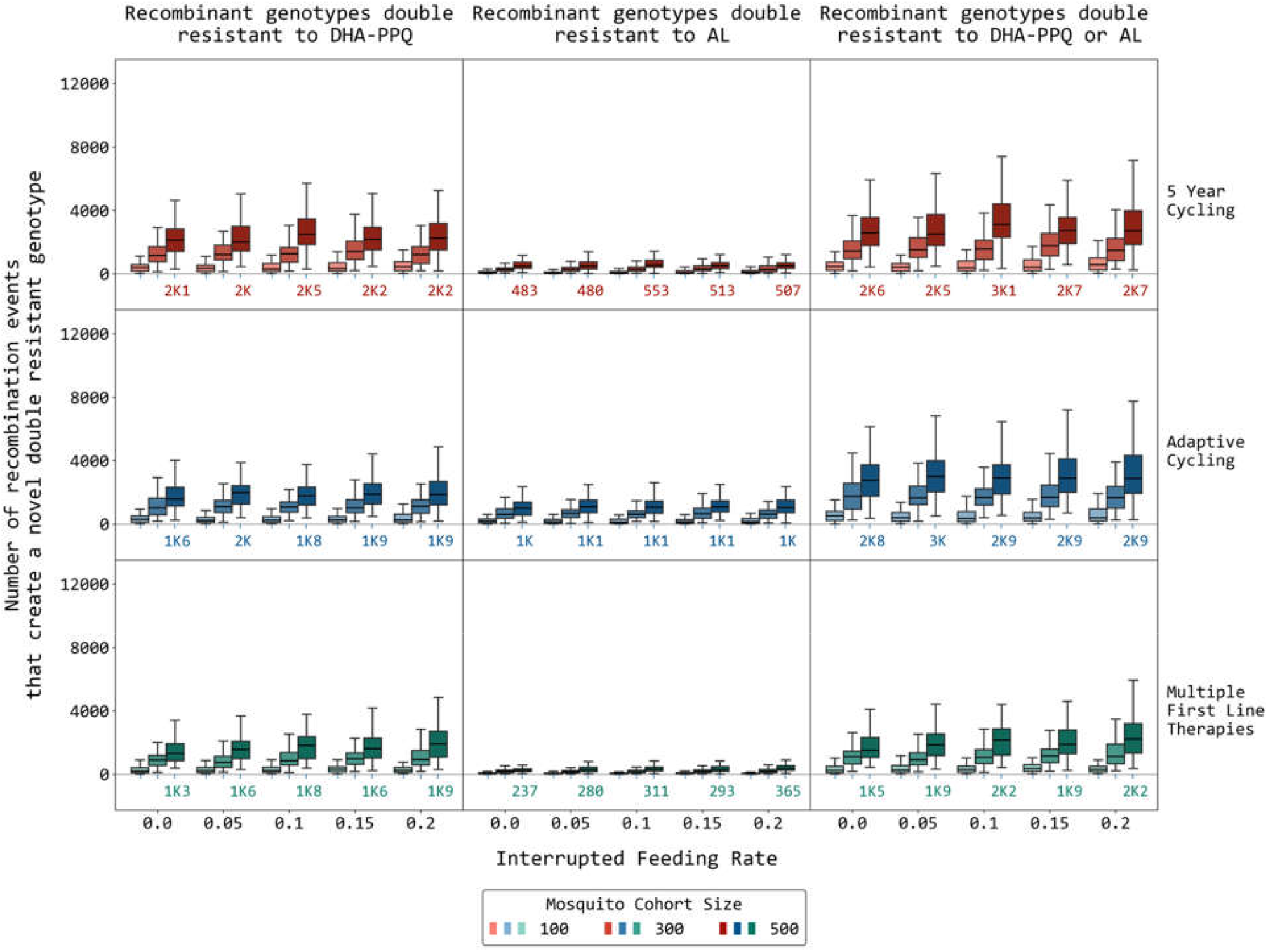
Number of recombination events that produce novel recombinant double-resistant genotypes. This figure has the same settings and format as in Figure 1 in the main text but prevalence is 25% and *c*_*R*_ = 0.005.

**Figure S5a:**
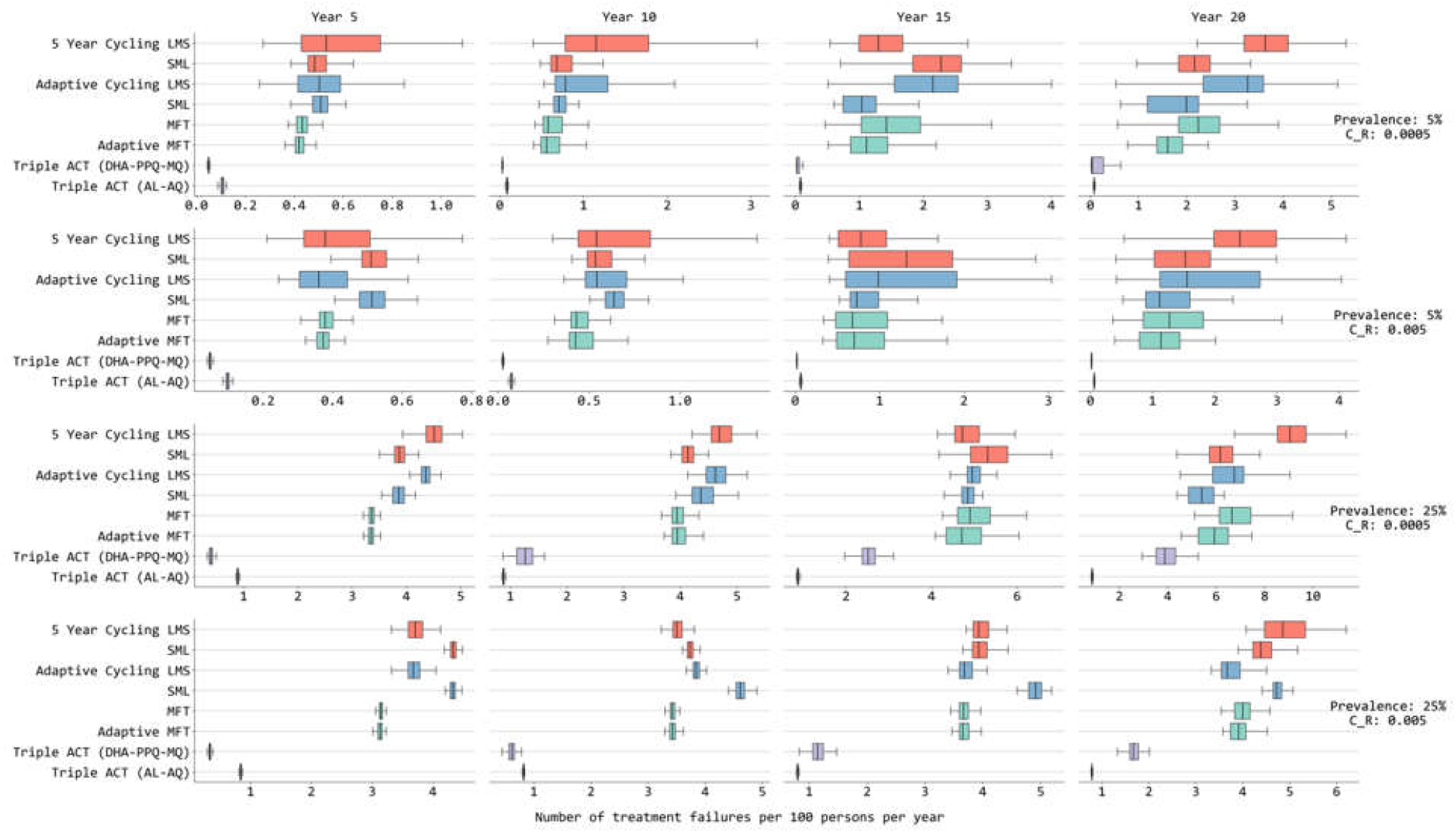
Number of treatment failures per 100 persons per year, counted through to year 5, to year 10, to year 15, and to year 20. This figure has the same settings and format as in Figure 3 in the main text but shows the NTF results using different endpoints.

**Figure S6a:**
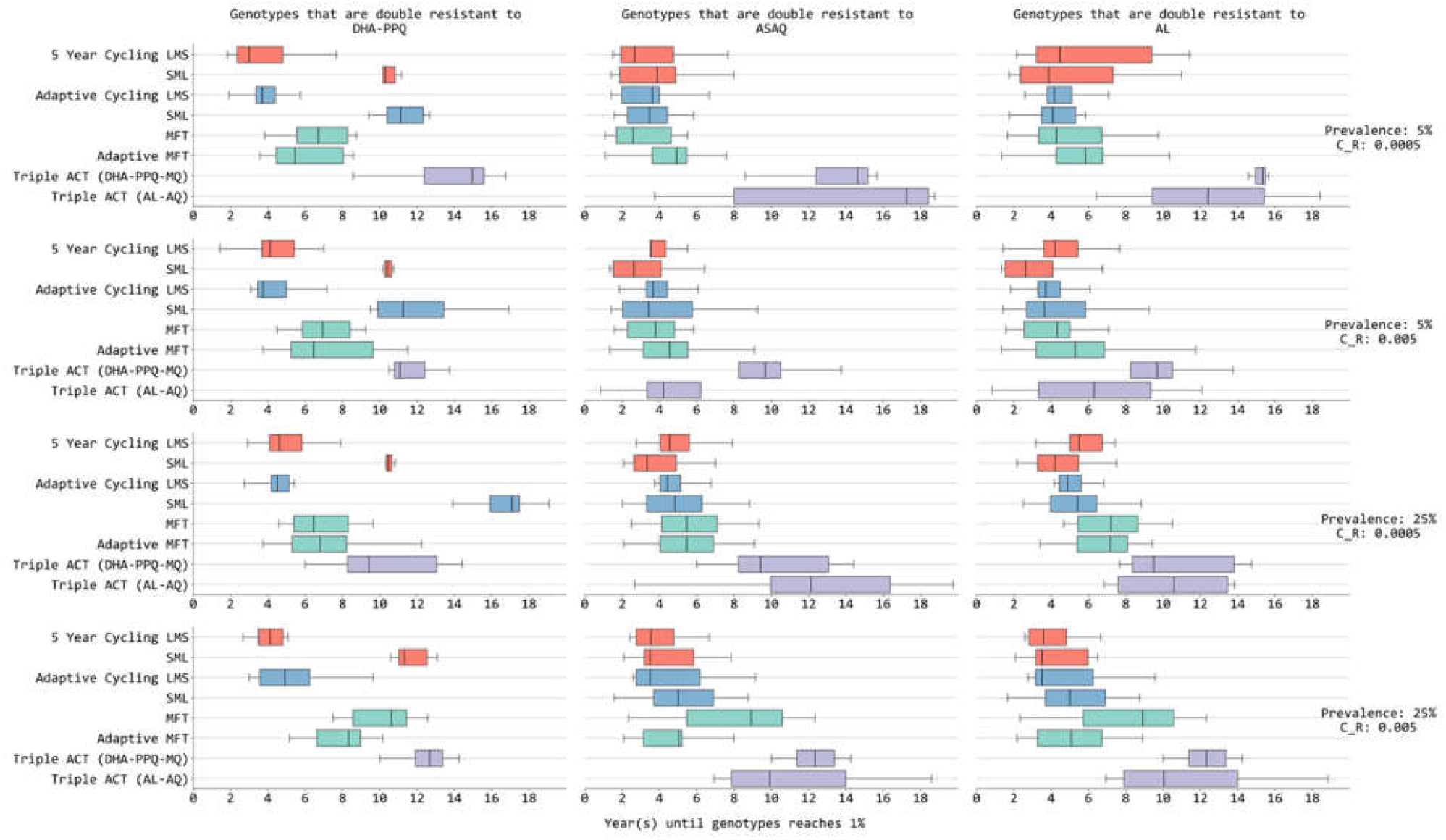
Years (*x*-axis) until double-resistant genotype frequency reaches 0.01. Four rows show different malaria prevalence and cost-of-resistance scenarios. Different colors are for different treatment policies: 5-year cycling (light red), adaptive cycling (light blue), multiple first-line therapies (sea green) and triple ACT (violet). LMS indicates that the order of drugs used in cycling strategies is based on the half-life of the partner drug (**<u>L</u>**ong: DHA-PPQ, **<u>M</u>**edium: ASAQ, **<u>S</u>**hort: AL). When no boxplot is shown, this indicates that the genotype frequency is below 0.01 for the entire 20-year period of simulation.

**Figure S7a:**
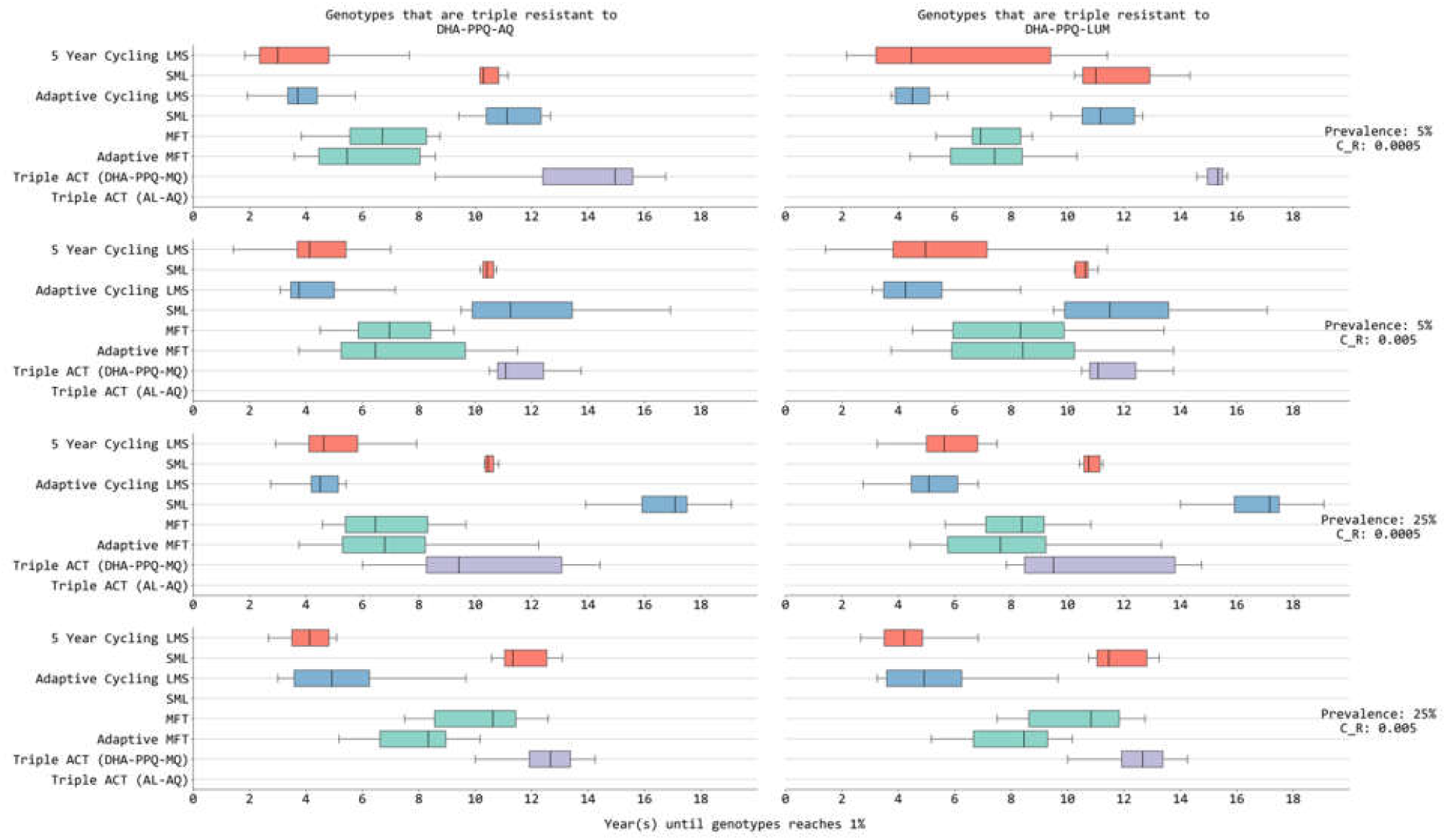
Years until triple-resistant genotype frequency reaches 0.01. A single resistance mutation is sufficient to label the parasites as resistant to a particular drug. The four rows show different malaria prevalence and cost of resistance levels. Columns show the two triple therapies (DHA-PPQ-AQ and DHA-PPQ-LUM) to which triple-resistance could emerge. In each panel, the *x*-axis shows the number of years that it takes for a genotype frequency to reach 0.01 under different treatment strategies: 5-year cycling (light red), adaptive cycling (light blue), multiple first-line therapies (sea green) and triple ACT (violet). LMS indicates the order of therapies used in a cycling strategy based on the half-life of partner drugs (Long: DHA-PPQ, Medium: ASAQ, Short: AL). When no boxplot is shown, this indicates that the genotype frequency is below 0.01 for the entire 20-year period of simulation.

**Figure S8a:**
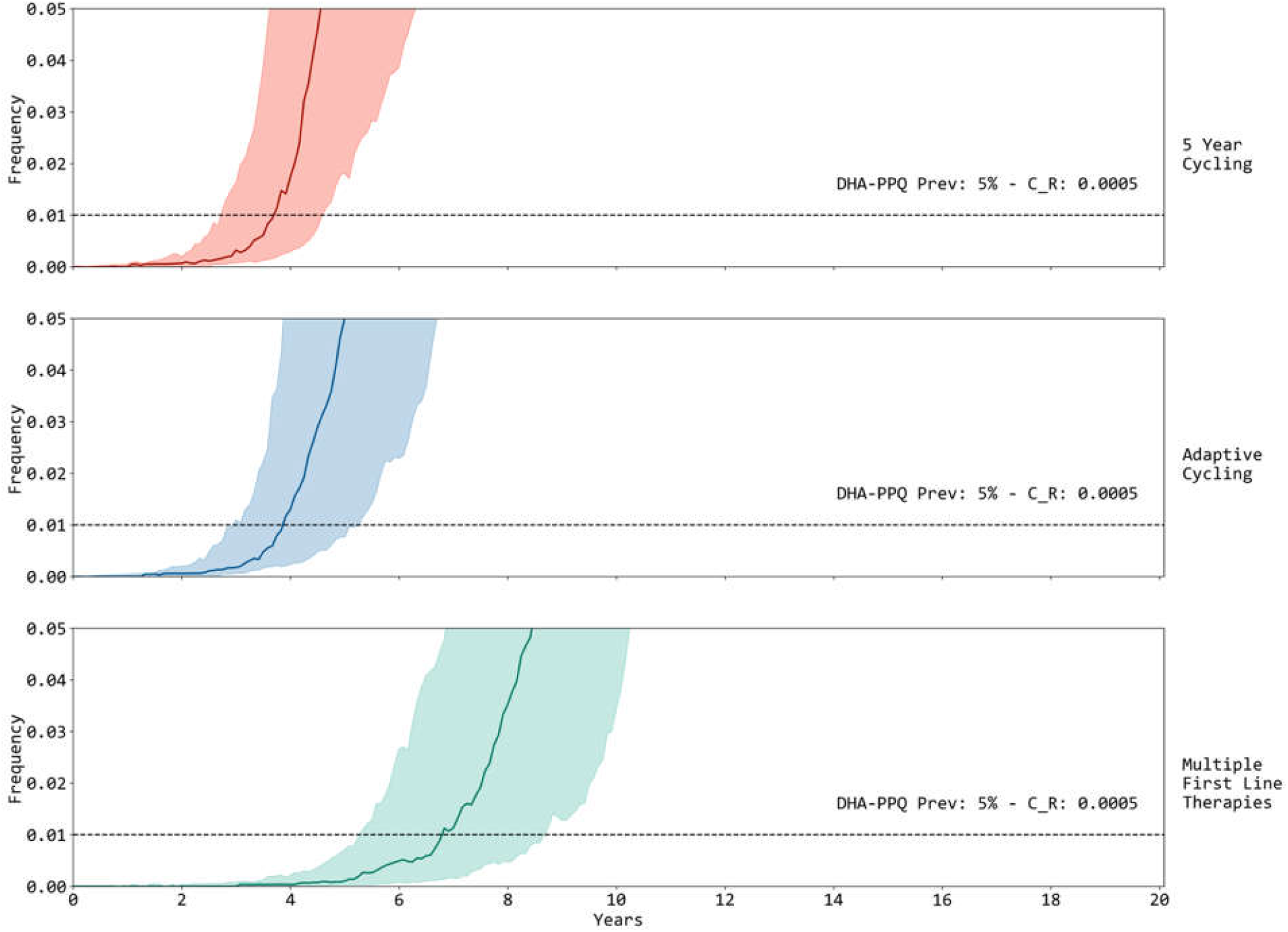
Genotype frequency of double-resistants to DHA-PPQ under three different strategies. Year zero indicates the year that the strategy is deployed, and the frequency range is limited to 0.00 to 0.05 to focus on the early period of emergence. Solid line and shaded regions correspond to median and IQR, respectively. Cycling is LMS.

**Figure S9a:**
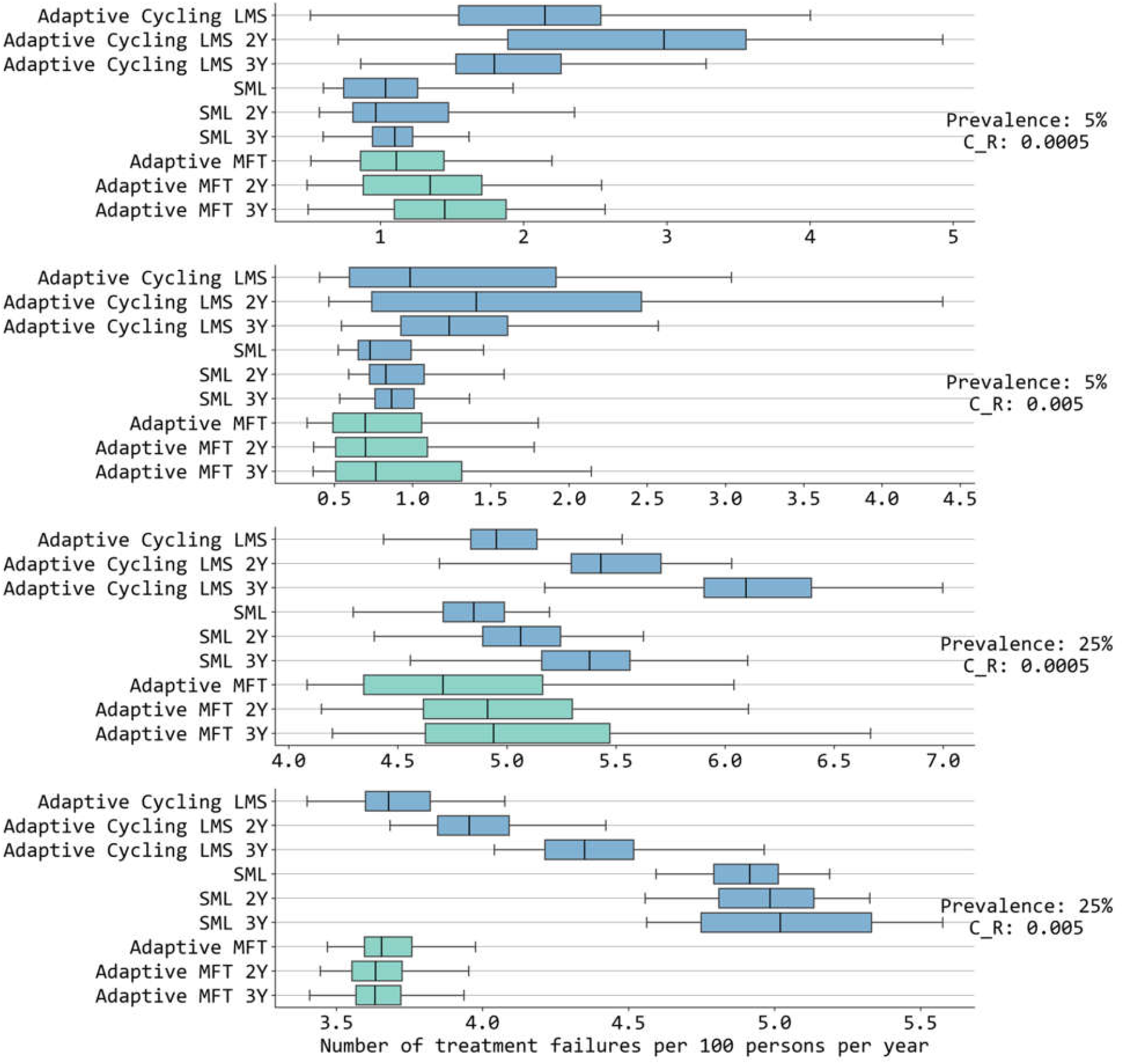
Number of treatment failures under adaptive cycling and adaptive MFT policies with different delay times to a policy switch. In our model, adaptive policies include a one-year delay between the trigger (e.g. resistance levels are too high) and the implementation of a new approach. In this panel, “2Y” and “3Y” show the results of two-year and three-year delays. The *x*-axis shows the number of treatment failures per 100 people per year. LMS indicates the order of drugs used in cycling strategies based on efficient period of partner drugs (**<u>L</u>**ong: DHA-PPQ, **<u>M</u>**edium: ASAQ, **<u>S</u>**hort: AL).

**Figure S10a:**
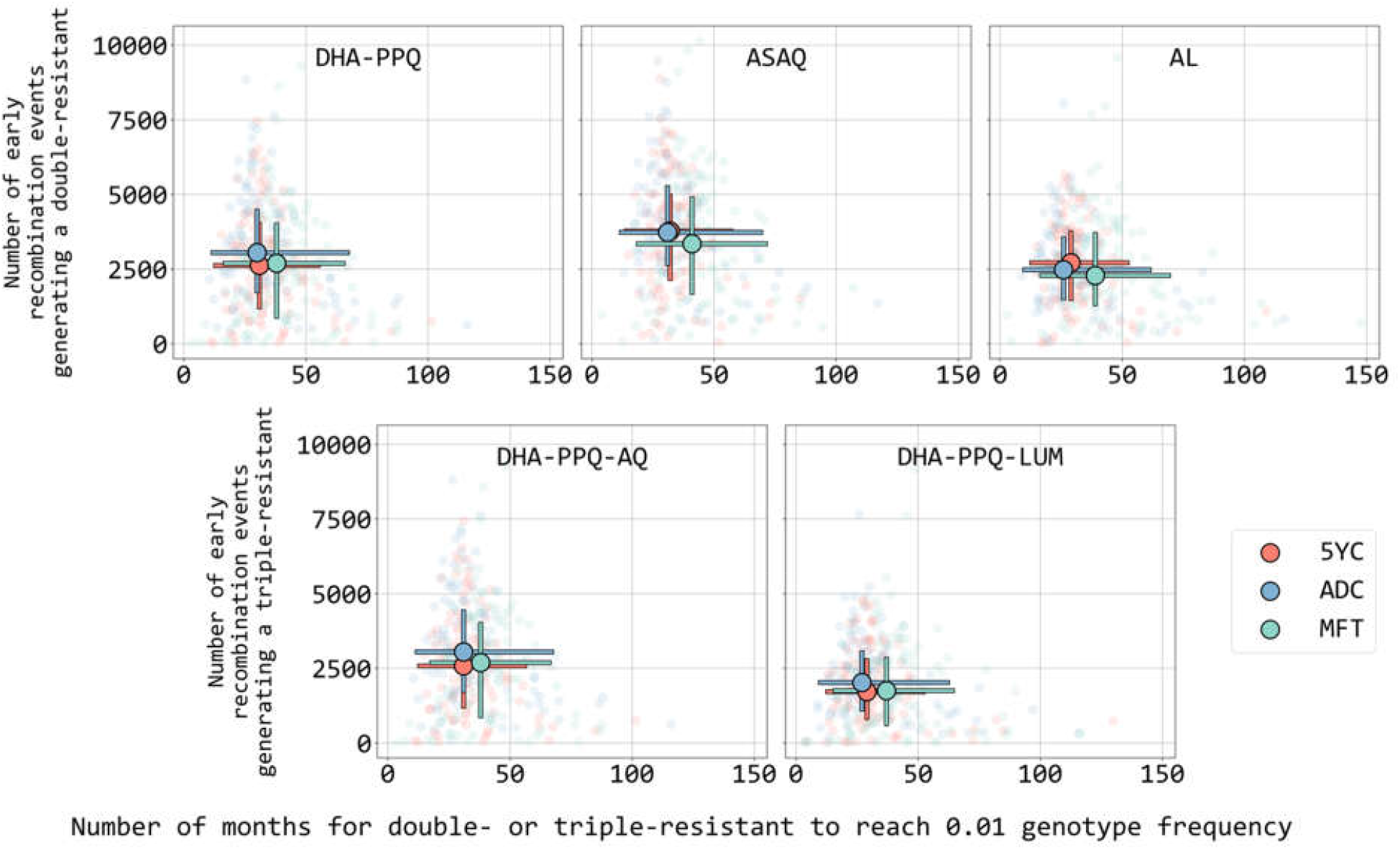
Number of *early recombination events* (*y*-axis) creating a double-resistant or triple-resistant genotype (see text inside each panel). An *early recombination event* is one that occurs prior to the time that the frequency of this MDR genotype reaches 0.01. The format of this figure is the same as Figure 5 in main text but *c*_*R*_ = 0.005. Circles show the median values for three strategies: five-year LMS cycling in red (5YC), adaptive LMS cycling in blue (ADC), and multiple first-line therapies in green (MFT). Bars show interquartile ranges, and the full set of outcomes is shown as a light scatterplot in the background. The *x*-axes count number of months *after first appearance*.

**Figure S11a:**
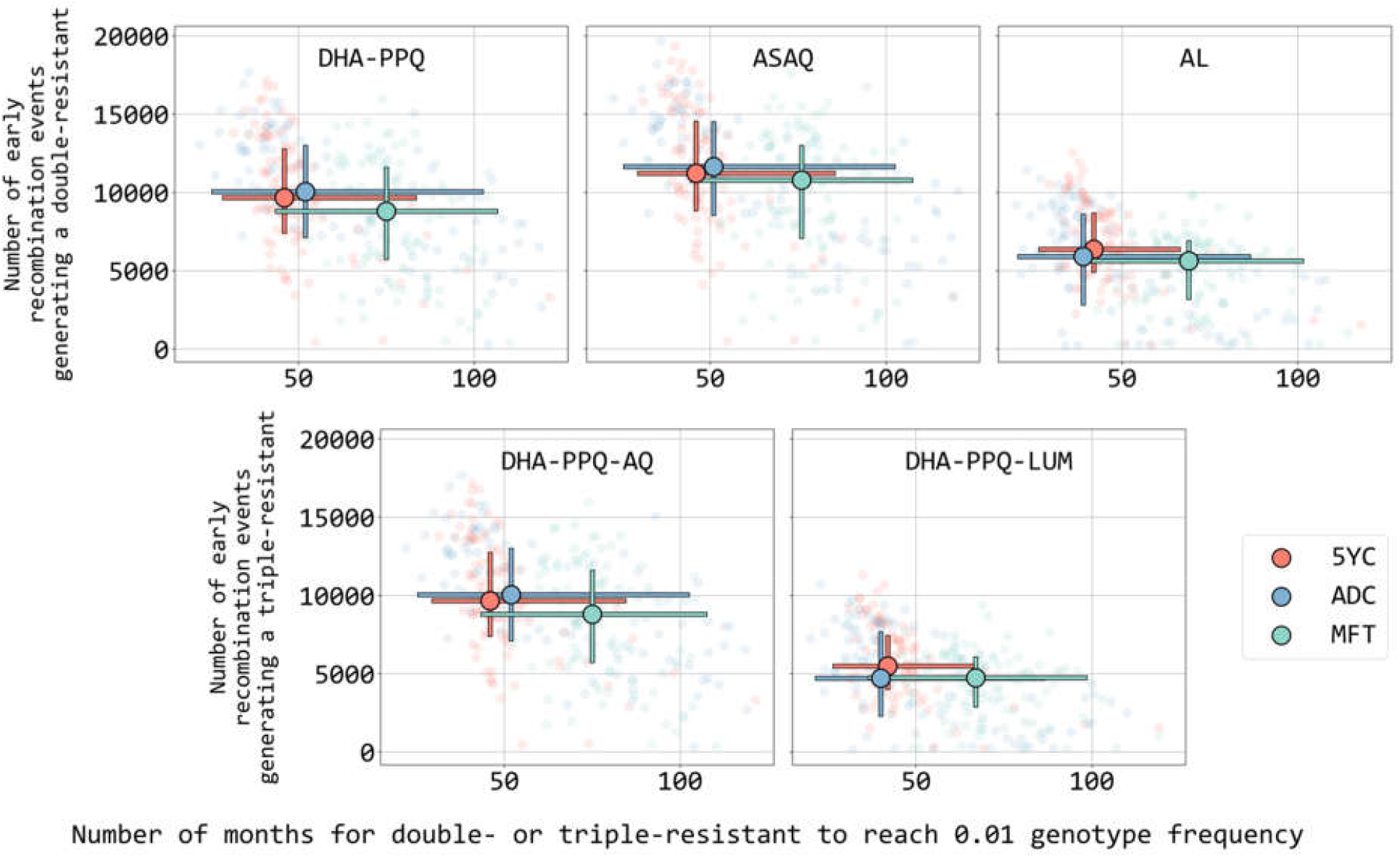
Number of *early recombination events* (*y*-axis) creating a double-resistant or triple-resistant genotype (see text inside each panel). An *early recombination event* is one that occurs prior to the time that the frequency of this MDR genotype reaches 0.01. The format of this figure is the same as Figure 5 in main text but prevalence is 25% and *c*_*R*_ = 0.0005. Circles show the median values for three strategies: five-year LMS cycling in red (5YC), adaptive LMS cycling in blue (ADC), and multiple first-line therapies in green (MFT). Bars show interquartile ranges, and the full set of outcomes is shown as a light scatterplot in the background. The *x*-axes count number of months *after first appearance*.

**Figure S12a:**
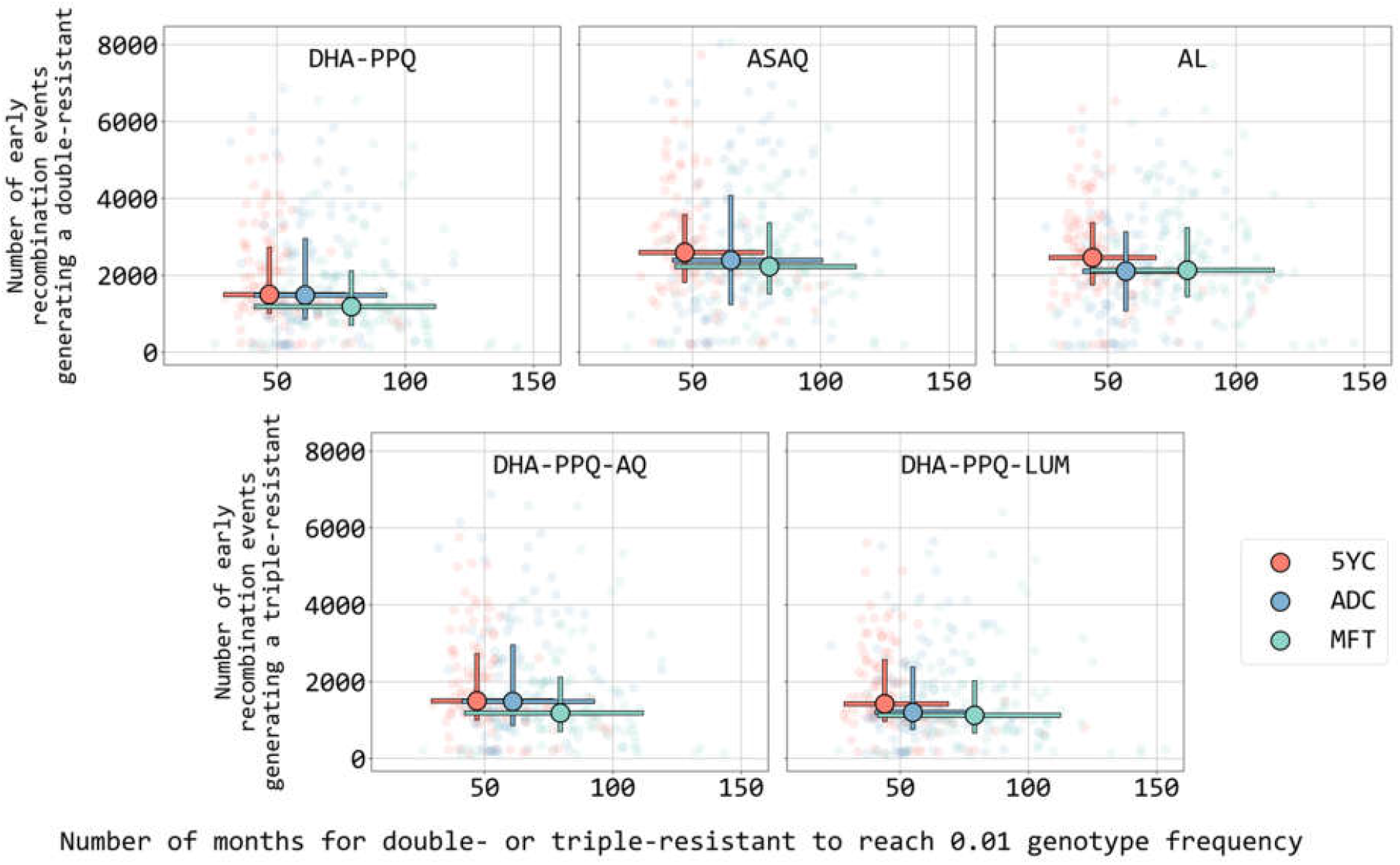
Number of *early recombination events* (*y*-axis) creating a double-resistant or triple-resistant genotype (see text inside each panel). An *early recombination event* is one that occurs prior to the time that the frequency of this MDR genotype reaches 0.01. The format of this figure is the same as Figure 5 in main text but prevalence is 25% and *c*_*R*_ = 0.005. Circles show the median values for three strategies: five-year LMS cycling in red (5YC), adaptive LMS cycling in blue (ADC), and multiple first-line therapies in green (MFT). Bars show interquartile ranges, and the full set of outcomes is shown as a light scatterplot in the background. The *x*-axes count number of months *after first appearance*.

**Figure S13a:**
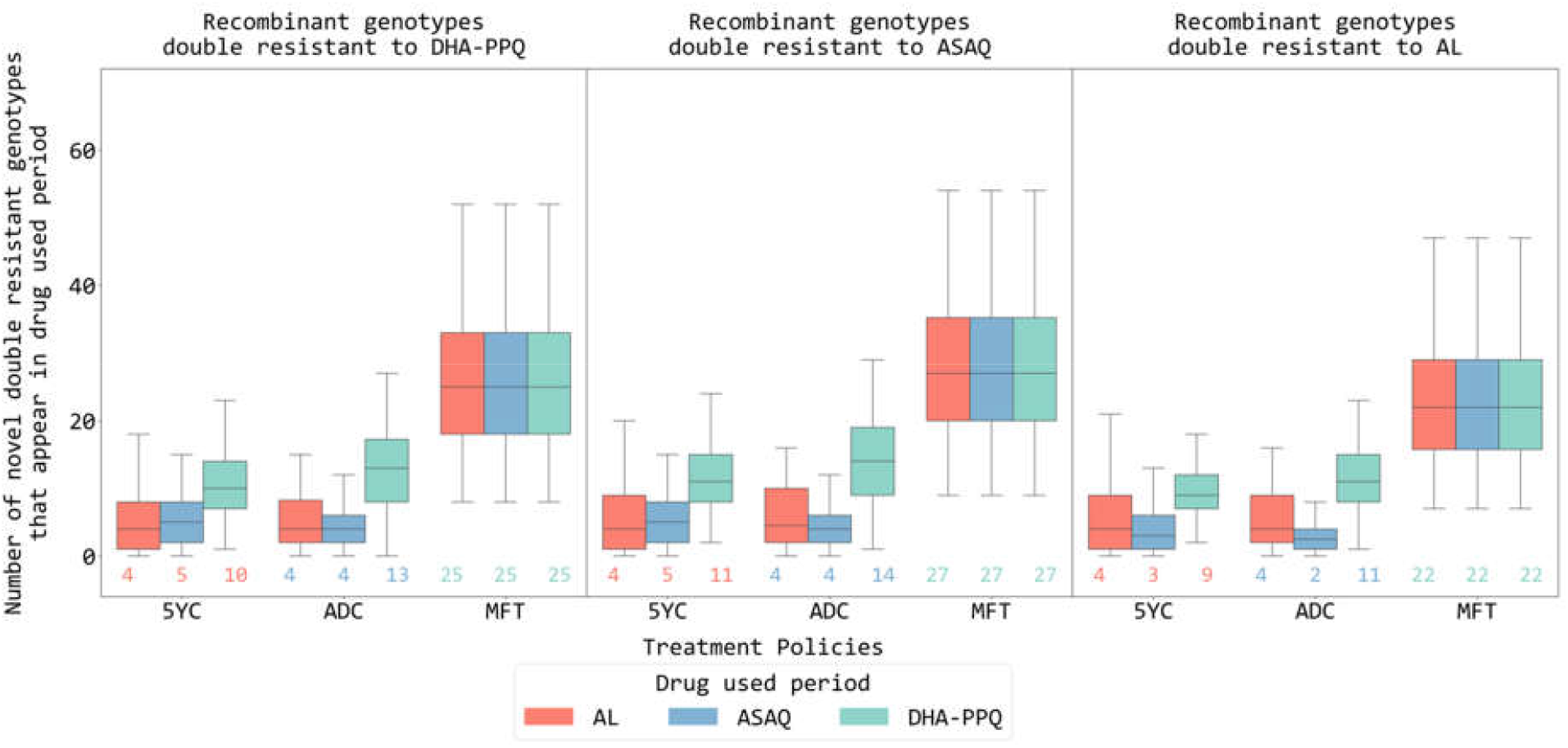
The *y*-axis shows the number of different double-resistant genotypes produced by recombination under each strategy. The boxplot colors show under which ‘drug use period’ this new genetic diversity was generated. For MFT the entire 15-year period is considered a ‘drug use period’ for each of the three ACTs. The visual comparisons to make are red box plots to red box plots, blue box plots to blue box plots, etc. Cycling strategies are LMS. In this figure the prevalence is 5% with daily cost of resistance is 0.0005 and mosquito cohort size is 100 with 20% interrupted feeding rate. <u>Under MFT, recombination generates a greater diversity of genotypes</u> (when compared to cycling strategies) but (1) there are fewer of these recombination events under MFT, and (2) these different recombinant genotypes have a slower evolutionary path to 0.01 genotype frequency (see Figures S10-S12).

**Figure S14a:**
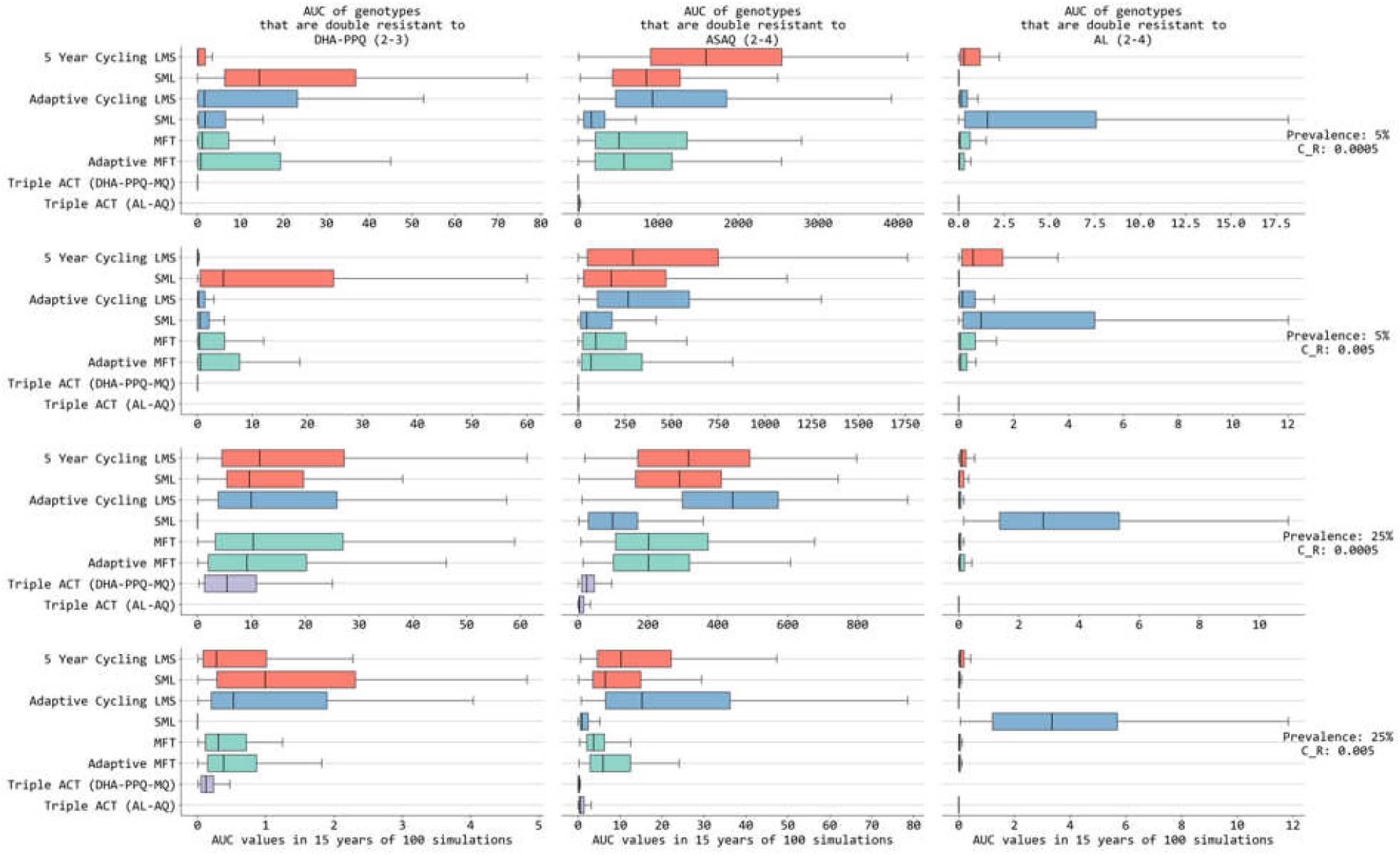
AUC (total area under the frequency curve of a maximally-resistant genotype) of genotypes that are double resistant to DHA-PPQ, ASAQ, and AL (in the three columns). This is the same AUC measure of multi-drug resistance risk as presented in Figures 4 and 5 or Li et al (*Nat Commun*, 15:1390, 2024). The codes “2-3” and “2-4” at top indicate that the genotypes are double-resistant (first digit) to a specific drug with 3 or 4 mutations (second digit). The four rows show different malaria scenarios with different prevalence and cost of resistance. In each panel, the *x*-axis shows the AUC values of double-resistant genotypes under different treatment strategies. Different colors are for different treatment policies: 5-year cycling (light red), adaptive cycling (light blue), multiple first-line therapies (sea green) and triple ACT (violet). LMS indicate the order of drugs used in cycling strategies based on half-life of partner drugs (Long: DHA-PPQ, Medium: ASAQ, Short: AL). When no boxplot is shown this means that no double-resistant genotypes were generated.

**Figure S15a:**
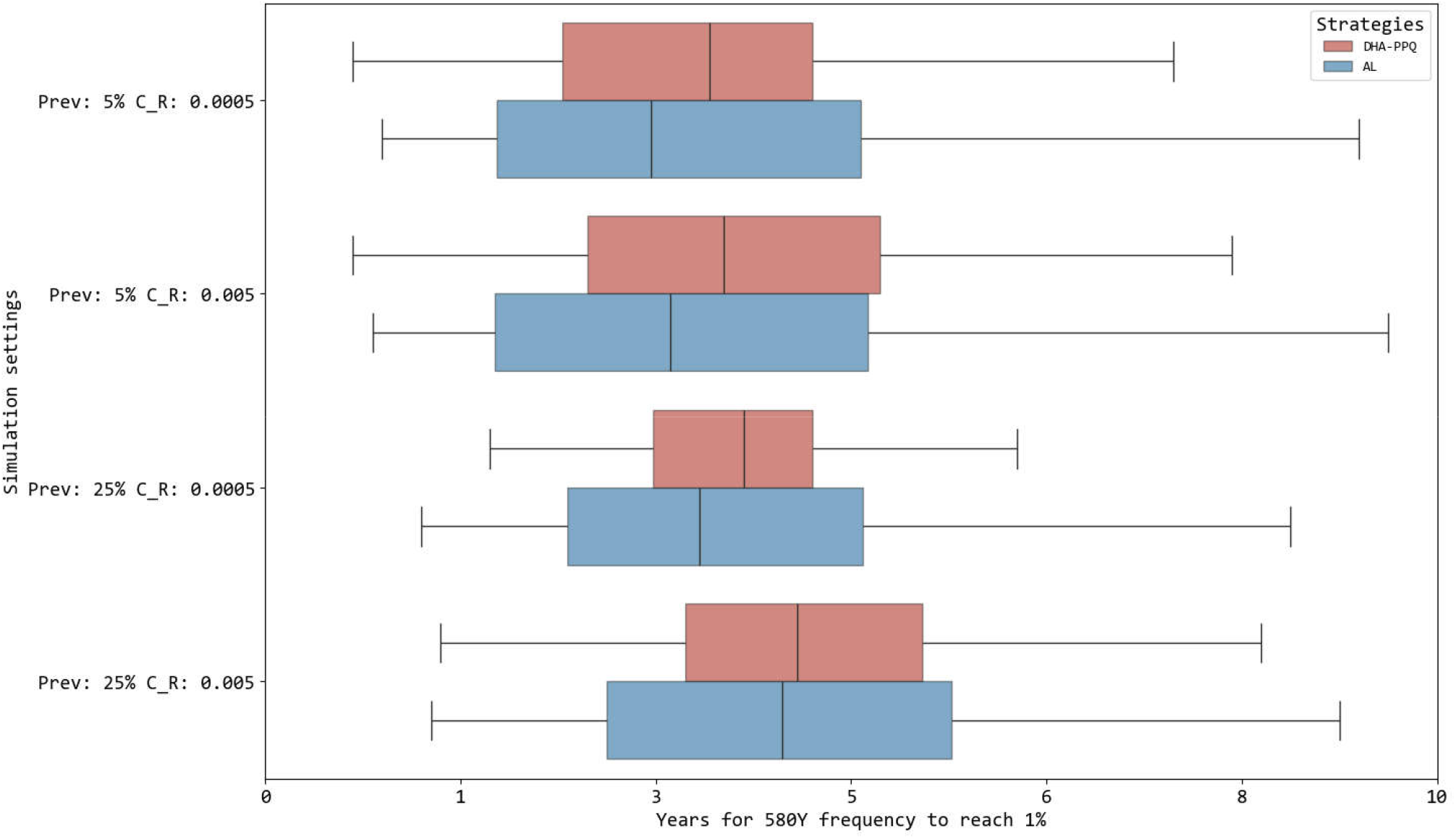
Time to 0.01 allele frequency of 580Y mutation in four epidemiological settings using single first-line therapy (red for DHA-PPQ and blue for AL). The *y*-axis shows the prevalence and *c*_*R*_ settings while the *x*-axis shows the number of years to reach 0.01 frequency.

**Figure S16a:**
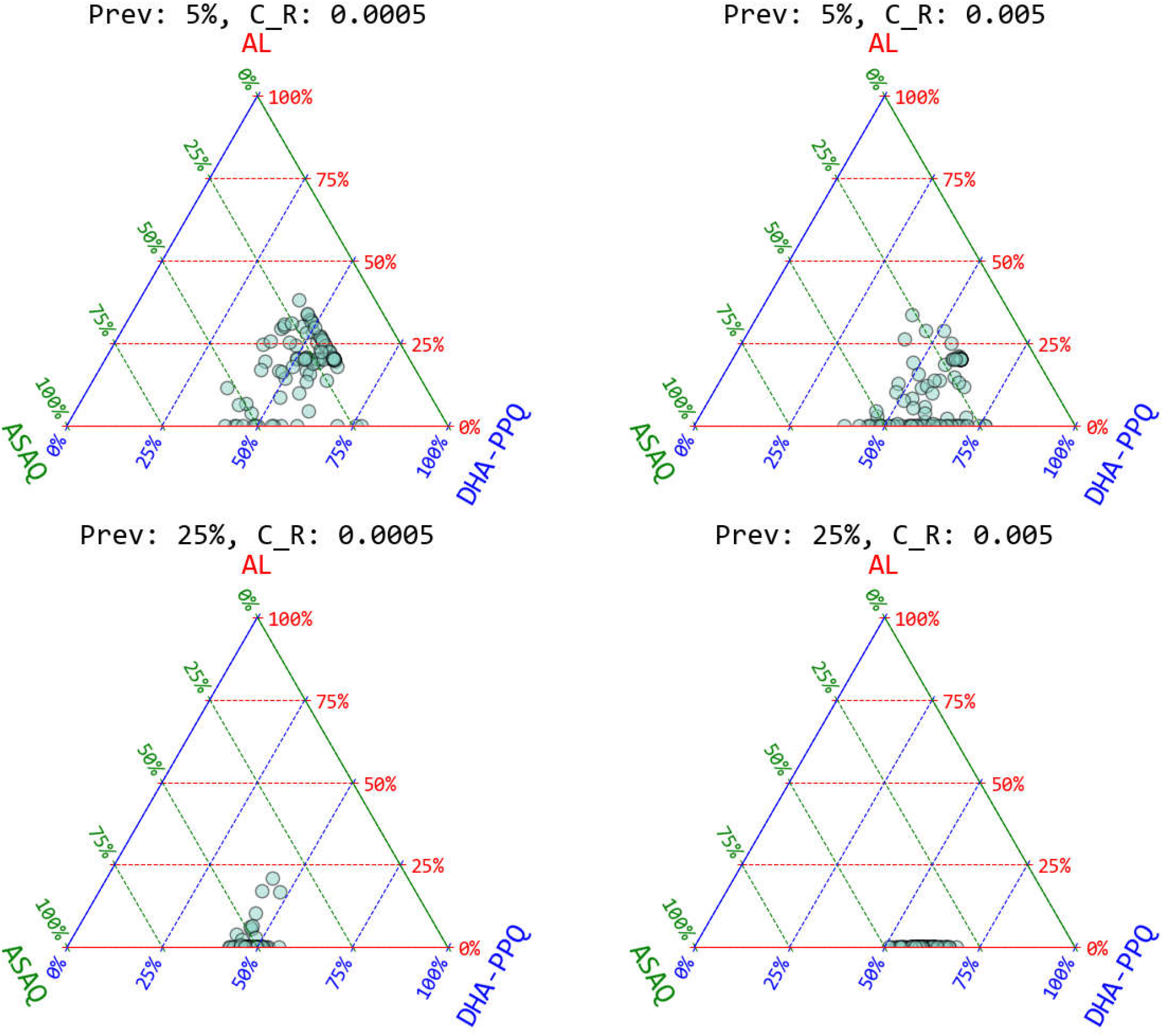
Barycentric plot of drug-use percentage in LMS adaptive cycling strategies over 15 years. Top, left, and right vertices correspond to the therapies used. Each dot corresponds to one simulation. Note that AL appears to be used for less time than the other therapies, as it is third in sequence in an LMS approach. At 25% prevalence and *c*_*R*_ = 0.005, DHA-PPQ is used for the majority of the 15-year period, after which there is a switch to ASAQ which is used through year 15 with no final switch to AL during this period.

